## Supporting Information for "Causal relationships between obesity and the leading causes of death in women and men"

### S1 Supporting Information

#### Contents

Abbreviations 2

Instruments 3

Instrument strength and selection 4

UK Biobank quality control 5

Analyses performed in British ancestry only subset 5

Mendelian randomization analyses using different weighting approaches 6

Pleiotropy-robust Mendelian randomization analyses 7

Analyses performed using the same number of cases and controls 7

Analyses performed using different diabetes diagnosis criteria 7

Table A. Characteristics of UK Biobank Participants included in the study. 9

Table B. Disease outcome definitions and diagnosis codes used to define cases and controls. 10

Table C. Instrument strength and estimates. 17

Table D. Linear regressions of the genetic risk scores with blood pressure traits. 19

Table E. Genetic risk score associations with smoking status. 20

Table F. Logistic regressions of the genetic risk scores with disease outcomes. 21

Table G. Mendelian randomization analyses with disease outcomes, unadjusted and adjusted for smoking status. 25

Table H. MR-Egger intercept test and MR-Egger estimates for the disease outcomes. 28

Table I. Logistic regressions of the genetic risk scores with disease outcomes using the same number of cases and controls for men and women compared to using all available cases and controls. 31

Table J. Mendelian randomization analyses for the associations between the obesity traits and having been or being a smoker. 36

Table K. Mendelian randomization analyses with blood pressure using different SNP-selection and weighting approaches, unadjusted for smoking. 38

Table L. Mendelian randomization analyses with blood pressure, unadjusted for smoking in British ancestry only subset. 41

Table M. Mendelian randomization analyses for blood pressure traits, unadjusted and adjusted for smoking status. 42

Table N. Two-sample Mendelian randomization analyses for fasting insulin and fasting glucose. 43

Figure A. SNP- and weight selection flowchart for all GRS construction approaches. 46

Figure B. Estimates and trait variance explained for the different sex-specific SNP weighting approaches using the same set of SNPs, separated by trait and sex. 48

Figure C. Genetic risk scores association with disease outcomes, stratified by sex. 49

Figure D. Effect of obesity risk factors on disease outcomes, stratified by sex, using GIANT 2015 sex-specific estimates as weights. 51

References 57

#### Abbreviations

BMI, Body mass index; CAD: coronary artery disease; CLD, chronic liver disease; COPD, chronic obstructive pulmonary disease; DBP, diastolic blood pressure; FG, fasting glucose; FI, fasting insulin; GIANT, Genetic Investigation of ANthropometric Traits; GRS, genetic risk score; GWAS, genome-wide association study; MR, Mendelian randomization; NCBI, the National Center for Biotechnology Information; NAFLD, non-alcoholic fatty liver disease; OR, odds ratio; T1D, type 1 diabetes; T2D, type 2 diabetes; SBP, systolic blood pressure; SD, standard deviation; SNP, single nucleotide polymorphism; WHO, the World Health Organization; WHR, waist-hip-ratio; WHRadjBMI, waist-hip-ratio adjusted for body mass index

#### Instruments

*Rationale*

In Mendelian randomization (MR), single nucleotide polymorphisms (SNPs) robustly associated with a trait are typically combined, weighted by their effect estimates, into a genetic risk score (GRS) (1). This GRS can then be used as an instrumental variable to investigate for causal associations between the trait and an outcome (1,2). A few studies have looked at sex-differences using genetic risk for obesity traits and MR methodology but have not thoroughly evaluated different SNP-selection and weighting approaches for sex-stratified analyses (3–5). As SNPs may have different effects in men and women (for example assessed by a $P_{het}$-value for sexual heterogeneity) (6), we constructed and compared six different approaches for constructing sex-specific GRSs for body mass index (BMI), waist-hip-ratio (WHR), and WHR adjusted for BMI (WHRadjBMI) (Fig A,B in S1 Supporting Information). In addition, we compared these six GRS construction approaches with a more common, but not sex-specific, approach to construct GRSs: using the same SNPs in men and women and weighting them by their combined-sexes effect estimates and applying them separately in each sex. In total, we considered five separate approaches to construction of the GRS;

**1. Sex-specific estimates approach:** Using the same SNPs for men and women, and weighting each SNP by its sex-specific effect estimate

**2. P-heterogeneity Bonferroni approach:** Using the same SNPs for men and women, and weighting each SNP either by its sex-specific or combined-sexes effect estimate, depending on if the SNPs $P_{het}$-value for sexual heterogeneity is below a Bonferroni threshold or not

**3-5. P-heterogeneity false discovery rate (FDR) approaches, using a 1%, 5% or 10% FDR threshold:** Using the same SNPs for men and women, and weighting each SNP either by its sex-specific or combined-sexes effect estimate, depending on if the SNP's $P_{het}$-value for sexual heterogeneity is below the corresponding FDR threshold or not

**6. Primary SNPs in each sex approach:** Using only the sex-specific primary SNPs in each sex, and weighting each SNP by its sex-specific effect estimate

**7. Combined-sexes approach:** Using the same SNPs for men and women and weighting each SNP by its combined-sexes effect estimate for both men and women. This approach was used for all combined-sexes analyses. We also used this approach separately in men and women for comparison with the sex-specific GRS construction approaches in the evaluation of the GRSs

These approaches were selected so as to only include SNPs robustly associated with the obesity trait under investigation, while also taking advantage of the increased sample size in the combined-sexes genome-wide association study (GWAS) and the possibly more precise estimates for SNPs without sexual heterogeneity. We therefore evaluated different $P_{het}$-value thresholds (including Bonferroni adjusted P-value and different FDR-value thresholds) for whether to weight SNPs by their sex-specific or combined-sexes weights in the P-heterogeneity Bonferroni and FDR approaches (methods 2-5 above).

*Selection of single nucleotide polymorphisms and weights for the instruments*

For the sex-specific estimates approach and the P-heterogeneity approaches, the primary ("index") associated SNPs per locus in any of the men, women, or combined-sexes European analyses from a recent GWAS with sex-specific summary statistics for each obesity trait were taken forward (available here: https://github.com/lindgrengroup/fatdistnGWAS) (6). The total sample sizes were up to 806,834 for BMI, 697,734 for WHR, and 694,649 for WHRadjBMI (6). In the original GWAS, primary genome-wide significant ($P<5\times10^{-9}$) SNPs were identified through proximal and joint conditional analysis using GCTA in associated ($P<5\times10^{-9}$) loci. The loci was defined by the inclusion of SNPs associated with the obesity trait P<0.05 within 5 Mb and in linkage disequilibrium (LD; $r^2>0.05$) with a top SNP (associated with the obesity trait $P<5\times10^{-9}$) (6). Overlapping loci were merged before GCTA analysis; for details see original study (6).

We then pruned these SNPs for independence, so that not correlated SNPs from different sex-specific analyses would be included, and to obtain a set of the same SNPs for all sex-strata. Independent SNPs were selected by taking all the SNPs specified as "index" SNPs in the men, women or combined-sexes analyses and then keeping the SNP with the lowest combined-sexes P-value within each 1 Mb sliding window for BMI, WHR, and WHRadjBMI separately. We later excluded SNPs based on long-distance LD ($r^2$>0.05), see section "*Exclusion of single nucleotide polymorphisms in the genetic risk scores*" below.

For the sex-specific estimates approach, European sex-specific estimates were used as weights. For the P-heterogeneity approaches, the $P_{het}$-values ("psexdiff") between men and women estimates from the original GWAS determined what weights were used, where it had been computed as;

$$t_{diff} = \frac{\beta_1 - \beta_2}{\sqrt{se1^2 + se2^2 - 2r \cdot se1 \cdot se2}}$$

where $\beta_1$ and $\beta_2$ corresponds to the estimates and $se_1$ and $se_2$ the standard errors for men and women respectively, and $r$ the Spearman rank correlation coefficient for the men and women SNP effects, and the $t_{diff}$-statistic is approximately normally distributed (6,7). For the P-heterogeneity Bonferroni approach, if $P_{het}$<0.05/number of SNPs included in the GRS for a genetic variant, it was weighted by the sex-specific European estimate, otherwise it was weighted by the estimate from the combined-sexes European meta-analysis. For the P-heterogeneity false discovery rate (FDR) approaches, sex-specific weights were used if the $P_{het}$-value was below the specified FDR threshold (1%, 5%, and 10% tested), otherwise the combined-sexes estimates were used as weights. The primary SNPs in each sex approach consisted of only including the specified "index" SNPs in each sex-specific European analysis for each obesity trait (6).

In the combined-sexes analyses, the same set of SNPs as in the sex-specific estimates approach were used; that is, all "index" SNPs in any of the men, women, or combined-sexes European analyses pruned for independence as described above (6). These were then weighted by their combined-sexes European estimates from the original GWAS (6). We also conducted sex-specific linear regression for these GRSs with their respective obesity trait for comparison.

*Exclusion of single nucleotide polymorphisms in the genetic risk scores*

For all GRSs, any SNPs found to be non-biallelic in the UK Biobank were excluded (n=6, rs7798002, rs73646205, rs10828247, rs10892873, rs699370, rs805768), as this would violate the additive assumption. Four SNPs were also excluded as they failed post-imputation quality control (rs11500477, rs144926207, rs547943994, rs9638713). To further ascertain that there was no LD between SNPs in a GRS, we computed the $r^2$ between all pairs of SNPs included in an instrument–for pairs of correlated ($r^2$>0.05) SNPs included in the same instrument, the SNP in each pair with the highest P-value was excluded (rs1025395, rs9302652, rs8189480).

#### Instrument strength and selection

Three assumptions must be fulfilled for an instrument variable to be valid, of which only the first one can be directly tested;

i. Association with the risk factor
ii. No association with any confounders of the risk factor-outcome relationship
iii. Only affects the outcome through the risk factor (i.e. no horizontal pleiotropy) (8,9)

We assessed the strengths of associations between the GRSs and their respective obesity trait by computing trait variance explained and the F-statistics. First, we standardized the obesity traits by rank inverse normal transformation of the residuals after regression of the obesity trait on baseline age, $age^2$, assessment centre, and if applicable sex, after any sample quality exclusions. This was done separately for men and women in the sex-specific analyses, but jointly in the combined analyses. For WHRadjBMI, we also adjusted for BMI. We thereafter performed linear regression, adjusting for array type and 10 principal components. Trait variance explained was defined as the increase in adjusted $R^2$ by inclusion of the GRS in the linear regression model. F-statistics were computed using the formula $F = \left(\frac{n-k-1}{k}\right)\left(\frac{R^2}{1-R^2}\right)$, where n is the number of observations, k=1 for number of instruments, and $R^2$ the unadjusted $R^2$ (10). The high F-statistics and high trait variance explained by the instruments indicated that the first instrument variable assumption was fulfilled (Table C in S1 Supporting Information) (11).

Both the combined-sexes weighted GRSs applied separately in each sex and most sex-specific approaches indicated significant heterogeneity (assessed by computing P-values from Cochran's Q test (12)) between the male and female estimates in their respective obesity trait, in particular for the waist-related GRSs. An evaluation of the different approaches indicated that the instruments improved (as judged by increased trait variance explained, increased F-statistics, and similarity of men and women SD-estimates in their respective obesity trait) the more sex-specific estimates were used as weights instead of combined-sexes estimates. This was particularly true for the waist-related traits and is illustrated in Fig B in S1 Supporting Information, were we compare the different weighting approaches using the same set of SNPs (also see Table C in S1 Supporting Information).

In summary, the sex-specific estimates GRS approach had the highest range of values of trait variance explained and F-statistics, and with no significant heterogeneity between men and women with their respective obesity association in SD-units in any trait. This implies that a 1-unit higher sex-specific GRS corresponded to similar increase in each obesity trait in men and women, facilitating the comparisons of disease associations between sexes. We therefore decided to use the sex-specific estimates approach as our main approach.

#### UK Biobank quality control

Individuals that had withdrawn consent, sex-chromosome aneuploidy, mismatch between self-reported and genetically inferred sex, reported incompatible ancestries in different assessments, more than 10 third degree relatives, that were heterozygosity or missingness outliers, that were not included in the autosome phasing or in the kinship calculations (as they were excluded because of for example extreme heterozygosity by the UK Biobank) were excluded (13). The remaining individuals were pruned to have no individuals related to a second degree or higher. The sample was then restricted to those with self-reported "White" ancestry. For all analyses involving anthropometric traits and blood pressure, women who were pregnant or unsure about being pregnant were excluded (N=279). For analyses involving anthropometric traits, individuals with BMI <15 kg/$m^2$ were excluded due to the potential for low BMI arising as a result of advanced disease (N=29).

We performed post-imputation quality control (6,18). All SNPs had an imputation info score >0.3. One SNP (rs547943994) failed our minor allele frequency threshold of >0.01%, one SNP failed our Hardy-Weinberg equilibrium exact test threshold of $P>1\times10^{-6}$ (rs9638713) and two SNPs failed our missing call rate threshold of <0.05 (rs144926207, rs11500477); these were excluded, as were non-biallelic SNPs (rs7798002, rs73646205, rs10828247, rs10892873, rs699370, and rs805768) (6,18). We also assessed the LD between all SNPs included in an instrument, using PLINK v1.90b3 (19) (--r2 inter-chr and --ld-window-r2 0). For the LD-computations, we excluded one sample in each pair related to a 3rd degree or higher (as computed by the UK biobank (14)) in our quality controlled samples, giving a final sample size of N=380,262. We then excluded the SNP with the highest P-value in each correlated ($r^2>0.05$) pair (rs1025395, rs9302652, and rs8189480). UK Biobank genotype quality control and analyses were performed in PLINK v1.90b3 (19).

#### Analyses performed in British ancestry only subset

Recent studies have highlighted potential population stratification in both Genetic Investigation of ANthropometric Traits (GIANT) and UKBB (20,21). The concern has been raised particularly for height, but there has also been suggestive evidence for BMI being affected, even if the bias is likely to be minor (20,21). Another study has indicated that GRS distributions can differ slightly between different European populations (22). We therefore conducted analyses both in all Europeans (self-report of any "white" category) in the main analyses and in a subset of British participants only (in the

"White British ancestry subset" as classified by the UK Biobank (14), defined as self-report as "British" and similar ancestry based on principal components analysis) as a sensitivity analysis.

The results were highly similar to those performed in all Europeans (Fig F and Tables J,L in S1 Supporting Information). All estimates in the disease outcome MR analyses remained robust, with the exception of the effect of WHR on COPD and renal failure in women (full European sample: $P=2.1\times10^{-4}$ for COPD and $P=2.6\times10^{-4}$ for renal failure; British ancestry only: P=0.002 for COPD and P=0.001 for renal failure, P-value threshold <0.001), and ischemic stroke in men (full European sample: $P=6.6\times10^{-5}$; British ancestry only: P=0.001). The $P_{het}$-values for sexual heterogeneity between the male and female estimates all still surpassed our multiple testing threshold, in spite of a loss of >10% of samples.

#### Mendelian randomization analyses using different weighting approaches

Care should be taken when conducting MR analyses using weights that have been derived in the same dataset as the MRs are performed in, as this can exacerbate weak instrument bias (1,23). To prevent weak instrument bias when a single dataset is used for both the risk-factor-instrument and outcome-instrument regressions (23), we took the following measures;

i. Only SNPs associated with their respective obesity traits at $P<5\times10^{-9}$ were included and combined into a single instrument for each trait (the GRSs)
ii. We quantified the F-statistic: the lowest F-statistic of any GRS taken forward was 4,921 (instruments with F-statistics <10 are usually denoted as weak) (11)

Taken together, this means that any potential weak instrument bias should be negligible (23). Despite this, we conducted MR analyses using additional weighting strategies (1,24) to ascertain robustness.

This included;

i. All sex-specific SNP- and weight-selection approaches that we had initially analyzed (Fig A in S1 Supporting Information), including the P-heterogeneity approaches and the primary SNPs associated in each sex only
ii. Use of unweighted allele scores; by constructing GRSs using the same set of SNPs as in the main analysis but giving each risk allele a weight of 1 when computing the GRSs
iii. Sex-specific weighting of the SNPs included in the main analysis by their GIANT 2015 (25,26) estimates, using the same approach as in the main analysis (after lift-over of the GIANT 2015 SNPs to dbSNP build 151). SNPs that were not present in GIANT were excluded (final number of SNPs: BMI, all sex-strata: 478 SNPs; WHR combined-sexes: 264; WHR women: 258; WHR men: 253; WHRadjBMI combined-sexes: 275; WHRadjBMI women: 263; WHRadjBMI men: 248) (25,26)

For the disease outcome MRs, the SNP- and weight selection approaches using different P-heterogeneity thresholds were highly similar as to the main approach using sex-specific estimates as weights. Whereas the BMI-lung cancer and WHRadjBMI-NAFLD associations in women did not surpass multiple testing correction for some of the weight selection approaches using different P-heterogeneity thresholds, the estimates were highly similar. The same Cochran's Q tests that had evidence against the null hypothesis in the main analysis had it in the all the P-heterogeneity weight-selection approaches.

For the primary SNPs in each sex only approach (comprising much fewer SNPs), unweighted, and GIANT-weighted approaches, all analyses significant in the main analysis (both estimates and Cochran's Q tests) were at least nominally (P<0.05) significant, except for the $P_{het}$-value between male and female estimates in the WHR-renal failure association (main analysis: $P_{het}=3.6\times10^{-4}$; primary SNPs in each sex only approach: $P_{het}=0.34$; unweighted approach: $P_{het}=2.4\times10^{-4}$; GIANT-weighted approach: $P_{het}=0.18$). Still, the male point estimate was higher than the female point estimate regardless of approach when assessing the WHR-renal failure association. In all, the sensitivity analyses supported the main findings. For the disease outcome MR results using the GIANT 2015 weighted GRSs and the unweighted GRSs, see Fig D,E in S1 Supporting Information.

For the obesity trait-risk factor MRs, the alternative weighting strategies supported the main findings. The P-heterogeneity approaches provided estimates similar to those from the sex-specific estimates approach. For results using the GIANT 2015 weighted GRSs and the unweighted GRSs, see Tables J,K in S1 Supporting Information.

#### Pleiotropy-robust Mendelian randomization analyses

If the genetic instruments are pleiotropic, i.e. affect the outcome through other pathways than the risk factor, this may bias results (27,28). The MR-Egger intercept test is a method to assess if the instruments show evidence of directionally biased pleiotropy, and the MR-Egger and weighted median-based MR methods can be used to obtain estimates more robust to pleiotropy, albeit with lower power to detect causal effects (28,29). As these methods require summary-level data, we derived SNP estimates for the disease outcomes using logistic regression and an additive model in PLINK (19). Adjustments were made for baseline age, sex, array, and 10 principal components. For a few SNPs we could not derive estimates for certain traits because of non-convergence in the logistic model; these were excluded (see Table H in S1 Supporting Information for number of SNPs included in each analysis). We thereafter used the package "MendelianRandomization" (30) to obtain results for the MR-Egger intercept test and estimates using the MR-Egger, IVW, and median-weighted methods for the main sex-specific estimates approach for the significant disease MR results. Cochran's Q test was used to assess sexual heterogeneity between the estimates (12).

A few of the GRSs used in the main analysis showed evidence of directional pleiotropy in the MR-Egger intercept test (P-value threshold set at <0.002 (=0.05/25) for 25 obesity trait-outcome combinations, Table H in S1 Supporting Information). This included the combined-sexes WHR GRS in CAD, the male BMI GRS in COPD and the male WHR GRS in renal failure, and the female WHR GRS in COPD and type 2 diabetes (T2D). The MR-Egger point estimates for all these associations were of lower magnitude than the point estimates from the main MR analyses. Despite this, for all the disease outcome MR analyses where one sex had significantly higher estimates than the other (women higher estimate: BMI-T2D; men higher estimates: WHR-COPD, WHR-renal failure, WHR and WHRadjBMI-chronic renal failure), the point estimates using the IVW, MR-Egger, and weighted-median methods were all higher in the same sex as in the main analysis. This indicates that the sex differences we see are robust using different MR methods to compute causal estimates.

#### Analyses performed using the same number of cases and controls

The number of cases differed substantially between the sexes for a number of outcomes, and for infertility we were unable to obtain reliable estimates in the men-only analyses because of the low number of cases (N=85). We therefore also performed all the sex-stratified GRSs in a random subsample in each sex category, using the same number of cases and controls in the sex-specific analyses and compared with the main analyses. This yielded similar results as to the main logistic regressions (Table I in S1 Supporting Information) and indicates that the observed sex differences are not due to power differences.

#### Analyses performed using different diabetes diagnosis criteria

For T1D and T2D, case status was decided using a validated algorithm from Eastwood *et al* for prevalent diabetes in the UKBB (31). The algorithm uses self-report data to classify all individuals in the UKBB into different categories. For both T1D and T2D, individuals could be assigned a possible or a probable case status, with greater evidence for the type-specific diabetes in probable cases. To ascertain robustness, we conducted the disease outcome MRs using probable cases only as a sensitivity analysis, which yielded highly similar results as in the main analysis.

Using probable cases only, a 1-SD higher BMI increased risk of T2D in all sex-strata (combined-sexes: OR 3.15, 95% CI 2.94-3.38, $P<1\times10^{-200}$; men: OR 2.77, 95% CI 2.54-3.02, $P=1.60\times10^{-117}$; women: OR 3.76 95% CI 3.35-4.22, $P=1.08\times10^{-111}$), with higher estimates in women than in men ($P_{het}=3.5\times10^{-5}$). A 1-SD higher WHR also increased risk of T2D in the combined sexes analysis (OR 3.66, 95% CI 3.34-4.02, $P=2.43\times10^{-167}$) as well as the men-only (OR 3.28, 95% CI 2.85-3.77, $P=4.31\times10^{-63}$) and women-only (OR 3.75, 95% CI 3.32-4.24, $P=1.02\times10^{-101}$) analyses, but with no difference between the sexes ($P_{het}$=0.1). WHRadjBMI also increased risk of T2D (combined sexes: OR

2.11, 95% CI 1.95-2.28, $P=3.82\times10^{-79}$; men: OR 1.82, 95% CI 1.61-2.07, $P=3.57\times10^{-21}$; women: OR 2.19, 95% CI 1.97-2.42, $P=1.85\times10^{-51}$, per 1-SD higher WHRadjBMI), and similar to the main analysis there was no sexual heterogeneity in the estimates ($P_{het}$=0.03).

Results were also very similar using a stricter T1D case definition as to the main results. A 1-SD higher BMI significantly increased risk of T1D with OR 1.61 (95% CI 1.30-2.01, $P=1.81\times10^{-5}$) in the combined sexes analysis. In men, a 1-SD higher BMI was not significantly associated with increased risk of T1D (OR 1.41; 95% CI 1.05-1.88, P=0.02), similar to the main analysis. In women, T1D risk increased with OR 1.87 (95% CI 1.35-2.59, $P=1.55\times10^{-4}$) per 1-SD higher BMI. There was no difference between the sexes ($P_{het}$=0.2). In all, the same obesity trait-disease outcome combinations and tests for sexual heterogeneity were significant using the stricter diabetes case definitions as in the main analysis.

**Table A. Characteristics of UK Biobank Participants included in the study.**

| Characteristic | Men | Women |
|---|---|---|
| Individuals, N (%) | 195,041 (46.1) | 228,466 (53.9) |
| British, N (%) | 174,261 (89.3) | 201,958 (88.4) |
| Age, mean (SD), years | 57.0 (8.1) | 56.6 (7.9) |
| UK BiLEVE array, N (%) | 23,216 (11.9) | 22,848 (10.0) |
| Body mass index, mean (SD), kg/m2 | 27.9 (4.2) | 27.0 (5.1) |
| Waist circumference, mean (SD), cm | 97.1 (11.3) | 84.5 (12.5) |
| Hip circumference, mean (SD), cm | 103.5 (7.6) | 103.3 (10.3) |
| Waist-hip-ratio, mean (SD) | 0.94 (0.07) | 0.82 (0.07) |
| Systolic blood pressure, mean (SD), mmHg | 144.8 (19.4) | 137.9 (21.2) |
| Diastolic blood pressure, mean (SD), mmHg | 86.6 (11.0) | 82.3 (11.1) |
| Type 2 diabetes cases, N (%) | 11,785 (6.0) | 6,544 (2.9) |
| Coronary artery disease cases, N (%) | 24,444 (12.5) | 11,578 (5.1) |
| Breast cancer cases, N (%) | - | 14,325 (6.3) |
| Chronic liver disease cases, N (%) | 824 (0.4) | 543 (0.2) |
| Colorectal cancer cases, N (%) | 3,150 (1.6) | 2,380 (1.0) |
| COPD cases, N (%) | 7,874 (4.0) | 6,799 (3.0) |
| Dementia cases, N (%) | 583 (0.3) | 447 (0.2) |
| Infertility cases, N (%) | 85 (0.0) | 1,593 (0.7) |
| Lung cancer cases, N (%) | 1,477 (0.8) | 1,239 (0.5) |
| NAFLD cases, N (%) | 909 (0.5) | 775 (0.3) |
| Renal failure cases, N (%) | 5,697 (2.9) | 3,909 (1.7) |
| Renal failure, acute, cases, N (%) | 3,035 (1.6) | 1,651 (0.7) |
| Renal failure, chronic, cases, N (%) | 2,581 (1.3) | 2,018 (0.9) |
| Stroke cases, N (%) | 6,334 (3.2) | 4,454 (1.9) |
| Stroke, hemorrhagic, cases, N (%) | 928 (0.5) | 979 (0.4) |
| Stroke, ischemic, cases, N (%) | 2,171 (1.1) | 1,180 (0.5) |
| Type 1 diabetes cases, N (%) | 826 (0.4) | 677 (0.3) |

COPD, chronic obstructive pulmonary disease; NAFLD, non-alcoholic fatty liver disease; SD, standard deviation.

[a]Participants were denoted as "British" if they were in the British ancestry subset as defined by the UK Biobank (23) (based on self-report of British ancestry and similar ancestry according to principal components analysis)

[b]UK BiLEVE array is the number of participants genotyped on that array as opposed to the UK Biobank Axiom array

**Table B. Disease outcome definitions and diagnosis codes used to define cases and controls.**

| Outcome | ICD-10[a] | ICD-9[b] | OPCS-4[c] | NI, Non-cancer[d] | NI, Cancer[e] | NI, operation[f] | Self-reported Diagnosis by Doctor[g] | Description |
|---|---|---|---|---|---|---|---|---|
| Breast cancer | C50; C500; C501; C502; C503; C504; C505; C506; C508; C509; Z853 | 174; 1740; 1741; 1742; 1743; 1744; 1745; 1746; 1748; 1749; V103 | | | 1002 | | | Codes for breast cancer, including personal history codes. Run in women only |
| CAD | I20; I200; I201; I208; I209; I21; I210; I211; I212; I213; I214; I219; I21X; I22; I220; I221; I228; I229; I23; I230; I231; I232; I233; I234; I235; I236; I238; I24; I240; I241; I248; I249; I251; I252; I255; I256; I258; I259 | 410; 4109; 411; 4119; 412; 4129; 413; 4139; 4140; 4148; 4149 | K40; K401; K402; K403; K404; K408; K409; K41; K411; K412; K413; K414; K418; K419; K42; K421; K422; K423; K424; K428; K429; K43; K431; K432; K433; K434; K438; K439; K44; K441; K442; K448; K449; K45; K451; K452; K453; K454; K455; K456; K458; K459; K46; K461; K462; K463; K464; K465; K468; K469; K49; K491; K492; K493; K494; K498; K499; K501; K75; K751; K752; K753; K754; K758; K759 | 1074; 1075 | | 1070; 1095; 1523 | 1; 2 | Codes for myocardial infarction, percutaneous transluminal coronary angioplasty, coronary artery bypass grafting, chronic ischemic heart disease and angina. Outcome definition from Nelson et al (32); SOFT CAD definition including angina. |

| **Outcome** | **ICD-10**[a] | **ICD-9**[b] | **OPCS-4**[c] | **NI, Non-cancer**[d] | **NI, Cancer**[e] | **NI, operation**[f] | **Self-reported Diagnosis by Doctor**[g] | **Description** |
|---|---|---|---|---|---|---|---|---|
| CAD - control group exclusions | I250; I253; I254 | 4141 | | | | | | Participants were excluded from the CAD-control group if they had these codes pertaining to heart aneurysm and atherosclerotic cardiovascular disease. Outcome definition from Nelson et al (32); SOFT CAD definition including angina. |
| COPD | J41; J410; J411; J418; J42; J43; J431; J432; J438; J439; J44; J440; J441; J448; J449 | 491; 4910; 4911; 4912; 4918; 4919; 492; 4929 | | 1112; 1113; 1472 | | | 6 | Codes for chronic bronchitis, emphysema, COPD, or complications specified from COPD |
| Chronic liver disease | K702; K703; K704; K717; K721; K74; K740; K741; K742; K743; K744; K745; K746 | 27103; 4562; 571; 5712; 5715; 57150; 57151; 57158; 57159; 5716 | | 1604; 1158 | | | | Codes for fibrosis, sclerosis, cirrhosis of liver and liver failure, including if caused by alcohol, toxic liver disease, biliary cirrhosis, and other causes as well as codes defining complications specified as caused by these |
| Colorectal Cancer | C18; C180; C181; C182; C183; C184; C185; C186; C187; C188; C189; C19; C20 | 153; 1530; 1531; 1532; 1533; 1534; 1535; 1536; 1537; 1538; 1539; 154; 1540; 1541 | | | 1020; 1022; 1023 | | | Codes for cancers from caecum to rectum, including appendix |

| Outcome | ICD-10[a] | ICD-9[b] | OPCS-4[c] | NI, Non-cancer[d] | NI, Cancer[e] | NI, operation[f] | Self-reported Diagnosis by Doctor[g] | Description |
|---|---|---|---|---|---|---|---|---|
| Dementia | F00; F000; F001; F002; F009; F01; F010; F011; F012; F013; F018; F019; F03; G30; G300; G301; G308; G309 | 2900; 2901; 2902; 2903; 2904; 3310 | | 1263 | | | | Codes for dementia in Alzheimer's, vascular dementia, unspecified dementia, senile and presenile dementia |
| Infertility | N97; N970; N971; N972; N973; N978; N979; N46 | 628; 6280; 6281; 6282; 6283; 6284; 6288; 6289; 606; 6069 | | 1403; 1404 | | | | Codes for male or female infertility of different anatomical origins. Sex-specific codes applied in relevant sex only |
| Lung cancer | C33; C34; C340; C341; C342; C343; C348; C349; Z851 | 162; 1620; 1622; 1623; 1624; 1625; 1628; 1629; V101 | | | 1001; 1027; 1028; 1080 | | | Codes for cancers in trachea, bronchi, and lungs, including personal history codes |
| NAFLD | K760 | | | | | | | Code for fatty liver disease |
| Renal Failure | N17; N170; N171; N172; N178; N179; N18; N180; N181; N182; N183; N184; N185; N188; N189; N19; I120; I131; I132; Z992 | 584; 5845; 5846; 5847; 5848; 5849; 585; 5859; 586; 5869 | L746; X40; X401; X402; X403; X404; X405; X406; X407; X408; X409; X41; X411; X412; X418; X419; X42; X421; X428; X429 | 1192; 1193; 1194 | | 1476; 1580; 1581; 1582 | | Codes for both acute, chronic and unspecified renal failure and chronic kidney disease. Also includes renal failure from hypertensive disease and various dialysis procedures in ICD-10 and OPCS-4 codes |
| Renal Failure - acute | N17; N170; N171; N172; N178; N179 | 584; 5845; 5846; 5847; 5848; 5849 | | | | | | Codes that are specifically for acute renal failure |

| Outcome | ICD-10[a] | ICD-9[b] | OPCS-4[c] | NI, Non-cancer[d] | NI, Cancer[e] | NI, operation[f] | Self-reported Diagnosis by Doctor[g] | Description |
|---|---|---|---|---|---|---|---|---|
| Renal Failure - chronic | N18; N180; N181; N182; N183; N184; N185; N188; N189 | 585; 5859 | | | | | | Codes that are specifically for chronic kidney disease |
| Renal Failure - control group exclusions | N17; N170; N171; N172; N178; N179; N18; N180; N181; N182; N183; N184; N185; N188; N189; N19; I120; I131; I132; Z992 | 584; 5845; 5846; 5847; 5848; 5849; 585; 5859; 586; 5869 | L746; X40; X401; X402; X403; X404; X405; X406; X407; X408; X409; X41; X411; X412; X418; X419; X42; X421; X428; X429 | 1192; 1193; 1194 | | 1476; 1580; 1581; 1582 | | Participants were excluded from the renal failure control groups, including acute renal failure and chronic kidney disease, if they had these codes pertaining to renal failure |
| Stroke | I60; I600; I601; I602; I603; I604; I605; I606; I607; I608; I609; I61; I610; I611; I612; I613; I614; I615; I616; I618; I619; I63; I630; I631; I632; I633; I634; I635; I636; I638; I639; I64 | 430; 4309; 431; 4319; 434; 4340; 4341; 4349; 436; 4369 | | 1081; 1086; 1491; 1583 | | | 3 | Codes for subarachnoid and intracerebral hemorrhages and cerebral infarctions including cerebral thrombosis and embolism, and unspecified stroke. Does not included transient cerebral ischaemia but includes acute but ill-defined cerebrovascular disease |

| **Outcome** | **ICD-10**[a] | **ICD-9**[b] | **OPCS-4**[c] | **NI, Non-cancer**[d] | **NI, Cancer**[e] | **NI, operation**[f] | **Self-reported Diagnosis by Doctor**[g] | **Description** |
|---|---|---|---|---|---|---|---|---|
| Stroke - control group exclusions | I60; I600; I601; I602; I603; I604; I605; I606; I607; I608; I609; I61; I610; I611; I612; I613; I614; I615; I616; I618; I619; I63; I630; I631; I632; I633; I634; I635; I636; I638; I639; I64; G45; G450; G451; G452; G453; G454; G458; G459 | 430; 4309; 431; 4319; 434; 4340; 4341; 4349; 436; 4369; 435; 4359 | | 1081; 1082; 1086; 1491; 1583 | | | 3 | Participants were excluded from the stroke control groups, including hemorrhagic and ischemic stroke, if they had these codes pertaining to stroke and transient ischemic attacks |
| Stroke - hemorrhagic | I60; I600; I601; I602; I603; I604; I605; I606; I607; I608; I609; I61; I610; I611; I612; I613; I614; I615; I616; I618; I619 | 430; 4309; 431; 4319 | | 1086; 1491 | | | | Codes that denote hemorrhagic stroke |
| Stroke - ischemic | I63; I630; I631; I632; I633; I634; I635; I636; I638; I639 | 434; 4340; 4341; 4349 | | 1583 | | | | Codes that denote ischemic stroke |

| Outcome | ICD-10[a] | ICD-9[b] | OPCS-4[c] | NI, Non-cancer[d] | NI, Cancer[e] | NI, operation[f] | Self-reported Diagnosis by Doctor[g] | Description |
|---|---|---|---|---|---|---|---|---|
| Type 1 diabetes | | | | | | | | Algorithm sorts participants to likely diabetes status by using information on e.g. self-reported diabetes diagnosis, age of diagnosis, medications, start of insulin within a year of diagnosis, and other self-reported data at the baseline visit. Outcome definition from Eastwood et al (31) algorithm. Probable and possible cases or probable only. Controls defined as diabetes unlikely. |

| Outcome | ICD-10[a] | ICD-9[b] | OPCS-4[c] | NI, Non-cancer[d] | NI, Cancer[e] | NI, operation[f] | Self-reported Diagnosis by Doctor[g] | Description |
|---|---|---|---|---|---|---|---|---|
| Type 2 diabetes | | | | | | | | Algorithm sorts participants to likely diabetes status by using information on e.g. self-reported diabetes diagnosis, age of diagnosis, medications, start of insulin within a year of diagnosis, and other self-reported data at the baseline visit. Outcome definition from Eastwood et al (31) algorithm. Probable and possible cases or probable only. Controls defined as diabetes unlikely. |

CAD, coronary artery disease; COPD, chronic obstructive pulmonary disease; NAFLD, non-alcoholic fatty liver disease; NI, nurse interview.
The following data fields were used for each diagnosis code type;
[a]ICD-10 codes: 41202 (main diagnosis), 41204 (secondary diagnoses), 40006 (cancer register), 40001 (primary cause of death), 40002 (secondary causes of death)
[b]ICD-9: 41203 (main diagnosis), 41205 (secondary diagnoses), 40013 (cancer register)
[c]OPCS-4 (operative procedures): 41200 (main code), 41210 (secondary code)
[d]Self-reported at nurse's interview, non-cancer codes: 20002
[e]Self-reported at nurse's interview, cancer codes: 20001
[f]Self-reported at nurse's interview, operation codes: 20004
[g]Self-reported diagnosis from doctor: 6150 (for stroke and CAD) and 6152 (for COPD)

**Table C. Instrument strength and estimates.**

| GRS | N SNPs | Sex-Strata | Trait | $P^{ab}$ | $F^{a}$ | $R^{2a}$ | Estimate (95% CI)[a] | $P_{het}^{ac}$ |
|---|---|---|---|---|---|---|---|---|
| **BMI, combined estimates** | **565** | **Combined** | **BMI** | **$<1\times10^{-200}$** | **26,466** | **5.90%** | **1.01 (1.00,1.02)** | **-** |
| BMI, combined estimates | 565 | Men | BMI | $<1\times10^{-200}$ | 12,466 | 6.03% | 1.02 (1.00,1.04) | 0.34 |
| **BMI, male sex-specific estimates** | **565** | **Men** | **BMI** | **$<1\times10^{-200}$** | **12,826** | **6.19%** | **1.02 (1.00,1.04)** | **0.04** |
| BMI, male P-heterogeneity Bonferroni | 565 | Men | BMI | $<1\times10^{-200}$ | 12,490 | 6.04% | 1.02 (1.00,1.04) | 0.25 |
| BMI, male P-heterogeneity FDR 1% | 565 | Men | BMI | $<1\times10^{-200}$ | 12,490 | 6.04% | 1.02 (1.00,1.04) | 0.25 |
| BMI, male P-heterogeneity FDR 5% | 565 | Men | BMI | $<1\times10^{-200}$ | 12,539 | 6.06% | 1.02 (1.00,1.03) | 0.67 |
| BMI, male P-heterogeneity FDR 10% | 565 | Men | BMI | $<1\times10^{-200}$ | 12,561 | 6.07% | 1.02 (1.00,1.03) | 0.77 |
| BMI, primary SNPs in males | 220 | Men | BMI | $<1\times10^{-200}$ | 9,057 | 4.45% | 1.01 (0.99,1.03) | 0.05 |
| BMI, combined estimates | 565 | Women | BMI | $<1\times10^{-200}$ | 14,160 | 5.86% | 1.01 (0.99,1.03) | 0.34 |
| **BMI, female sex-specific estimates** | **565** | **Women** | **BMI** | **$<1\times10^{-200}$** | **14,406** | **5.96%** | **1.00 (0.98,1.01)** | **0.04** |
| BMI, female P-heterogeneity Bonferroni | 565 | Women | BMI | $<1\times10^{-200}$ | 14,174 | 5.87% | 1.01 (0.99,1.02) | 0.25 |
| BMI, female P-heterogeneity FDR 1% | 565 | Women | BMI | $<1\times10^{-200}$ | 14,174 | 5.87% | 1.01 (0.99,1.02) | 0.25 |
| BMI, female P-heterogeneity FDR 5% | 565 | Women | BMI | $<1\times10^{-200}$ | 14,211 | 5.88% | 1.01 (1.00,1.03) | 0.67 |
| BMI, female P-heterogeneity FDR 10% | 565 | Women | BMI | $<1\times10^{-200}$ | 14,217 | 5.88% | 1.01 (1.00,1.03) | 0.77 |
| BMI, primary SNPs in females | 278 | Women | BMI | $<1\times10^{-200}$ | 10,265 | 4.32% | 0.98 (0.97,1.00) | 0.05 |
| **WHR, combined estimates** | **324** | **Combined** | **WHR** | **$<1\times10^{-200}$** | **14,932** | **3.41%** | **1.03 (1.02,1.05)** | **-** |
| WHR, combined estimates | 324 | Men | WHR | $<1\times10^{-200}$ | 3,873 | 1.95% | 0.78 (0.76,0.80) | $1.24\times10^{-157}$ |
| **WHR, male sex-specific estimates** | **324** | **Men** | **WHR** | **$<1\times10^{-200}$** | **4,921** | **2.47%** | **1.01 (0.98,1.04)** | **0.04** |
| WHR, male P-heterogeneity Bonferroni | 324 | Men | WHR | $<1\times10^{-200}$ | 4,636 | 2.33% | 0.93 (0.90,0.96) | $2.48\times10^{-20}$ |
| WHR, male P-heterogeneity FDR 1% | 324 | Men | WHR | $<1\times10^{-200}$ | 4,758 | 2.39% | 0.96 (0.93,0.98) | $4.74\times10^{-11}$ |
| WHR, male P-heterogeneity FDR 5% | 324 | Men | WHR | $<1\times10^{-200}$ | 4,819 | 2.42% | 0.97 (0.94,1.00) | $1.01\times10^{-07}$ |
| WHR, male P-heterogeneity FDR 10% | 324 | Men | WHR | $<1\times10^{-200}$ | 4,838 | 2.42% | 0.98 (0.95,1.01) | $8.55\times10^{-06}$ |
| WHR, primary SNPs in males | 77 | Men | WHR | $<1\times10^{-200}$ | 2,842 | 1.44% | 0.99 (0.95,1.02) | 0.02 |
| WHR, combined estimates | 324 | Women | WHR | $<1\times10^{-200}$ | 11,690 | 4.88% | 1.23 (1.21,1.26) | $1.24\times10^{-157}$ |
| **WHR, female sex-specific estimates** | **324** | **Women** | **WHR** | **$<1\times10^{-200}$** | **12,661** | **5.27%** | **1.04 (1.03,1.06)** | **0.04** |
| WHR, female P-heterogeneity Bonferroni | 324 | Women | WHR | $<1\times10^{-200}$ | 12,443 | 5.18% | 1.09 (1.07,1.10) | $2.48\times10^{-20}$ |

| GRS | N SNPs | Sex-Strata | Trait | $P^{ab}$ | $F^a$ | $R^{2a}$ | Estimate (95% CI)[a] | $P_{het}^{ac}$ |
|---|---|---|---|---|---|---|---|---|
| WHR, female P-heterogeneity FDR 1% | 324 | Women | WHR | $<1\times10^{-200}$ | 12,542 | 5.22% | 1.07 (1.05,1.09) | $4.74\times10^{-11}$ |
| WHR, female P-heterogeneity FDR 5% | 324 | Women | WHR | $<1\times10^{-200}$ | 12,593 | 5.24% | 1.06 (1.04,1.08) | $1.01\times10^{-07}$ |
| WHR, female P-heterogeneity FDR 10% | 324 | Women | WHR | $<1\times10^{-200}$ | 12,604 | 5.24% | 1.06 (1.04,1.08) | $8.55\times10^{-06}$ |
| WHR, primary SNPs in females | 203 | Women | WHR | $<1\times10^{-200}$ | 11,255 | 4.71% | 1.04 (1.02,1.06) | 0.02 |
| **WHRadjBMI, combined estimates** | **337** | **Combined** | **WHRadjBMI** | **$<1\times10^{-200}$** | **19,516** | **4.42%** | **1.03 (1.02,1.05)** | **-** |
| WHRadjBMI, combined estimates | 337 | Men | WHRadjBMI | $<1\times10^{-200}$ | 4,363 | 2.20% | 0.73 (0.71,0.75) | $<1\times10^{-200}$ |
| **WHRadjBMI, male sex-specific estimates** | **337** | **Men** | **WHRadjBMI** | **$<1\times10^{-200}$** | **5,548** | **2.78%** | **1.02 (1.00,1.05)** | **0.24** |
| WHRadjBMI, male P-heterogeneity Bonferroni | 337 | Men | WHRadjBMI | $<1\times10^{-200}$ | 5,132 | 2.57% | 0.91 (0.88,0.93) | $1.04\times10^{-33}$ |
| WHRadjBMI, male P-heterogeneity FDR 1% | 337 | Men | WHRadjBMI | $<1\times10^{-200}$ | 5,311 | 2.66% | 0.94 (0.92,0.97) | $7.88\times10^{-18}$ |
| WHRadjBMI, male P-heterogeneity FDR 5% | 337 | Men | WHRadjBMI | $<1\times10^{-200}$ | 5,407 | 2.71% | 0.97 (0.95,1.00) | $1.23\times10^{-08}$ |
| WHRadjBMI, male P-heterogeneity FDR 10% | 337 | Men | WHRadjBMI | $<1\times10^{-200}$ | 5,445 | 2.73% | 0.99 (0.97,1.02) | $1.59\times10^{-04}$ |
| WHRadjBMI, primary SNPs in males | 90 | Men | WHRadjBMI | $<1\times10^{-200}$ | 3,850 | 1.94% | 1.02 (0.98,1.05) | 0.30 |
| WHRadjBMI, combined estimates | 337 | Women | WHRadjBMI | $<1\times10^{-200}$ | 16,431 | 6.74% | 1.27 (1.26,1.29) | $<1\times10^{-200}$ |
| **WHRadjBMI, female sex-specific estimates** | **337** | **Women** | **WHRadjBMI** | **$<1\times10^{-200}$** | **17,263** | **7.06%** | **1.04 (1.03,1.06)** | **0.24** |
| WHRadjBMI, female P-heterogeneity Bonferroni | 337 | Women | WHRadjBMI | $<1\times10^{-200}$ | 17,026 | 6.97% | 1.09 (1.08,1.11) | $1.04\times10^{-33}$ |
| WHRadjBMI, female P-heterogeneity FDR 1% | 337 | Women | WHRadjBMI | $<1\times10^{-200}$ | 17,128 | 7.00% | 1.08 (1.06,1.09) | $7.88\times10^{-18}$ |
| WHRadjBMI, female P-heterogeneity FDR 5% | 337 | Women | WHRadjBMI | $<1\times10^{-200}$ | 17,179 | 7.02% | 1.06 (1.05,1.08) | $1.23\times10^{-08}$ |
| WHRadjBMI, female P-heterogeneity FDR 10% | 337 | Women | WHRadjBMI | $<1\times10^{-200}$ | 17,200 | 7.03% | 1.05 (1.04,1.07) | $1.59\times10^{-04}$ |
| WHRadjBMI, primary SNPs in females | 264 | Women | WHRadjBMI | $<1\times10^{-200}$ | 17,220 | 7.04% | 1.03 (1.02,1.05) | 0.30 |

BMI, body mass index; F, F-statistic; FDR, false discovery rate; GRS, genetic risk score; N, Number of; P, P-value; $R^2$, trait variance explained; WHR, waist-hip-ratio; WHRadjBMI, waist-hip-ratio adjusted for body mass index.

The sex-specific estimates approach used as the main approach in bold.

[a]The estimates, P-values, F-statistics, and trait variance explained for each GRS in their respective trait per 1-unit higher GRS, corresponding to a predicted 1-SD higher obesity trait

[b]P-value threshold set at <0.001 (=0.05/45) for 45 instruments assessed

[c]$P_{het}$ from Cochran's Q test for comparisons between the male and female estimates, either between the combined-sexes estimates approach applied in each sex and obesity trait separately, or between similar sex-specific approaches for each obesity trait. $P_{het}$-threshold set at <0.002 (=0.05/21) for 21 male-female estimate comparisons

**Table D. Linear regressions of the genetic risk scores with blood pressure traits.**

| GRS | Outcome | N SNPs | Sex-Strata | Estimate, Clin (95% CI)[a] | P, Clin[b] | $P_{het}$, Clin[c] | Estimate, SD (95% CI)[a] | P, SD[b] | $P_{het}$, SD[c] |
|---|---|---|---|---|---|---|---|---|---|
| BMI, combined | SBP | 565 | Combined | 3.57 (3.32,3.81) | $3.86\times10^{-181}$ | - | 0.19 (0.18,0.21) | $3.22\times10^{-190}$ | - |
| BMI, men | SBP | 565 | Men | 3.48 (3.14,3.82) | $3.78\times10^{-88}$ | 0.64 | 0.20 (0.18,0.21) | $3.50\times10^{-93}$ | 0.703 |
| BMI, women | SBP | 565 | Women | 3.59 (3.26,3.92) | $5.77\times10^{-100}$ | | 0.19 (0.17,0.21) | $4.89\times10^{-104}$ | |
| BMI, combined | DBP | 565 | Combined | 2.96 (2.82,3.10) | $<1\times10^{-200}$ | - | 0.27 (0.26,0.29) | $<1\times10^{-200}$ | - |
| BMI, men | DBP | 565 | Men | 2.65 (2.45,2.86) | $3.71\times10^{-143}$ | $9.51\times10^{-05}$ | 0.25 (0.23,0.27) | $1.83\times10^{-147}$ | $2.42\times10^{-04}$ |
| BMI, women | DBP | 565 | Women | 3.20 (3.02,3.39) | $<1\times10^{-200}$ | | 0.29 (0.28,0.31) | $<1\times10^{-200}$ | |
| WHR, combined | SBP | 324 | Combined | 4.11 (3.78,4.44) | $2.19\times10^{-134}$ | - | 0.22 (0.21,0.24) | $7.55\times10^{-140}$ | - |
| WHR, men | SBP | 324 | Men | 3.65 (3.11,4.18) | $1.81\times10^{-40}$ | 0.76 | 0.20 (0.17,0.23) | $5.08\times10^{-42}$ | 0.824 |
| WHR, women | SBP | 324 | Women | 3.75 (3.38,4.12) | $7.58\times10^{-88}$ | | 0.20 (0.18,0.22) | $9.31\times10^{-92}$ | |
| WHR, combined | DBP | 324 | Combined | 2.76 (2.57,2.95) | $9.53\times10^{-181}$ | - | 0.26 (0.24,0.27) | $2.27\times10^{-183}$ | - |
| WHR, men | DBP | 324 | Men | 2.82 (2.50,3.14) | $4.39\times10^{-67}$ | 0.034 | 0.26 (0.23,0.29) | $1.86\times10^{-68}$ | 0.029 |
| WHR, women | DBP | 324 | Women | 2.41 (2.20,2.62) | $3.25\times10^{-112}$ | | 0.22 (0.20,0.24) | $5.53\times10^{-114}$ | |
| WHRadjBMI, combined | SBP | 337 | Combined | 2.35 (2.06,2.63) | $1.49\times10^{-57}$ | - | 0.13 (0.11,0.14) | $5.02\times10^{-60}$ | - |
| WHRadjBMI, men | SBP | 337 | Men | 1.61 (1.10,2.13) | $7.13\times10^{-10}$ | 0.033 | 0.09 (0.06,0.12) | $3.66\times10^{-10}$ | 0.058 |
| WHRadjBMI, women | SBP | 337 | Women | 2.27 (1.95,2.59) | $2.42\times10^{-44}$ | | 0.12 (0.10,0.14) | $1.08\times10^{-46}$ | |
| WHRadjBMI, combined | DBP | 337 | Combined | 1.11 (0.94,1.27) | $9.43\times10^{-39}$ | - | 0.10 (0.09,0.12) | $1.57\times10^{-39}$ | - |
| WHRadjBMI, men | DBP | 337 | Men | 0.98 (0.68,1.29) | $2.95\times10^{-10}$ | 0.674 | 0.09 (0.06,0.12) | $2.10\times10^{-10}$ | 0.667 |
| WHRadjBMI, women | DBP | 337 | Women | 1.06 (0.88,1.24) | $1.48\times10^{-30}$ | | 0.10 (0.08,0.11) | $3.91\times10^{-31}$ | |

BMI, body mass index; CI, confidence interval; Clin, clinical; DBP, diastolic blood pressure; GRS, genetic risk scores; N, number of; P, P-value. SBP, systolic blood pressure; SD, standard deviation; WHR, waist-hip-ratio; WHRadjBMI, waist-hip-ratio adjusted for body mass index.

[a]Estimates per 1-unit higher GRS, corresponding to a predicted 1-SD higher obesity trait, outcome given in clinical units (mmHg) or SD-units, for linear regressions using the sex-specific estimates approach for constructing genetic risk scores

[b]P-value threshold set at <0.003 (=0.05/15) for 15 obesity trait-risk factor combinations in the study (DBP, SBP, fasting glucose, fasting insulin and smoking status)

[c]$P_{het}$ from Cochran's Q test between male and female estimates for matching traits. $P_{het}$ threshold set at <0.003 (=0.05/15) for 15 obesity trait-risk factor combinations in the study (DBP, SBP, fasting glucose, fasting insulin and smoking status)

**Table E. Genetic risk score associations with smoking status.**

| GRS | Outcome | Sex-Strata | N SNPs | OR (95% CI)[a] | P[b] | $P_{het}$[c] |
|---|---|---|---|---|---|---|
| BMI, combined | Having smoked/smoker | Combined | 565 | 1.32 (1.29, 1.36) | $3.52\times10^{-103}$ | - |
| BMI, men | Having smoked/smoker | Men | 565 | 1.38 (1.33, 1.44) | $2.97\times10^{-66}$ | $2.21\times10^{-04}$ |
| BMI, women | Having smoked/smoker | Women | 565 | 1.26 (1.22, 1.30) | $2.36\times10^{-39}$ | |
| WHR, combined | Having smoked/smoker | Combined | 324 | 1.24 (1.20, 1.28) | $3.74\times10^{-35}$ | - |
| WHR, men | Having smoked/smoker | Men | 324 | 1.46 (1.38, 1.55) | $4.63\times10^{-38}$ | $8.04\times10^{-14}$ |
| WHR, women | Having smoked/smoker | Women | 324 | 1.12 (1.08, 1.17) | $2.21\times10^{-09}$ | |
| WHRadjBMI, combined | Having smoked/smoker | Combined | 337 | 1.03 (1.00, 1.06) | 0.06 | - |
| WHRadjBMI, men | Having smoked/smoker | Men | 337 | 1.10 (1.04, 1.16) | 0.001 | 0.008 |
| WHRadjBMI, women | Having smoked/smoker | Women | 337 | 1.00 (0.97, 1.04) | 0.83 | |

BMI, body mass index; CI, confidence interval; GRS, genetic risk score; N, number of; OR, odds ratio; P, P-value; WHR, waist-hip ratio; WHRadjBMI, waist-hip-ratio adjusted for body mass index.

[a]Odds ratio per 1-unit change in the GRSs, using the sex-specific estimates approach for constructing genetic risk scores

[b]P-value threshold set at <0.003 (=0.05/15) for 15 obesity trait-risk factor combinations in the study (diastolic and systolic blood pressure, fasting glucose, fasting insulin and smoking status)

[c]$P_{het}$ from Cochran's Q test for comparisons between male and female estimates for the same obesity trait. $P_{het}$ threshold set at <0.003 (=0.05/15) for 15 obesity trait-risk factor combinations in the study (diastolic and systolic blood pressure, fasting glucose, fasting insulin and smoking status)

**Table F. Logistic regressions of the genetic risk scores with disease outcomes.**

| Outcome | Sex-strata | N cases | BMI OR (95% CI)[a] | BMI P[b] | BMI $P_{het}$[c] | WHR OR (95% CI)[a] | WHR P[b] | WHR $P_{het}$[c] | WHRadjBMI OR (95% CI)[a] | WHRadjBMI P[b] | WHRadjBMI $P_{het}$[c] |
|---|---|---|---|---|---|---|---|---|---|---|---|
| Type 2 Diabetes | Combined | 18,329 | 3.08 (2.90,3.28) | $4.40\times10^{-272}$ | - | 3.61 (3.32,3.93) | $5.40\times10^{-199}$ | - | 2.14 (1.99,2.30) | $2.80\times10^{-91}$ | - |
| Type 2 Diabetes | Men | 11,785 | 2.73 (2.53,2.95) | $1.70\times10^{-142}$ | $3.40\times10^{-05}$ | 3.21 (2.85,3.63) | $8.40\times10^{-80}$ | 0.10 | 1.90 (1.69,2.13) | $1.70\times10^{-27}$ | 0.08 |
| Type 2 Diabetes | Women | 6,544 | 3.58 (3.23,3.96) | $2.60\times10^{-134}$ | | 3.68 (3.29,4.12) | $4.10\times10^{-115}$ | | 2.17 (1.97,2.39) | $1.30\times10^{-55}$ | |
| CAD | Combined | 36,022 | 1.66 (1.59,1.74) | $6.50\times10^{-102}$ | - | 1.76 (1.66,1.88) | $1.80\times10^{-71}$ | - | 1.40 (1.33,1.48) | $9.50\times10^{-34}$ | - |
| CAD | Men | 24,444 | 1.63 (1.54,1.73) | $1.00\times10^{-64}$ | 0.63 | 1.73 (1.58,1.89) | $8.90\times10^{-34}$ | 0.19 | 1.40 (1.29,1.53) | $3.30\times10^{-15}$ | 0.22 |
| CAD | Women | 11,578 | 1.67 (1.55,1.81) | $1.70\times10^{-38}$ | | 1.59 (1.46,1.74) | $4.70\times10^{-26}$ | | 1.31 (1.22,1.41) | $1.10\times10^{-12}$ | |
| Breast Cancer | Women | 14,325 | 0.98 (0.91,1.05) | 0.56 | - | 0.95 (0.88,1.03) | 0.23 | - | 0.97 (0.90,1.03) | 0.30 | - |
| CLD | Combined | 1,367 | 1.64 (1.32,2.05) | $1.00\times10^{-05}$ | - | 1.88 (1.40,2.53) | $2.90\times10^{-05}$ | - | 1.34 (1.03,1.74) | 0.03 | - |
| CLD | Men | 824 | 1.40 (1.05,1.85) | 0.02 | 0.09 | 1.64 (1.06,2.54) | 0.03 | 0.76 | 1.38 (0.91,2.11) | 0.13 | 0.51 |
| CLD | Women | 543 | 2.04 (1.45,2.88) | $4.70\times10^{-05}$ | | 1.79 (1.22,2.63) | 0.003 | | 1.16 (0.83,1.61) | 0.39 | |
| Colorectal Cancer | Combined | 5,530 | 1.01 (0.91,1.13) | 0.83 | - | 1.19 (1.02,1.38) | 0.02 | - | 1.14 (1.00,1.30) | 0.05 | - |
| Colorectal Cancer | Men | 3,150 | 1.06 (0.91,1.22) | 0.46 | 0.23 | 1.31 (1.04,1.64) | 0.02 | 0.13 | 1.28 (1.03,1.60) | 0.02 | 0.25 |
| Colorectal Cancer | Women | 2,380 | 0.92 (0.78,1.09) | 0.34 | | 1.05 (0.87,1.26) | 0.63 | | 1.10 (0.94,1.29) | 0.25 | |
| COPD | Combined | 14,673 | 1.66 (1.55,1.78) | $5.50\times10^{-47}$ | - | 1.47 (1.34,1.61) | $6.00\times10^{-16}$ | - | 1.04 (0.96,1.13) | 0.39 | - |
| COPD | Men | 7,874 | 1.55 (1.41,1.70) | $4.90\times10^{-20}$ | 0.12 | 1.87 (1.62,2.17) | $4.40\times10^{-17}$ | $9.20\times10^{-06}$ | 1.23 (1.07,1.42) | 0.004 | 0.007 |
| COPD | Women | 6,799 | 1.72 (1.56,1.90) | $7.60\times10^{-27}$ | | 1.24 (1.11,1.38) | $1.80\times10^{-04}$ | | 0.98 (0.89,1.07) | 0.62 | |
| Dementia | Combined | 1,030 | 1.36 (1.06,1.76) | 0.02 | - | 0.73 (0.52,1.03) | 0.07 | - | 0.68 (0.50,0.92) | 0.01 | - |

| Outcome | Sex-strata | N cases | BMI OR (95% CI)[a] | BMI P[b] | BMI $P_{het}$[c] | WHR OR (95% CI)[a] | WHR P[b] | WHR $P_{het}$[c] | WHRadjBMI OR (95% CI)[a] | WHRadjBMI P[b] | WHRadjBMI $P_{het}$[c] |
|---|---|---|---|---|---|---|---|---|---|---|---|
| Dementia | Men | 583 | 1.18 (0.84,1.65) | 0.34 | 0.13 | 0.58 (0.35,0.99) | 0.04 | 0.20 | 0.60 (0.36,0.98) | 0.04 | 0.30 |
| Dementia | Women | 447 | 1.75 (1.19,2.55) | 0.004 | | 0.91 (0.60,1.39) | 0.67 | | 0.83 (0.57,1.19) | 0.31 | |
| Infertility | Combined | 1,678 | 1.01 (0.83,1.24) | 0.89 | - | 1.06 (0.81,1.39) | 0.67 | - | 1.00 (0.79,1.26) | 0.97 | - |
| Infertility | Men | 85 | 0.68 (0.28,1.63) | 0.39 | 0.35 | 2.14 (0.55,8.39) | 0.27 | 0.3 | 1.37 (0.37,5.08) | 0.64 | 0.62 |
| Infertility | Women | 1,593 | 1.04 (0.85,1.28) | 0.67 | | 1.03 (0.83,1.30) | 0.77 | | 0.98 (0.81,1.19) | 0.86 | |
| Lung Cancer | Combined | 2,716 | 1.33 (1.14,1.56) | $3.40\times10^{-04}$ | - | 1.16 (0.94,1.44) | 0.16 | - | 0.94 (0.78,1.14) | 0.54 | - |
| Lung Cancer | Men | 1,477 | 1.19 (0.96,1.47) | 0.11 | 0.16 | 1.02 (0.74,1.42) | 0.89 | 0.47 | 0.79 (0.57,1.08) | 0.14 | 0.20 |
| Lung Cancer | Women | 1,239 | 1.48 (1.18,1.87) | $7.00\times10^{-04}$ | | 1.19 (0.93,1.54) | 0.17 | | 1.01 (0.81,1.26) | 0.92 | |
| NAFLD | Combined | 1,684 | 2.92 (2.39,3.56) | $5.70\times10^{-26}$ | - | 2.69 (2.06,3.51) | $3.70\times10^{-13}$ | - | 1.63 (1.29,2.06) | $4.80\times10^{-05}$ | - |
| NAFLD | Men | 909 | 2.71 (2.08,3.55) | $2.40\times10^{-13}$ | 0.54 | 2.40 (1.58,3.64) | $4.30\times10^{-05}$ | 0.7 | 1.31 (0.88,1.96) | 0.18 | 0.41 |
| NAFLD | Women | 775 | 3.07 (2.31,4.10) | $2.00\times10^{-14}$ | | 2.66 (1.93,3.66) | $2.10\times10^{-09}$ | | 1.61 (1.22,2.12) | $7.30\times10^{-04}$ | |
| Renal Failure | Combined | 9,606 | 1.67 (1.53,1.82) | $2.70\times10^{-32}$ | - | 1.62 (1.44,1.81) | $1.20\times10^{-16}$ | - | 1.16 (1.05,1.29) | 0.003 | - |
| Renal Failure | Men | 5,697 | 1.61 (1.44,1.79) | $1.60\times10^{-17}$ | 0.26 | 1.94 (1.63,2.30) | $2.90\times10^{-14}$ | $5.30\times10^{-04}$ | 1.31 (1.11,1.54) | 0.001 | 0.05 |
| Renal Failure | Women | 3,909 | 1.77 (1.56,2.02) | $6.10\times10^{-18}$ | | 1.31 (1.13,1.51) | $3.00\times10^{-04}$ | | 1.06 (0.94,1.20) | 0.34 | |
| Renal Failure - Acute | Combined | 4,686 | 1.82 (1.61,2.05) | $2.70\times10^{-22}$ | - | 1.57 (1.33,1.84) | $5.20\times10^{-08}$ | - | 1.12 (0.97,1.29) | 0.12 | - |
| Renal Failure - Acute | Men | 3,035 | 1.73 (1.49,2.01) | $3.40\times10^{-13}$ | 0.32 | 1.87 (1.48,2.35) | $1.20\times10^{-07}$ | 0.01 | 1.18 (0.94,1.47) | 0.15 | 0.54 |
| Renal Failure - Acute | Women | 1,651 | 1.97 (1.61,2.40) | $2.30\times10^{-11}$ | | 1.25 (1.00,1.56) | 0.05 | | 1.07 (0.89,1.30) | 0.47 | |

| Outcome | Sex-strata | N cases | BMI OR (95% CI)[a] | BMI P[b] | BMI $P_{het}$[c] | WHR OR (95% CI)[a] | WHR P[b] | WHR $P_{het}$[c] | WHRadjBMI OR (95% CI)[a] | WHRadjBMI P[b] | WHRadjBMI $P_{het}$[c] |
|---|---|---|---|---|---|---|---|---|---|---|---|
| Renal Failure - Chronic | Combined | 4,599 | 1.81 (1.60,2.05) | $9.90\times10^{-22}$ | - | 1.74 (1.48,2.05) | $2.50\times10^{-11}$ | - | 1.22 (1.06,1.41) | 0.006 | - |
| Renal Failure - Chronic | Men | 2,581 | 1.87 (1.59,2.19) | $2.50\times10^{-14}$ | 0.71 | 2.33 (1.82,3.00) | $3.40\times10^{-11}$ | $1.30\times10^{-04}$ | 1.67 (1.31,2.12) | $3.00\times10^{-05}$ | $3.50\times10^{-04}$ |
| Renal Failure - Chronic | Women | 2,018 | 1.78 (1.49,2.13) | $3.00\times10^{-10}$ | | 1.25 (1.02,1.53) | 0.03 | | 0.97 (0.82,1.16) | 0.75 | |
| Stroke | Combined | 10,788 | 1.40 (1.30,1.52) | $1.00\times10^{-16}$ | - | 1.35 (1.21,1.50) | $3.70\times10^{-08}$ | - | 1.15 (1.04,1.26) | 0.005 | - |
| Stroke | Men | 6,334 | 1.42 (1.28,1.57) | $3.80\times10^{-11}$ | 0.84 | 1.57 (1.33,1.84) | $5.80\times10^{-08}$ | 0.009 | 1.22 (1.05,1.42) | 0.01 | 0.40 |
| Stroke | Women | 4,454 | 1.39 (1.23,1.57) | $8.60\times10^{-08}$ | | 1.18 (1.03,1.35) | 0.02 | | 1.12 (1.00,1.26) | 0.05 | |
| Stroke - Hemorrhagic | Combined | 1,907 | 1.25 (1.03,1.50) | 0.02 | - | 1.37 (1.07,1.76) | 0.01 | - | 1.15 (0.92,1.44) | 0.2 | - |
| Stroke - Hemorrhagic | Men | 928 | 1.24 (0.95,1.62) | 0.11 | 0.90 | 1.38 (0.91,2.09) | 0.13 | 0.54 | 1.11 (0.74,1.64) | 0.62 | 0.99 |
| Stroke - Hemorrhagic | Women | 979 | 1.27 (0.98,1.64) | 0.07 | | 1.18 (0.89,1.57) | 0.25 | | 1.10 (0.86,1.41) | 0.44 | |
| Stroke - Ischemic | Combined | 3,351 | 1.38 (1.20,1.59) | $8.10\times10^{-06}$ | - | 1.20 (1.00,1.46) | 0.06 | - | 1.16 (0.98,1.37) | 0.09 | - |
| Stroke - Ischemic | Men | 2,171 | 1.43 (1.20,1.70) | $6.50\times10^{-05}$ | 0.56 | 1.33 (1.01,1.74) | 0.04 | 0.33 | 1.27 (0.98,1.65) | 0.07 | 0.72 |
| Stroke - Ischemic | Women | 1,180 | 1.31 (1.04,1.65) | 0.02 | | 1.10 (0.85,1.43) | 0.47 | | 1.19 (0.95,1.49) | 0.12 | |
| Type 1 Diabetes | Combined | 1,503 | 1.65 (1.34,2.04) | $3.10\times10^{-06}$ | - | 1.15 (0.86,1.52) | 0.35 | - | 1.06 (0.83,1.36) | 0.64 | - |
| Type 1 Diabetes | Men | 826 | 1.46 (1.10,1.93) | 0.009 | 0.23 | 1.15 (0.74,1.78) | 0.54 | 0.81 | 1.29 (0.85,1.97) | 0.23 | 0.22 |
| Type 1 Diabetes | Women | 677 | 1.89 (1.39,2.57) | $5.40\times10^{-05}$ | | 1.07 (0.76,1.51) | 0.69 | | 0.94 (0.70,1.26) | 0.67 | |

BMI, body mass index; CI, confidence interval; CAD, coronary artery disease; COPD, chronic obstructive pulmonary disease; NAFLD, non-alcoholic fatty liver disease; OR, odds ratio; P, P-value.

For a visual presentation of the data see Figure C.

[a]Results per 1-unit higher weighted GRS, corresponding to a predicted 1-SD higher obesity trait, and for the sex-specific estimates approach for constructing genetic risk scores

[b]P-value threshold set at <0.001 (=0.05/51) for number of obesity GRS-outcome combinations tested
[c]$P_{het}$ from Cochran's Q test for comparisons between male and female estimates for matching traits, threshold set at <0.001 (=0.05/48) for number of obesity GRS-outcome combinations tested (fewer Cochran's Q tests since breast cancer was evaluated in women only)

**Table G. Mendelian randomization analyses with disease outcomes, unadjusted and adjusted for smoking status.**

| Exposure | Outcome | Sex-Strata | OR (95% CI), unadjusted[a] | P, unadjusted[b] | $P_{het}$, unadjusted[c] | OR (95% CI), adjusted[a] | P, adjusted[b] | $P_{het}$, adjusted[c] | % Change[d] |
|---|---|---|---|---|---|---|---|---|---|
| BMI | CAD | Combined | 1.66 (1.58, 1.74) | $8.21\times10^{-100}$ | - | 1.61 (1.54, 1.69) | $8.99\times10^{-88}$ | - | -3 |
| BMI | CAD | Men | 1.63 (1.54, 1.72) | $3.82\times10^{-63}$ | 0.49 | 1.57 (1.49, 1.67) | $2.85\times10^{-54}$ | 0.37 | -4 |
| BMI | CAD | Women | 1.68 (1.55, 1.82) | $8.02\times10^{-38}$ | | 1.65 (1.52, 1.78) | $1.07\times10^{-34}$ | | -2 |
| BMI | CLD | Combined | 1.64 (1.32, 2.05) | $8.51\times10^{-06}$ | - | 1.58 (1.27, 1.97) | $4.80\times10^{-05}$ | - | -4 |
| BMI | CLD | Men | 1.38 (1.05, 1.82) | 0.02 | 0.07 | 1.31 (1.00, 1.74) | 0.05 | 0.06 | -5 |
| BMI | CLD | Women | 2.08 (1.47, 2.94) | $3.28\times10^{-05}$ | | 2.03 (1.43, 2.87) | $6.63\times10^{-05}$ | | -2 |
| BMI | COPD | Combined | 1.66 (1.55, 1.78) | $1.27\times10^{-46}$ | - | 1.53 (1.42, 1.64) | $3.62\times10^{-32}$ | - | -8 |
| BMI | COPD | Men | 1.54 (1.41, 1.70) | $6.04\times10^{-20}$ | 0.1 | 1.41 (1.29, 1.55) | $8.74\times10^{-13}$ | 0.06 | -8 |
| BMI | COPD | Women | 1.74 (1.57, 1.92) | $1.37\times10^{-26}$ | | 1.61 (1.46, 1.79) | $5.25\times10^{-20}$ | | -7 |
| BMI | Lung Cancer | Combined | 1.33 (1.14, 1.55) | $3.71\times10^{-04}$ | - | 1.20 (1.03, 1.41) | 0.02 | - | -10 |
| BMI | Lung Cancer | Men | 1.18 (0.96, 1.46) | 0.11 | 0.15 | 1.06 (0.86, 1.31) | 0.57 | 0.11 | -10 |
| BMI | Lung Cancer | Women | 1.48 (1.18, 1.87) | $7.69\times10^{-04}$ | | 1.37 (1.08, 1.72) | 0.008 | | -7 |
| BMI | NAFLD | Combined | 2.89 (2.37, 3.53) | $7.16\times10^{-26}$ | - | 2.84 (2.32, 3.46) | $1.18\times10^{-24}$ | - | -2 |
| BMI | NAFLD | Men | 2.67 (2.05, 3.47) | $2.72\times10^{-13}$ | 0.46 | 2.63 (2.01, 3.43) | $1.19\times10^{-12}$ | 0.48 | -1 |
| BMI | NAFLD | Women | 3.10 (2.32, 4.14) | $2.30\times10^{-14}$ | | 3.03 (2.27, 4.06) | $8.57\times10^{-14}$ | | -2 |
| BMI | Renal Failure | Combined | 1.66 (1.53, 1.81) | $4.68\times10^{-32}$ | - | 1.63 (1.50, 1.78) | $2.59\times10^{-29}$ | - | -2 |
| BMI | Renal Failure | Men | 1.60 (1.43, 1.78) | $1.96\times10^{-17}$ | 0.2 | 1.56 (1.40, 1.74) | $1.11\times10^{-15}$ | 0.19 | -3 |
| BMI | Renal Failure | Women | 1.78 (1.56, 2.03) | $8.93\times10^{-18}$ | | 1.75 (1.54, 2.00) | $9.12\times10^{-17}$ | | -2 |
| BMI | Renal Failure - Acute | Combined | 1.81 (1.60, 2.04) | $3.75\times10^{-22}$ | - | 1.76 (1.56, 1.99) | $4.49\times10^{-20}$ | - | -3 |
| BMI | Renal Failure - Acute | Men | 1.72 (1.48, 1.99) | $3.81\times10^{-13}$ | 0.27 | 1.67 (1.44, 1.94) | $1.01\times10^{-11}$ | 0.24 | -3 |
| BMI | Renal Failure - Acute | Women | 1.98 (1.62, 2.42) | $2.86\times10^{-11}$ | | 1.94 (1.58, 2.37) | $1.23\times10^{-10}$ | | -2 |
| BMI | Renal Failure - Chronic | Combined | 1.81 (1.60, 2.04) | $1.32\times10^{-21}$ | - | 1.77 (1.57, 2.00) | $3.81\times10^{-20}$ | - | -2 |
| BMI | Renal Failure - Chronic | Men | 1.85 (1.58, 2.17) | $2.92\times10^{-14}$ | 0.81 | 1.81 (1.54, 2.13) | $3.58\times10^{-13}$ | 0.85 | -2 |
| BMI | Renal Failure - Chronic | Women | 1.79 (1.49, 2.15) | $3.55\times10^{-10}$ | | 1.77 (1.47, 2.12) | $1.03\times10^{-09}$ | | -1 |
| BMI | Stroke | Combined | 1.40 (1.29, 1.52) | $1.16\times10^{-16}$ | - | 1.37 (1.26, 1.48) | $2.18\times10^{-14}$ | - | -2 |

| Exposure | Outcome | Sex-Strata | OR (95% CI), unadjusted[a] | P, unadjusted[b] | $P_{het}$, unadjusted[c] | OR (95% CI), adjusted[a] | P, adjusted[b] | $P_{het}$, adjusted[c] | % Change[d] |
|---|---|---|---|---|---|---|---|---|---|
| BMI | Stroke | Men | 1.41 (1.27, 1.56) | $4.10\times10^{-11}$ | 0.91 | 1.37 (1.24, 1.52) | $2.01\times10^{-09}$ | 0.99 | -3 |
| BMI | Stroke | Women | 1.40 (1.24, 1.58) | $8.93\times10^{-08}$ | | 1.37 (1.21, 1.55) | $4.22\times10^{-07}$ | | -2 |
| BMI | Stroke - Ischaemic | Combined | 1.38 (1.20, 1.59) | $8.15\times10^{-06}$ | - | 1.35 (1.17, 1.55) | $3.54\times10^{-05}$ | - | -2 |
| BMI | Stroke - Ischaemic | Men | 1.42 (1.19, 1.68) | $6.62\times10^{-05}$ | 0.6 | 1.38 (1.16, 1.64) | $2.60\times10^{-04}$ | 0.67 | -3 |
| BMI | Stroke - Ischaemic | Women | 1.31 (1.04, 1.66) | 0.02 | | 1.30 (1.02, 1.64) | 0.03 | | -1 |
| BMI | Type 1 Diabetes | Combined | 1.68 (1.35, 2.08) | $3.13\times10^{-06}$ | - | 1.68 (1.35, 2.09) | $3.27\times10^{-06}$ | - | 0 |
| BMI | Type 1 Diabetes | Men | 1.47 (1.10, 1.96) | 0.009 | 0.21 | 1.47 (1.10, 1.96) | 0.009 | 0.21 | 0 |
| BMI | Type 1 Diabetes | Women | 1.93 (1.40, 2.67) | $5.55\times10^{-05}$ | | 1.94 (1.41, 2.68) | $5.24\times10^{-05}$ | | 1 |
| BMI | Type 2 Diabetes | Combined | 3.18 (2.97, 3.39) | $<1\times10^{-200}$ | - | 3.12 (2.92, 3.33) | $<1\times10^{-200}$ | - | -2 |
| BMI | Type 2 Diabetes | Men | 2.79 (2.58, 3.03) | $5.70\times10^{-135}$ | $1.44\times10^{-05}$ | 2.73 (2.51, 2.96) | $1.72\times10^{-126}$ | $5.71\times10^{-06}$ | -2 |
| BMI | Type 2 Diabetes | Women | 3.77 (3.38, 4.20) | $1.67\times10^{-128}$ | | 3.74 (3.36, 4.17) | $2.67\times10^{-126}$ | | -1 |
| WHR | CAD | Combined | 1.74 (1.63, 1.85) | $9.19\times10^{-70}$ | - | 1.70 (1.60, 1.81) | $7.80\times10^{-64}$ | - | -2 |
| WHR | CAD | Men | 1.73 (1.58, 1.90) | $1.12\times10^{-32}$ | 0.11 | 1.68 (1.53, 1.84) | $8.01\times10^{-28}$ | 0.24 | -3 |
| WHR | CAD | Women | 1.57 (1.44, 1.70) | $9.80\times10^{-26}$ | | 1.55 (1.43, 1.69) | $1.52\times10^{-24}$ | | -1 |
| WHR | CLD | Combined | 1.85 (1.39, 2.47) | $2.91\times10^{-05}$ | - | 1.80 (1.35, 2.41) | $7.27\times10^{-05}$ | - | -3 |
| WHR | COPD | Combined | 1.45 (1.33, 1.59) | $7.81\times10^{-16}$ | - | 1.37 (1.25, 1.50) | $3.62\times10^{-11}$ | - | -6 |
| WHR | COPD | Men | 1.89 (1.62, 2.19) | $6.64\times10^{-17}$ | $3.71\times10^{-06}$ | 1.71 (1.47, 1.99) | $5.36\times10^{-12}$ | $8.84\times10^{-05}$ | -10 |
| WHR | COPD | Women | 1.22 (1.10, 1.36) | $2.12\times10^{-04}$ | | 1.18 (1.06, 1.31) | 0.003 | | -3 |
| WHR | NAFLD | Combined | 2.62 (2.02, 3.39) | $4.06\times10^{-13}$ | - | 2.59 (1.99, 3.37) | $1.21\times10^{-12}$ | - | -1 |
| WHR | NAFLD | Men | 2.38 (1.57, 3.61) | $4.41\times10^{-05}$ | 0.79 | 2.36 (1.54, 3.61) | $8.14\times10^{-05}$ | 0.79 | -1 |
| WHR | NAFLD | Women | 2.55 (1.88, 3.47) | $2.16\times10^{-09}$ | | 2.53 (1.86, 3.45) | $3.46\times10^{-09}$ | | -1 |
| WHR | Renal Failure | Combined | 1.60 (1.43, 1.78) | $1.25\times10^{-16}$ | - | 1.58 (1.41, 1.76) | $1.80\times10^{-15}$ | - | -1 |
| WHR | Renal Failure | Men | 1.93 (1.63, 2.29) | $4.15\times10^{-14}$ | $3.64\times10^{-04}$ | 1.90 (1.59, 2.26) | $6.90\times10^{-13}$ | $6.35\times10^{-04}$ | -2 |
| WHR | Renal Failure | Women | 1.30 (1.13, 1.49) | $2.63\times10^{-04}$ | | 1.29 (1.12, 1.48) | $4.29\times10^{-04}$ | | -1 |
| WHR | Renal Failure - Acute | Combined | 1.55 (1.32, 1.81) | $4.92\times10^{-08}$ | - | 1.52 (1.30, 1.78) | $2.50\times10^{-07}$ | - | -2 |
| WHR | Renal Failure - Acute | Men | 1.86 (1.48, 2.35) | $1.33\times10^{-07}$ | 0.01 | 1.81 (1.43, 2.29) | $8.63\times10^{-07}$ | 0.02 | -3 |
| WHR | Renal Failure - Acute | Women | 1.24 (1.00, 1.54) | 0.04 | | 1.23 (0.99, 1.52) | 0.06 | | -1 |

| Exposure | Outcome | Sex-Strata | OR (95% CI), unadjusted[a] | P, unadjusted[b] | $P_{het}$, unadjusted[c] | OR (95% CI), adjusted[a] | P, adjusted[b] | $P_{het}$, adjusted[c] | % Change[d] |
|---|---|---|---|---|---|---|---|---|---|
| WHR | Renal Failure - Chronic | Combined | 1.72 (1.47, 2.01) | $2.06\times10^{-11}$ | - | 1.70 (1.45, 1.99) | $8.95\times10^{-11}$ | - | -1 |
| WHR | Renal Failure - Chronic | Men | 2.33 (1.81, 2.99) | $4.16\times10^{-11}$ | $1.04\times10^{-04}$ | 2.29 (1.77, 2.96) | $2.44\times10^{-10}$ | $1.68\times10^{-04}$ | -2 |
| WHR | Renal Failure - Chronic | Women | 1.24 (1.03, 1.51) | 0.03 | | 1.24 (1.02, 1.50) | 0.03 | | 0 |
| WHR | Stroke | Combined | 1.34 (1.21, 1.49) | $3.73\times10^{-08}$ | - | 1.32 (1.19, 1.46) | $3.07\times10^{-07}$ | - | -1 |
| WHR | Stroke | Men | 1.57 (1.33, 1.84) | $6.44\times10^{-08}$ | 0.006 | 1.52 (1.29, 1.79) | $7.78\times10^{-07}$ | 0.01 | -3 |
| WHR | Stroke | Women | 1.17 (1.03, 1.33) | 0.02 | | 1.16 (1.02, 1.32) | 0.02 | | -1 |
| WHR | Type 2 Diabetes | Combined | 3.62 (3.32, 3.95) | $8.82\times10^{-187}$ | - | 3.60 (3.30, 3.93) | $1.94\times10^{-181}$ | - | -1 |
| WHR | Type 2 Diabetes | Men | 3.36 (2.95, 3.83) | $7.48\times10^{-74}$ | 0.41 | 3.31 (2.90, 3.78) | $3.76\times10^{-69}$ | 0.33 | -1 |
| WHR | Type 2 Diabetes | Women | 3.61 (3.22, 4.04) | $2.07\times10^{-110}$ | | 3.61 (3.22, 4.04) | $1.80\times10^{-109}$ | | 0 |
| WHRadjBMI | CAD | Combined | 1.39 (1.31, 1.46) | $1.84\times10^{-33}$ | - | 1.38 (1.31, 1.46) | $1.32\times10^{-32}$ | - | -1 |
| WHRadjBMI | CAD | Men | 1.40 (1.29, 1.53) | $5.06\times10^{-15}$ | 0.16 | 1.39 (1.28, 1.52) | $3.24\times10^{-14}$ | 0.21 | -1 |
| WHRadjBMI | CAD | Women | 1.30 (1.21, 1.39) | $1.35\times10^{-12}$ | | 1.30 (1.21, 1.39) | $1.59\times10^{-12}$ | | 0 |
| WHRadjBMI | NAFLD | Combined | 1.60 (1.28, 2.01) | $4.81\times10^{-05}$ | - | 1.60 (1.27, 2.01) | $5.26\times10^{-05}$ | - | 0 |
| WHRadjBMI | NAFLD | Men | 1.30 (0.88, 1.93) | 0.18 | 0.43 | 1.30 (0.88, 1.92) | 0.19 | 0.42 | 0 |
| WHRadjBMI | NAFLD | Women | 1.58 (1.21, 2.06) | $7.28\times10^{-04}$ | | 1.58 (1.21, 2.06) | $7.34\times10^{-04}$ | | 0 |
| WHRadjBMI | Renal Failure - Chronic | Men | 1.64 (1.30, 2.08) | $3.11\times10^{-05}$ | $3.27\times10^{-04}$ | 1.64 (1.29, 2.07) | $4.19\times10^{-05}$ | $3.84\times10^{-04}$ | 0 |
| WHRadjBMI | Renal Failure - Chronic | Women | 0.97 (0.82, 1.15) | 0.74 | | 0.97 (0.82, 1.15) | 0.72 | | 0 |
| WHRadjBMI | Type 2 Diabetes | Combined | 2.10 (1.96, 2.26) | $2.31\times10^{-89}$ | - | 2.10 (1.95, 2.26) | $1.16\times10^{-88}$ | - | 0 |
| WHRadjBMI | Type 2 Diabetes | Men | 1.90 (1.69, 2.14) | $6.47\times10^{-27}$ | 0.15 | 1.89 (1.68, 2.13) | $3.14\times10^{-26}$ | 0.13 | -1 |
| WHRadjBMI | Type 2 Diabetes | Women | 2.12 (1.93, 2.33) | $7.47\times10^{-55}$ | | 2.12 (1.93, 2.33) | $7.99\times10^{-55}$ | | 0 |

BMI, body mass index; CI, confidence interval; CAD, coronary artery disease; COPD, chronic obstructive pulmonary disease; CLD, chronic liver disease; NAFLD, non-alcoholic fatty liver disease; OR, odds ratio; P, P-value; SD, standard deviation.

[a]Results per 1-SD higher obesity trait, using the sex-specific estimates approach for constructing genetic risk scores.

[b]P-value-threshold set at <0.002 (=0.05/51) for 51 obesity trait-disease outcome combinations

[c]$P_{het}$-values from Cochran's Q test for comparisons between male and female estimates for matching traits. $P_{het}$-threshold set at <0.001 (=0.05/48) for 48 male-female disease estimate comparisons in the study (fewer Cochran's Q tests since breast cancer was investigated in women only)

[d]% Change is the relative change in the adjusted for smoking status point estimate compared to the unadjusted for smoking status point estimate.

**Table H. MR-Egger intercept test and MR-Egger estimates for the disease outcomes.**

| Outcome | Exposure | Sex-strata | N SNPs used (N SNPs GRS)[a] | Intercept (95% CI) | P intercept[b] | OR (95% CI)[c] | P estimate[d] | $P_{het}$[e] |
|---|---|---|---|---|---|---|---|---|
| CAD | BMI | Combined | 565 (565) | 0.004 (0.001, 0.007) | 0.004 | 1.33 (1.10, 1.59) | 0.003 | - |
| CAD | BMI | Men | 565 (565) | 0.004 (0.001, 0.007) | 0.01 | 1.35 (1.12, 1.62) | 0.002 | 0.6 |
| CAD | BMI | Women | 565 (565) | 0.006 (0.002, 0.010) | 0.005 | 1.24 (0.97, 1.58) | 0.08 | |
| CAD | WHR | Combined | 324 (324) | 0.008 (0.003, 0.012) | 0.001 | 1.13 (0.85, 1.52) | 0.40 | - |
| CAD | WHR | Men | 324 (324) | 0.004 (-0.000, 0.008) | 0.07 | 1.38 (1.02, 1.87) | 0.04 | 0.94 |
| CAD | WHR | Women | 324 (324) | 0.004 (-0.001, 0.008) | 0.12 | 1.36 (1.07, 1.72) | 0.01 | |
| CAD | WHRadjBMI | Combined | 337 (337) | -0.001 (-0.005, 0.003) | 0.67 | 1.45 (1.19, 1.78) | $2.99\times10^{-04}$ | - |
| CAD | WHRadjBMI | Men | 337 (337) | 0.002 (-0.002, 0.006) | 0.29 | 1.25 (0.97, 1.60) | 0.08 | 0.49 |
| CAD | WHRadjBMI | Women | 337 (337) | -0.002 (-0.006, 0.002) | 0.43 | 1.39 (1.16, 1.66) | $2.72\times10^{-04}$ | |
| CLD | BMI | Combined | 565 (565) | 0.008 (-0.002, 0.018) | 0.11 | 1.05 (0.57, 1.92) | 0.88 | - |
| CLD | BMI | Men | 565 (565) | 0.009 (-0.003, 0.020) | 0.13 | 0.87 (0.43, 1.74) | 0.69 | 0.32 |
| CLD | BMI | Women | 565 (565) | 0.005 (-0.010, 0.021) | 0.51 | 1.56 (0.63, 3.91) | 0.34 | |
| CLD | WHR | Combined | 323 (324) | -0.003 (-0.016, 0.011) | 0.70 | 2.25 (0.96, 5.24) | 0.06 | - |
| COPD | BMI | Combined | 565 (565) | 0.003 (-0.000, 0.007) | 0.08 | 1.40 (1.11, 1.77) | 0.004 | - |
| COPD | BMI | Men | 565 (565) | 0.008 (0.004, 0.012) | $2.10\times10^{-04}$ | 1.03 (0.80, 1.32) | 0.83 | 0.01 |
| COPD | BMI | Women | 565 (565) | 0.000 (-0.005, 0.006) | 0.88 | 1.73 (1.26, 2.36) | $6.49\times10^{-04}$ | |
| COPD | WHR | Combined | 324 (324) | 0.006 (0.001, 0.011) | 0.02 | 1.04 (0.75, 1.44) | 0.82 | - |
| COPD | WHR | Men | 324 (324) | -0.002 (-0.007, 0.002) | 0.30 | 2.22 (1.60, 3.07) | $1.77\times10^{-06}$ | $1.68\times10^{-06}$ |
| COPD | WHR | Women | 324 (324) | 0.011 (0.005, 0.016) | $1.74\times10^{-04}$ | 0.77 (0.58, 1.02) | 0.07 | |
| Lung Cancer | BMI | Combined | 565 (565) | 0.006 (-0.001, 0.014) | 0.11 | 0.95 (0.60, 1.51) | 0.84 | - |
| Lung Cancer | BMI | Men | 565 (565) | 0.007 (-0.002, 0.016) | 0.12 | 0.82 (0.48, 1.41) | 0.47 | 0.46 |
| Lung Cancer | BMI | Women | 565 (565) | 0.005 (-0.006, 0.016) | 0.35 | 1.13 (0.60, 2.12) | 0.71 | |
| NAFLD | BMI | Combined | 565 (565) | 0.008 (-0.002, 0.017) | 0.10 | 1.91 (1.08, 3.39) | 0.03 | - |
| NAFLD | BMI | Men | 565 (565) | 0.000 (-0.011, 0.011) | 0.98 | 2.76 (1.44, 5.29) | 0.002 | 0.08 |
| NAFLD | BMI | Women | 565 (565) | 0.019 (0.005, 0.033) | 0.007 | 1.09 (0.48, 2.47) | 0.83 | |
| NAFLD | WHR | Combined | 324 (324) | 0.003 (-0.010, 0.016) | 0.68 | 2.32 (1.03, 5.24) | 0.04 | - |

| Outcome | Exposure | Sex-strata | N SNPs used (N SNPs GRS)[a] | Intercept (95% CI) | P intercept[b] | OR (95% CI)[c] | P estimate[d] | $P_{het}$[e] |
|---|---|---|---|---|---|---|---|---|
| NAFLD | WHR | Men | 323 (324) | 0.000 (-0.013, 0.013) | 0.95 | 2.40 (0.94, 6.13) | 0.07 | 0.94 |
| NAFLD | WHR | Women | 324 (324) | 0.004 (-0.011, 0.019) | 0.64 | 2.28 (1.06, 4.93) | 0.04 | |
| NAFLD | WHRadjBMI | Combined | 337 (337) | 0.002 (-0.009, 0.013) | 0.74 | 1.51 (0.83, 2.74) | 0.18 | - |
| NAFLD | WHRadjBMI | Men | 336 (337) | 0.010 (-0.002, 0.022) | 0.09 | 0.75 (0.34, 1.68) | 0.49 | 0.26 |
| NAFLD | WHRadjBMI | Women | 337 (337) | 0.005 (-0.009, 0.019) | 0.47 | 1.34 (0.73, 2.43) | 0.34 | |
| Renal Failure | BMI | Combined | 565 (565) | 0.003 (-0.001, 0.007) | 0.17 | 1.45 (1.14, 1.85) | 0.003 | - |
| Renal Failure | BMI | Men | 565 (565) | 0.002 (-0.002, 0.007) | 0.32 | 1.44 (1.09, 1.90) | 0.01 | 0.57 |
| Renal Failure | BMI | Women | 565 (565) | 0.002 (-0.004, 0.007) | 0.57 | 1.64 (1.16, 2.31) | 0.005 | |
| Renal Failure | WHR | Combined | 324 (324) | 0.007 (0.002, 0.013) | 0.006 | 1.06 (0.76, 1.47) | 0.73 | - |
| Renal Failure | WHR | Men | 324 (324) | 0.009 (0.004, 0.014) | $7.77\times10^{-04}$ | 1.14 (0.79, 1.63) | 0.48 | 0.35 |
| Renal Failure | WHR | Women | 324 (324) | 0.008 (0.002, 0.015) | 0.01 | 0.90 (0.64, 1.27) | 0.55 | |
| Renal Failure - Acute | BMI | Combined | 565 (565) | 0.001 (-0.005, 0.007) | 0.72 | 1.74 (1.23, 2.45) | 0.002 | - |
| Renal Failure - Acute | BMI | Men | 565 (565) | 0.001 (-0.005, 0.007) | 0.78 | 1.67 (1.14, 2.45) | 0.008 | 0.71 |
| Renal Failure - Acute | BMI | Women | 565 (565) | 0.001 (-0.008, 0.010) | 0.84 | 1.90 (1.12, 3.21) | 0.02 | |
| Renal Failure - Acute | WHR | Combined | 324 (324) | 0.008 (0.001, 0.016) | 0.03 | 0.97 (0.61, 1.54) | 0.88 | - |
| Renal Failure - Acute | WHR | Men | 324 (324) | 0.006 (-0.001, 0.013) | 0.09 | 1.29 (0.79, 2.12) | 0.31 | 0.10 |
| Renal Failure - Acute | WHR | Women | 324 (324) | 0.012 (0.003, 0.022) | 0.01 | 0.72 (0.44, 1.18) | 0.20 | |
| Renal Failure - Chronic | BMI | Combined | 565 (565) | 0.005 (-0.000, 0.011) | 0.05 | 1.36 (0.98, 1.91) | 0.07 | - |
| Renal Failure - Chronic | BMI | Men | 565 (565) | 0.002 (-0.005, 0.008) | 0.61 | 1.75 (1.17, 2.61) | 0.006 | 0.24 |
| Renal Failure - Chronic | BMI | Women | 565 (565) | 0.007 (-0.001, 0.015) | 0.07 | 1.20 (0.75, 1.94) | 0.45 | |
| Renal Failure - Chronic | WHR | Combined | 324 (324) | 0.008 (0.001, 0.016) | 0.03 | 1.08 (0.68, 1.72) | 0.74 | - |
| Renal Failure - Chronic | WHR | Men | 324 (324) | 0.009 (0.002, 0.016) | 0.02 | 1.36 (0.80, 2.29) | 0.25 | 0.10 |
| Renal Failure - Chronic | WHR | Women | 324 (324) | 0.011 (0.002, 0.020) | 0.01 | 0.77 (0.49, 1.20) | 0.24 | |
| Renal Failure - Chronic | WHRadjBMI | Men | 337 (337) | 0.002 (-0.005, 0.009) | 0.58 | 1.50 (0.95, 2.35) | 0.08 | 0.13 |
| Renal Failure - Chronic | WHRadjBMI | Women | 337 (337) | 0.000 (-0.008, 0.009) | 0.91 | 0.96 (0.68, 1.37) | 0.83 | |
| Stroke | BMI | Combined | 565 (565) | 0.005 (0.001, 0.009) | 0.01 | 1.08 (0.86, 1.36) | 0.50 | - |

| Outcome | Exposure | Sex-strata | N SNPs used (N SNPs GRS)[a] | Intercept (95% CI) | P intercept[b] | OR (95% CI)[c] | P estimate[d] | $P_{het}$[e] |
|---|---|---|---|---|---|---|---|---|
| Stroke | BMI | Men | 565 (565) | 0.003 (-0.002, 0.007) | 0.26 | 1.25 (0.96, 1.64) | 0.10 | 0.33 |
| Stroke | BMI | Women | 565 (565) | 0.006 (0.000, 0.012) | 0.04 | 1.01 (0.72, 1.42) | 0.94 | |
| Stroke | WHR | Combined | 324 (324) | 0.001 (-0.005, 0.006) | 0.80 | 1.31 (0.95, 1.82) | 0.10 | - |
| Stroke | WHR | Men | 324 (324) | -0.002 (-0.007, 0.003) | 0.37 | 1.82 (1.30, 2.55) | $4.75\times10^{-04}$ | 0.003 |
| Stroke | WHR | Women | 324 (324) | 0.006 (-0.000, 0.012) | 0.06 | 0.90 (0.65, 1.25) | 0.54 | |
| Stroke - Ischemic | BMI | Combined | 565 (565) | 0.004 (-0.003, 0.010) | 0.27 | 1.15 (0.78, 1.70) | 0.49 | - |
| Stroke - Ischemic | BMI | Men | 565 (565) | 0.002 (-0.006, 0.009) | 0.68 | 1.34 (0.87, 2.06) | 0.19 | 0.40 |
| Stroke - Ischemic | BMI | Women | 565 (565) | 0.006 (-0.005, 0.016) | 0.28 | 0.97 (0.52, 1.80) | 0.91 | |
| Type 1 Diabetes | BMI | Combined | 565 (565) | -0.008 (-0.018, 0.002) | 0.12 | 2.57 (1.41, 4.68) | 0.002 | - |
| Type 1 Diabetes | BMI | Men | 565 (565) | -0.001 (-0.013, 0.010) | 0.80 | 1.56 (0.78, 3.09) | 0.21 | 0.09 |
| Type 1 Diabetes | BMI | Women | 565 (565) | -0.013 (-0.027, 0.001) | 0.06 | 3.88 (1.71, 8.82) | 0.001 | |
| Type 2 Diabetes | BMI | Combined | 565 (565) | -0.001 (-0.007, 0.005) | 0.74 | 3.32 (2.35, 4.69) | $1.13\times10^{-11}$ | - |
| Type 2 Diabetes | BMI | Men | 565 (565) | 0.004 (-0.001, 0.009) | 0.13 | 2.24 (1.64, 3.07) | $4.35\times10^{-07}$ | 0.001 |
| Type 2 Diabetes | BMI | Women | 565 (565) | -0.006 (-0.013, 0.001) | 0.07 | 5.15 (3.44, 7.73) | $2.22\times10^{-15}$ | |
| Type 2 Diabetes | WHR | Combined | 324 (324) | 0.008 (0.002, 0.015) | 0.02 | 2.27 (1.50, 3.43) | $1.09\times10^{-04}$ | - |
| Type 2 Diabetes | WHR | Men | 324 (324) | 0.004 (-0.002, 0.010) | 0.15 | 2.52 (1.68, 3.78) | $7.89\times10^{-06}$ | 0.61 |
| Type 2 Diabetes | WHR | Women | 324 (324) | 0.012 (0.005, 0.018) | $4.51\times10^{-04}$ | 2.20 (1.57, 3.08) | $3.87\times10^{-06}$ | |
| Type 2 Diabetes | WHRadjBMI | Combined | 337 (337) | 0.005 (-0.002, 0.013) | 0.17 | 1.67 (1.11, 2.49) | 0.01 | - |
| Type 2 Diabetes | WHRadjBMI | Men | 337 (337) | 0.008 (0.001, 0.015) | 0.02 | 1.20 (0.76, 1.89) | 0.42 | 0.03 |
| Type 2 Diabetes | WHRadjBMI | Women | 337 (337) | -0.001 (-0.008, 0.007) | 0.85 | 2.24 (1.64, 3.06) | $3.61\times10^{-07}$ | |

BMI, body mass index; CI, confidence interval; CAD, coronary artery disease; COPD, chronic obstructive pulmonary disease; CLD, chronic liver disease; GRS, genetic risk score; NAFLD, non-alcoholic fatty liver disease; OR, odds ratio; P, P-value; SD, standard deviation.

[a]N SNPs used is the number of SNPs where summary statistics were successfully computed and that were included in the analyses

[b]P-value intercept threshold set at <0.002 (=0.05/25) for 25 obesity trait-disease outcome combinations investigated with the intercept test

[c]Results per 1-SD higher obesity trait, using the sex-specific estimates approach to construct genetic risk scores

[d]P-value estimate threshold set at <0.001 (=0.05/51) for 51 obesity trait-disease outcome combinations

[e]$P_{het}$-values from Cochran's Q test for comparisons between male and female MR-Egger estimates for matching traits. $P_{het}$-threshold set at <0.001 (=0.05/48) for 48 male-female comparisons in the study (fewer Cochran's Q tests since breast cancer investigated in women only).

**Table I. Logistic regressions of the genetic risk scores with disease outcomes using the same number of cases and controls for men and women compared to using all available cases and controls.**

| Outcome | GRS | N SNPs | Sex-Strata | OR (95% CI), all available cases and controls for men and women[a] | P, all available cases and controls for men and women[b] | $P_{het}$, all available cases and controls for men and women[c] | OR (95% CI), same number of cases and controls for men and women[a] | P, same number of cases and controls for men and women[b] | $P_{het}$, same number of cases and controls for men and women[c] |
|---|---|---|---|---|---|---|---|---|---|
| CAD | BMI, men | 565 | Men | 1.63 (1.54, 1.73) | $9.97\times10^{-65}$ | 0.63 | 1.66 (1.54, 1.80) | $8.73\times10^{-37}$ | 0.81 |
| CAD | BMI, women | 565 | Women | 1.67 (1.55, 1.81) | $1.70\times10^{-38}$ | | 1.69 (1.56, 1.82) | $8.24\times10^{-39}$ | |
| CAD | WHR, men | 324 | Men | 1.73 (1.58, 1.89) | $8.94\times10^{-34}$ | 0.19 | 1.76 (1.55, 1.99) | $2.50\times10^{-19}$ | 0.23 |
| CAD | WHR, women | 324 | Women | 1.59 (1.46, 1.74) | $4.67\times10^{-26}$ | | 1.60 (1.47, 1.75) | $4.44\times10^{-26}$ | |
| CAD | WHRadjBMI, men | 337 | Men | 1.40 (1.29, 1.53) | $3.31\times10^{-15}$ | 0.22 | 1.37 (1.22, 1.54) | $1.57\times10^{-07}$ | 0.52 |
| CAD | WHRadjBMI, women | 337 | Women | 1.31 (1.22, 1.41) | $1.15\times10^{-12}$ | | 1.31 (1.21, 1.41) | $2.65\times10^{-12}$ | |
| CLD | BMI, men | 565 | Men | 1.40 (1.05, 1.85) | 0.02 | 0.09 | 1.39 (0.98, 1.96) | 0.06 | 0.12 |
| CLD | BMI, women | 565 | Women | 2.04 (1.45, 2.88) | $4.71\times10^{-05}$ | | 2.04 (1.45, 2.88) | $4.74\times10^{-05}$ | |
| CLD | WHR, men | 324 | Men | 1.64 (1.06, 2.54) | 0.03 | 0.76 | 1.68 (0.98, 2.88) | 0.06 | 0.85 |
| CLD | WHR, women | 324 | Women | 1.79 (1.22, 2.63) | 0.003 | | 1.79 (1.22, 2.62) | 0.003 | |
| CLD | WHRadjBMI, men | 337 | Men | 1.38 (0.91, 2.11) | 0.13 | 0.51 | 1.73 (1.03, 2.91) | 0.04 | 0.19 |
| CLD | WHRadjBMI, women | 337 | Women | 1.16 (0.83, 1.61) | 0.39 | | 1.15 (0.83, 1.61) | 0.40 | |
| COPD | BMI, men | 565 | Men | 1.55 (1.41, 1.70) | $4.92\times10^{-20}$ | 0.12 | 1.57 (1.42, 1.73) | $1.96\times10^{-18}$ | 0.17 |
| COPD | BMI, women | 565 | Women | 1.72 (1.56, 1.90) | $7.59\times10^{-27}$ | | 1.73 (1.56, 1.91) | $7.23\times10^{-27}$ | |
| COPD | WHR, men | 324 | Men | 1.87 (1.62, 2.17) | $4.40\times10^{-17}$ | $9.22\times10^{-06}$ | 1.84 (1.57, 2.15) | $2.53\times10^{-14}$ | $6.06\times10^{-05}$ |
| COPD | WHR, women | 324 | Women | 1.24 (1.11, 1.38) | $1.80\times10^{-04}$ | | 1.24 (1.11, 1.39) | $1.39\times10^{-04}$ | |
| COPD | WHRadjBMI, men | 337 | Men | 1.23 (1.07, 1.42) | 0.004 | 0.007 | 1.11 (0.96, 1.29) | 0.17 | 0.18 |
| COPD | WHRadjBMI, women | 337 | Women | 0.98 (0.89, 1.07) | 0.62 | | 0.98 (0.89, 1.08) | 0.71 | |
| Colorectal Cancer | BMI, men | 565 | Men | 1.06 (0.91, 1.22) | 0.46 | 0.23 | 1.05 (0.89, 1.25) | 0.53 | 0.29 |
| Colorectal Cancer | BMI, women | 565 | Women | 0.92 (0.78, 1.09) | 0.34 | | 0.93 (0.79, 1.10) | 0.38 | |

| Outcome | GRS | N SNPs | Sex-Strata | OR (95% CI), all available cases and controls for men and women[a] | P, all available cases and controls for men and women[b] | $P_{het}$, all available cases and controls for men and women[c] | OR (95% CI), same number of cases and controls for men and women[a] | P, same number of cases and controls for men and women[b] | $P_{het}$, same number of cases and controls for men and women[c] |
|---|---|---|---|---|---|---|---|---|---|
| Colorectal Cancer | WHR, men | 324 | Men | 1.31 (1.04, 1.64) | 0.02 | 0.13 | 1.22 (0.94, 1.58) | 0.13 | 0.33 |
| Colorectal Cancer | WHR, women | 324 | Women | 1.05 (0.87, 1.26) | 0.63 | | 1.04 (0.87, 1.25) | 0.66 | |
| Colorectal Cancer | WHRadjBMI, men | 337 | Men | 1.28 (1.03, 1.60) | 0.02 | 0.25 | 1.23 (0.96, 1.57) | 0.11 | 0.46 |
| Colorectal Cancer | WHRadjBMI, women | 337 | Women | 1.10 (0.94, 1.29) | 0.25 | | 1.10 (0.94, 1.29) | 0.25 | |
| Dementia | BMI, men | 565 | Men | 1.18 (0.84, 1.65) | 0.34 | 0.13 | 0.99 (0.67, 1.45) | 0.95 | 0.04 |
| Dementia | BMI, women | 565 | Women | 1.75 (1.19, 2.55) | 0.004 | | 1.74 (1.19, 2.54) | 0.004 | |
| Dementia | WHR, men | 324 | Men | 0.58 (0.35, 0.99) | 0.04 | 0.20 | 0.57 (0.32, 1.04) | 0.07 | 0.22 |
| Dementia | WHR, women | 324 | Women | 0.91 (0.60, 1.39) | 0.67 | | 0.91 (0.59, 1.38) | 0.64 | |
| Dementia | WHRadjBMI, men | 337 | Men | 0.60 (0.36, 0.98) | 0.04 | 0.30 | 0.66 (0.37, 1.18) | 0.16 | 0.53 |
| Dementia | WHRadjBMI, women | 337 | Women | 0.83 (0.57, 1.19) | 0.31 | | 0.82 (0.57, 1.19) | 0.30 | |
| Infertility | BMI, men | 565 | Men | 0.68 (0.28, 1.63) | 0.39 | 0.35 | 0.68 (0.28, 1.63) | 0.39 | 0.12 |
| Infertility | BMI, women | 565 | Women | 1.04 (0.85, 1.28) | 0.67 | | 1.83 (0.77, 4.37) | 0.17 | |
| Infertility | WHR, men | 324 | Men | 2.14 (0.55, 8.39) | 0.27 | 0.30 | 2.14 (0.55, 8.39) | 0.27 | 0.61 |
| Infertility | WHR, women | 324 | Women | 1.03 (0.83, 1.30) | 0.77 | | 1.38 (0.52, 3.64) | 0.52 | |
| Infertility | WHRadjBMI, men | 337 | Men | 1.37 (0.37, 5.08) | 0.64 | 0.62 | 1.37 (0.37, 5.08) | 0.64 | 0.87 |
| Infertility | WHRadjBMI, women | 337 | Women | 0.98 (0.81, 1.19) | 0.86 | | 1.57 (0.68, 3.62) | 0.29 | |
| Lung Cancer | BMI, men | 565 | Men | 1.19 (0.96, 1.47) | 0.11 | 0.16 | 1.25 (1.00, 1.58) | 0.05 | 0.29 |
| Lung Cancer | BMI, women | 565 | Women | 1.48 (1.18, 1.87) | $7.01 \times 10^{-04}$ | | 1.49 (1.19, 1.88) | $6.00 \times 10^{-04}$ | |
| Lung Cancer | WHR, men | 324 | Men | 1.02 (0.74, 1.42) | 0.89 | 0.47 | 0.99 (0.69, 1.42) | 0.95 | 0.40 |
| Lung Cancer | WHR, women | 324 | Women | 1.19 (0.93, 1.54) | 0.17 | | 1.20 (0.93, 1.54) | 0.17 | |
| Lung Cancer | WHRadjBMI, men | 337 | Men | 0.79 (0.57, 1.08) | 0.14 | 0.20 | 0.79 (0.56, 1.12) | 0.19 | 0.24 |
| Lung Cancer | WHRadjBMI, women | 337 | Women | 1.01 (0.81, 1.26) | 0.92 | | 1.01 (0.81, 1.26) | 0.90 | |

| Outcome | GRS | N SNPs | Sex-Strata | OR (95% CI), all available cases and controls for men and women[a] | P, all available cases and controls for men and women[b] | $P_{het}$, all available cases and controls for men and women[c] | OR (95% CI), same number of cases and controls for men and women[a] | P, same number of cases and controls for men and women[b] | $P_{het}$, same number of cases and controls for men and women[c] |
|---|---|---|---|---|---|---|---|---|---|
| NAFLD | BMI, men | 565 | Men | 2.71 (2.08, 3.55) | $2.43\times10^{-13}$ | 0.54 | 2.97 (2.22, 3.96) | $1.75\times10^{-13}$ | 0.86 |
| NAFLD | BMI, women | 565 | Women | 3.07 (2.31, 4.10) | $2.04\times10^{-14}$ | | 3.08 (2.31, 4.10) | $1.96\times10^{-14}$ | |
| NAFLD | WHR, men | 324 | Men | 2.40 (1.58, 3.64) | $4.28\times10^{-05}$ | 0.70 | 2.39 (1.52, 3.75) | $1.68\times10^{-04}$ | 0.71 |
| NAFLD | WHR, women | 324 | Women | 2.66 (1.93, 3.66) | $2.06\times10^{-09}$ | | 2.65 (1.93, 3.65) | $2.16\times10^{-09}$ | |
| NAFLD | WHRadjBMI, men | 337 | Men | 1.31 (0.88, 1.96) | 0.18 | 0.41 | 1.30 (0.84, 2.00) | 0.24 | 0.41 |
| NAFLD | WHRadjBMI, women | 337 | Women | 1.61 (1.22, 2.12) | $7.29\times10^{-04}$ | | 1.61 (1.22, 2.12) | $7.04\times10^{-04}$ | |
| Renal Failure | BMI, men | 565 | Men | 1.61 (1.44, 1.79) | $1.58\times10^{-17}$ | 0.26 | 1.56 (1.37, 1.78) | $3.07\times10^{-11}$ | 0.20 |
| Renal Failure | BMI, women | 565 | Women | 1.77 (1.56, 2.02) | $6.08\times10^{-18}$ | | 1.76 (1.54, 2.00) | $2.28\times10^{-17}$ | |
| Renal Failure | WHR, men | 324 | Men | 1.94 (1.63, 2.30) | $2.88\times10^{-14}$ | $5.32\times10^{-04}$ | 1.99 (1.62, 2.44) | $4.63\times10^{-11}$ | $7.83\times10^{-04}$ |
| Renal Failure | WHR, women | 324 | Women | 1.31 (1.13, 1.51) | $2.95\times10^{-04}$ | | 1.29 (1.12, 1.50) | $4.93\times10^{-04}$ | |
| Renal Failure | WHRadjBMI, men | 337 | Men | 1.31 (1.11, 1.54) | 0.001 | 0.05 | 1.34 (1.11, 1.63) | 0.003 | 0.05 |
| Renal Failure | WHRadjBMI, women | 337 | Women | 1.06 (0.94, 1.20) | 0.34 | | 1.06 (0.94, 1.20) | 0.35 | |
| Renal Failure - Acute | BMI, men | 565 | Men | 1.73 (1.49, 2.01) | $3.40\times10^{-13}$ | 0.32 | 1.75 (1.44, 2.14) | $3.63\times10^{-08}$ | 0.46 |
| Renal Failure - Acute | BMI, women | 565 | Women | 1.97 (1.61, 2.40) | $2.29\times10^{-11}$ | | 1.95 (1.60, 2.38) | $4.06\times10^{-11}$ | |
| Renal Failure - Acute | WHR, men | 324 | Men | 1.87 (1.48, 2.35) | $1.21\times10^{-07}$ | 0.01 | 1.61 (1.18, 2.20) | 0.003 | 0.18 |
| Renal Failure - Acute | WHR, women | 324 | Women | 1.25 (1.00, 1.56) | 0.05 | | 1.24 (0.99, 1.55) | 0.06 | |
| Renal Failure - Acute | WHRadjBMI, men | 337 | Men | 1.18 (0.94, 1.47) | 0.15 | 0.54 | 0.87 (0.65, 1.18) | 0.38 | 0.26 |
| Renal Failure - Acute | WHRadjBMI, women | 337 | Women | 1.07 (0.89, 1.30) | 0.47 | | 1.07 (0.89, 1.30) | 0.47 | |
| Renal Failure - Chronic | BMI, men | 565 | Men | 1.87 (1.59, 2.19) | $2.54\times10^{-14}$ | 0.71 | 2.00 (1.67, 2.40) | $5.25\times10^{-14}$ | 0.33 |
| Renal Failure - Chronic | BMI, women | 565 | Women | 1.78 (1.49, 2.13) | $3.05\times10^{-10}$ | | 1.76 (1.47, 2.11) | $6.35\times10^{-10}$ | |
| Renal Failure - Chronic | WHR, men | 324 | Men | 2.33 (1.82, 3.00) | $3.36\times10^{-11}$ | $1.33\times10^{-04}$ | 2.26 (1.70, 3.00) | $1.65\times10^{-08}$ | $6.46\times10^{-04}$ |
| Renal Failure - Chronic | WHR, women | 324 | Women | 1.25 (1.02, 1.53) | 0.03 | | 1.24 (1.01, 1.51) | 0.04 | |

| Outcome | GRS | N SNPs | Sex-Strata | OR (95% CI), all available cases and controls for men and women[a] | P, all available cases and controls for men and women[b] | $P_{het}$, all available cases and controls for men and women[c] | OR (95% CI), same number of cases and controls for men and women[a] | P, same number of cases and controls for men and women[b] | $P_{het}$, same number of cases and controls for men and women[c] |
|---|---|---|---|---|---|---|---|---|---|
| Renal Failure - Chronic | WHRadjBMI, men | 337 | Men | 1.67 (1.31, 2.12) | $3.01\times10^{-05}$ | $3.55\times10^{-04}$ | 1.63 (1.25, 2.14) | $3.81\times10^{-04}$ | 0.001 |
| Renal Failure - Chronic | WHRadjBMI, women | 337 | Women | 0.97 (0.82, 1.16) | 0.75 | | 0.97 (0.82, 1.15) | 0.73 | |
| Stroke | BMI, men | 565 | Men | 1.42 (1.28, 1.57) | $3.79\times10^{-11}$ | 0.84 | 1.40 (1.23, 1.58) | $9.66\times10^{-08}$ | 0.95 |
| Stroke | BMI, women | 565 | Women | 1.39 (1.23, 1.57) | $8.55\times10^{-08}$ | | 1.40 (1.24, 1.58) | $5.34\times10^{-08}$ | |
| Stroke | WHR, men | 324 | Men | 1.57 (1.33, 1.84) | $5.78\times10^{-08}$ | 0.009 | 1.55 (1.28, 1.87) | $8.55\times10^{-06}$ | 0.02 |
| Stroke | WHR, women | 324 | Women | 1.18 (1.03, 1.35) | 0.02 | | 1.18 (1.03, 1.35) | 0.02 | |
| Stroke | WHRadjBMI, men | 337 | Men | 1.22 (1.05, 1.42) | 0.01 | 0.40 | 1.31 (1.09, 1.58) | 0.004 | 0.15 |
| Stroke | WHRadjBMI, women | 337 | Women | 1.12 (1.00, 1.26) | 0.05 | | 1.12 (0.99, 1.26) | 0.06 | |
| Stroke - Hemorrhagic | BMI, men | 565 | Men | 1.24 (0.95, 1.62) | 0.11 | 0.90 | 1.24 (0.95, 1.62) | 0.11 | 0.86 |
| Stroke - Hemorrhagic | BMI, women | 565 | Women | 1.27 (0.98, 1.64) | 0.07 | | 1.29 (0.99, 1.67) | 0.06 | |
| Stroke - Hemorrhagic | WHR, men | 324 | Men | 1.38 (0.91, 2.09) | 0.13 | 0.54 | 1.38 (0.91, 2.09) | 0.13 | 0.61 |
| Stroke - Hemorrhagic | WHR, women | 324 | Women | 1.18 (0.89, 1.57) | 0.25 | | 1.21 (0.90, 1.63) | 0.20 | |
| Stroke - Hemorrhagic | WHRadjBMI, men | 337 | Men | 1.11 (0.74, 1.64) | 0.62 | 0.99 | 1.11 (0.74, 1.64) | 0.62 | 0.95 |
| Stroke - Hemorrhagic | WHRadjBMI, women | 337 | Women | 1.10 (0.86, 1.41) | 0.44 | | 1.12 (0.87, 1.45) | 0.36 | |
| Stroke - Ischemic | BMI, men | 565 | Men | 1.43 (1.20, 1.70) | $6.55\times10^{-05}$ | 0.56 | 1.55 (1.22, 1.96) | $2.92\times10^{-04}$ | 0.35 |
| Stroke - Ischemic | BMI, women | 565 | Women | 1.31 (1.04, 1.65) | 0.02 | | 1.32 (1.04, 1.67) | 0.02 | |
| Stroke - Ischemic | WHR, men | 324 | Men | 1.33 (1.01, 1.74) | 0.04 | 0.33 | 1.37 (0.95, 1.99) | 0.09 | 0.33 |
| Stroke - Ischemic | WHR, women | 324 | Women | 1.10 (0.85, 1.43) | 0.47 | | 1.10 (0.85, 1.42) | 0.48 | |
| Stroke - Ischemic | WHRadjBMI, men | 337 | Men | 1.27 (0.98, 1.65) | 0.07 | 0.72 | 1.14 (0.80, 1.62) | 0.46 | 0.85 |
| Stroke - Ischemic | WHRadjBMI, women | 337 | Women | 1.19 (0.95, 1.49) | 0.12 | | 1.19 (0.95, 1.49) | 0.14 | |
| Type 1 Diabetes | BMI, men | 565 | Men | 1.46 (1.10, 1.93) | 0.009 | 0.23 | 1.39 (1.02, 1.89) | 0.04 | 0.18 |
| Type 1 Diabetes | BMI, women | 565 | Women | 1.89 (1.39, 2.57) | $5.45\times10^{-05}$ | | 1.87 (1.37, 2.54) | $6.79\times10^{-05}$ | |

| Outcome | GRS | N SNPs | Sex-Strata | OR (95% CI), all available cases and controls for men and women[a] | P, all available cases and controls for men and women[b] | $P_{het}$, all available cases and controls for men and women[c] | OR (95% CI), same number of cases and controls for men and women[a] | P, same number of cases and controls for men and women[b] | $P_{het}$, same number of cases and controls for men and women[c] |
|---|---|---|---|---|---|---|---|---|---|
| Type 1 Diabetes | WHR, men | 324 | Men | 1.15 (0.74, 1.78) | 0.54 | 0.81 | 0.97 (0.60, 1.58) | 0.90 | 0.74 |
| Type 1 Diabetes | WHR, women | 324 | Women | 1.07 (0.76, 1.51) | 0.69 | | 1.07 (0.76, 1.51) | 0.69 | |
| Type 1 Diabetes | WHRadjBMI, men | 337 | Men | 1.29 (0.85, 1.97) | 0.23 | 0.22 | 1.26 (0.79, 2.00) | 0.33 | 0.29 |
| Type 1 Diabetes | WHRadjBMI, women | 337 | Women | 0.94 (0.70, 1.26) | 0.67 | | 0.94 (0.70, 1.26) | 0.67 | |
| Type 2 Diabetes | BMI, men | 565 | Men | 2.73 (2.53, 2.95) | $1.68\times10^{-142}$ | $3.45\times10^{-05}$ | 2.55 (2.30, 2.82) | $6.42\times10^{-72}$ | $5.55\times10^{-06}$ |
| Type 2 Diabetes | BMI, women | 565 | Women | 3.58 (3.23, 3.96) | $2.56\times10^{-134}$ | | 3.56 (3.21, 3.94) | $5.85\times10^{-132}$ | |
| Type 2 Diabetes | WHR, men | 324 | Men | 3.21 (2.85, 3.63) | $8.37\times10^{-80}$ | 0.10 | 3.01 (2.57, 3.53) | $7.11\times10^{-42}$ | 0.04 |
| Type 2 Diabetes | WHR, women | 324 | Women | 3.68 (3.29, 4.12) | $4.10\times10^{-115}$ | | 3.69 (3.29, 4.13) | $5.60\times10^{-114}$ | |
| Type 2 Diabetes | WHRadjBMI, men | 337 | Men | 1.90 (1.69, 2.13) | $1.65\times10^{-27}$ | 0.08 | 1.79 (1.53, 2.08) | $8.14\times10^{-14}$ | 0.04 |
| Type 2 Diabetes | WHRadjBMI, women | 337 | Women | 2.17 (1.97, 2.39) | $1.29\times10^{-55}$ | | 2.16 (1.96, 2.38) | $8.44\times10^{-55}$ | |

BMI, body mass index; CI, confidence interval; CAD, coronary artery disease; COPD, chronic obstructive pulmonary disease; NAFLD, non-alcoholic fatty liver disease; OR, odds ratio; P, P-value.

Logistic regression results for the disease outcomes on the genetic risk scores, using either all available cases for men and women or using the same number of cases and controls in both sexes. In the analyses using the same number of cases and controls, random subsamples of cases and controls were taken from the sex with the most of cases and controls, respectively.

[a]Results per 1-unit higher GRS, corresponding to a predicted 1-SD higher obesity trait, and using the sex-specific estimates approach to construct genetic risk scores

[b]P-value threshold set at <0.001 (=0.05/51) for number of obesity GRS-outcome combinations tested in each analysis

[c]$P_{het}$ from Cochran's Q test for comparisons between male and female estimates for matching traits. $P_{het}$ threshold set at <0.001 (=0.05/48) for number of male-female comparisons in each analysis (fewer Cochran's Q tests since breast cancer was evaluated in women only)

**Table J. Mendelian randomization analyses for the associations between the obesity traits and having been or being a smoker.**

| Exposure | Outcome | GRS | Sex-Strata | OR (95% CI) – Europeans[a] | P – Europeans[b] | $P_{het}$ – Europeans[c] | OR (95% CI) – British[a] | P – British[b] | $P_{het}$ - British[c] |
|---|---|---|---|---|---|---|---|---|---|
| **BMI** | **Having smoked/smoker** | **BMI, combined estimates** | **Combined** | **1.32 (1.29, 1.35)** | **$4.20\times10^{-100}$** | **-** | **1.32 (1.28, 1.35)** | **$4.27\times10^{-89}$** | **-** |
| BMI | Having smoked/smoker | BMI, combined GIANT weights | Combined | 1.28 (1.25, 1.32) | $2.18\times10^{-64}$ | - | 1.28 (1.24, 1.31) | $3.66\times10^{-55}$ | - |
| BMI | Having smoked/smoker | BMI, unweighted | Combined | 1.32 (1.29, 1.36) | $2.89\times10^{-89}$ | - | 1.32 (1.28, 1.36) | $1.40\times10^{-79}$ | - |
| **BMI** | **Having smoked/smoker** | **BMI, male sex-specific estimates** | **Men** | **1.37 (1.32, 1.43)** | **$2.20\times10^{-63}$** | **$8.38\times10^{-04}$** | **1.36 (1.31, 1.42)** | **$1.87\times10^{-54}$** | **0.01** |
| BMI | Having smoked/smoker | BMI, male GIANT weights | Men | 1.32 (1.27, 1.38) | $1.19\times10^{-37}$ | 0.05 | 1.30 (1.25, 1.36) | $8.29\times10^{-31}$ | 0.19 |
| BMI | Having smoked/smoker | BMI, unweighted | Men | 1.41 (1.36, 1.47) | $3.52\times10^{-62}$ | $2.31\times10^{-05}$ | 1.40 (1.34, 1.46) | $2.61\times10^{-54}$ | $4.61\times10^{-04}$ |
| **BMI** | **Having smoked/smoker** | **BMI, female sex-specific estimates** | **Women** | **1.26 (1.22, 1.30)** | **$1.21\times10^{-38}$** | **$8.38\times10^{-04}$** | **1.27 (1.22, 1.32)** | **$4.19\times10^{-36}$** | **0.01** |
| BMI | Having smoked/smoker | BMI, female GIANT weights | Women | 1.25 (1.20, 1.30) | $5.61\times10^{-27}$ | 0.05 | 1.25 (1.20, 1.31) | $3.86\times10^{-24}$ | 0.19 |
| BMI | Having smoked/smoker | BMI, unweighted | Women | 1.25 (1.21, 1.30) | $1.70\times10^{-32}$ | $2.31\times10^{-05}$ | 1.26 (1.21, 1.31) | $4.59\times10^{-30}$ | $4.61\times10^{-04}$ |
| **WHR** | **Having smoked/smoker** | **WHR, combined estimates** | **Combined** | **1.23 (1.19, 1.27)** | **$2.20\times10^{-34}$** | **-** | **1.22 (1.18, 1.26)** | **$1.86\times10^{-29}$** | **-** |
| WHR | Having smoked/smoker | WHR, combined GIANT weights | Combined | 1.26 (1.21, 1.31) | $4.69\times10^{-33}$ | - | 1.25 (1.20, 1.30) | $4.73\times10^{-28}$ | - |
| WHR | Having smoked/smoker | WHR, unweighted | Combined | 1.25 (1.21, 1.30) | $6.44\times10^{-35}$ | - | 1.25 (1.20, 1.30) | $1.03\times10^{-30}$ | - |
| **WHR** | **Having smoked/smoker** | **WHR, male sex-specific estimates** | **Men** | **1.47 (1.38, 1.56)** | **$1.06\times10^{-35}$** | **$2.24\times10^{-14}$** | **1.44 (1.35, 1.53)** | **$1.15\times10^{-29}$** | **$2.41\times10^{-11}$** |
| WHR | Having smoked/smoker | WHR, male GIANT weights | Men | 1.49 (1.38, 1.60) | $4.85\times10^{-26}$ | $3.27\times10^{-09}$ | 1.47 (1.36, 1.58) | $4.52\times10^{-22}$ | $1.38\times10^{-07}$ |
| WHR | Having smoked/smoker | WHR, unweighted | Men | 1.45 (1.35, 1.55) | $5.58\times10^{-26}$ | $6.95\times10^{-08}$ | 1.43 (1.33, 1.53) | $3.71\times10^{-22}$ | $4.60\times10^{-06}$ |
| **WHR** | **Having smoked/smoker** | **WHR, female sex-specific estimates** | **Women** | **1.12 (1.08, 1.16)** | **$3.35\times10^{-09}$** | **$2.24\times10^{-14}$** | **1.12 (1.08, 1.16)** | **$1.12\times10^{-08}$** | **$2.41\times10^{-11}$** |
| WHR | Having smoked/smoker | WHR, female GIANT weights | Women | 1.15 (1.10, 1.20) | $8.37\times10^{-10}$ | $3.27\times10^{-09}$ | 1.15 (1.10, 1.20) | $8.78\times10^{-09}$ | $1.38\times10^{-07}$ |

| Exposure | Outcome | GRS | Sex-Strata | OR (95% CI) – Europeans[a] | P – Europeans[b] | $P_{het}$ – Europeans[c] | OR (95% CI) – British[a] | P – British[b] | $P_{het}$ - British[c] |
|---|---|---|---|---|---|---|---|---|---|
| WHR | Having smoked/smoker | WHR, unweighted | Women | 1.16 (1.11, 1.21) | $1.47\times10^{-12}$ | $6.95\times10^{-08}$ | 1.17 (1.12, 1.22) | $2.94\times10^{-12}$ | $4.60\times10^{-06}$ |
| **WHRadjBMI** | **Having smoked/smoker** | **WHRadjBMI, combined estimates** | **Combined** | **1.03 (1.00, 1.06)** | **0.07** | **-** | **1.03 (1.00, 1.06)** | **0.07** | **-** |
| WHRadjBMI | Having smoked/smoker | WHRadjBMI, combined GIANT weights | Combined | 1.00 (0.97, 1.03) | 0.89 | - | 1.00 (0.96, 1.03) | 0.87 | - |
| WHRadjBMI | Having smoked/smoker | WHRadjBMI, unweighted | Combined | 1.02 (0.99, 1.05) | 0.26 | - | 1.02 (0.99, 1.06) | 0.23 | - |
| **WHRadjBMI** | **Having smoked/smoker** | **WHRadjBMI, male sex-specific estimates** | **Men** | **1.10 (1.04, 1.16)** | **0.001** | **0.006** | **1.10 (1.03, 1.16)** | **0.002** | **0.01** |
| WHRadjBMI | Having smoked/smoker | WHRadjBMI, male GIANT weights | Men | 1.07 (1.00, 1.16) | 0.06 | 0.05 | 1.08 (0.99, 1.17) | 0.07 | 0.06 |
| WHRadjBMI | Having smoked/smoker | WHRadjBMI, unweighted | Men | 1.11 (1.04, 1.18) | 0.003 | 0.005 | 1.11 (1.04, 1.20) | 0.003 | 0.007 |
| **WHRadjBMI** | **Having smoked/smoker** | **WHRadjBMI, female sex-specific estimates** | **Women** | **1.00 (0.97, 1.03)** | **0.85** | **0.006** | **1.01 (0.97, 1.04)** | **0.71** | **0.01** |
| WHRadjBMI | Having smoked/smoker | WHRadjBMI, female GIANT weights | Women | 0.99 (0.95, 1.03) | 0.53 | 0.05 | 0.99 (0.95, 1.03) | 0.58 | 0.06 |
| WHRadjBMI | Having smoked/smoker | WHRadjBMI, unweighted | Women | 0.99 (0.96, 1.03) | 0.74 | 0.005 | 1.00 (0.96, 1.04) | 0.88 | 0.007 |

BMI, body mass index; CI, confidence interval; GRS, genetic risk score; OR, odds ratio; P, P-value; SD, standard deviation; WHR, waist-hip-ratio; WHRadjBMI, waist-hip-ratio adjusted for body mass index.
The GRSs used as the main analyses in bold.
[a]Estimates per 1-SD higher obesity trait and with results both for all Europeans and for British ancestry only subset. Participants were denoted as "British" if they were in the British ancestry subset as defined by the UK Biobank (14) (based on self-report of British ancestry and similar ancestry according to principal components analysis)
[b]P-value threshold set to <0.003 (=0.05/15) for 15 obesity trait-risk factor combination, including fasting glucose, fasting insulin, diastolic and systolic blood pressure
[c]$P_{het}$-value from Cochran's Q test for heterogeneity between male and female estimates for matching traits and SNP-selection and weighting approaches. $P_{het}$ threshold set to <0.003 (=0.05/15) for 15 obesity trait-risk factor combination, including fasting glucose, fasting insulin, diastolic and systolic blood pressure

**Table K. Mendelian randomization analyses with blood pressure using different SNP-selection and weighting approaches, unadjusted for smoking.**

| Exposure | Outcome | GRS | Sex-strata | Estimate, Clin (95% CI)[a] | P (Clin)[b] | $P_{het}$ (Clin)[c] | Estimate, SD (95% CI)[a] | P (SD)[b] | $P_{het}$ (SD)[c] |
|---|---|---|---|---|---|---|---|---|---|
| **BMI** | **DBP** | **BMI, combined estimates** | **Combined** | **2.92 (2.78,3.07)** | **$<1\times10^{-200}$** | **-** | **0.27 (0.26,0.28)** | **$<1\times10^{-200}$** | **-** |
| BMI | DBP | BMI, combined GIANT weights | Combined | 2.67 (2.51,2.83) | $<1\times10^{-200}$ | - | 0.25 (0.23,0.26) | $<1\times10^{-200}$ | - |
| BMI | DBP | BMI, unweighted | Combined | 3.09 (2.94,3.24) | $<1\times10^{-200}$ | - | 0.29 (0.27,0.30) | $<1\times10^{-200}$ | - |
| **BMI** | **DBP** | **BMI, male sex-specific estimates** | **Men** | **2.60 (2.39,2.80)** | **$1.30\times10^{-136}$** | **$1.83\times10^{-05}$** | **0.24 (0.22,0.26)** | **$1.58\times10^{-140}$** | **$5.24\times10^{-05}$** |
| BMI | DBP | BMI, male giant weights | Men | 2.31 (2.07,2.54) | $1.01\times10^{-81}$ | $6.79\times10^{-05}$ | 0.22 (0.19,0.24) | $2.10\times10^{-84}$ | $2.08\times10^{-04}$ |
| BMI | DBP | BMI, unweighted | Men | 2.77 (2.54,2.99) | $8.37\times10^{-128}$ | $2.40\times10^{-04}$ | 0.26 (0.24,0.28) | $1.50\times10^{-131}$ | $5.62\times10^{-04}$ |
| **BMI** | **DBP** | **BMI, female sex-specific estimates** | **Women** | **3.22 (3.02,3.41)** | **$<1\times10^{-200}$** | **$1.83\times10^{-05}$** | **0.30 (0.28,0.31)** | **$<1\times10^{-200}$** | **$5.24\times10^{-05}$** |
| BMI | DBP | BMI, female GIANT weights | Women | 2.97 (2.74,3.19) | $4.40\times10^{-149}$ | $6.79\times10^{-05}$ | 0.27 (0.25,0.29) | $6.15\times10^{-149}$ | $2.08\times10^{-04}$ |
| BMI | DBP | BMI, unweighted | Women | 3.35 (3.13,3.56) | $<1\times10^{-200}$ | $2.40\times10^{-04}$ | 0.31 (0.29,0.33) | $<1\times10^{-200}$ | $5.62\times10^{-04}$ |
| **BMI** | **SBP** | **BMI, combined estimates** | **Combined** | **3.52 (3.28,3.77)** | **$5.67\times10^{-176}$** | **-** | **0.19 (0.18,0.20)** | **$1.52\times10^{-184}$** | **-** |
| BMI | SBP | BMI, combined GIANT weights | Combined | 3.14 (2.87,3.42) | $3.94\times10^{-113}$ | - | 0.17 (0.16,0.18) | $4.78\times10^{-118}$ | - |
| BMI | SBP | BMI, unweighted | Combined | 3.73 (3.47,3.99) | $2.59\times10^{-170}$ | - | 0.20 (0.19,0.22) | $8.32\times10^{-179}$ | - |
| **BMI** | **SBP** | **BMI, male sex-specific estimates** | **Men** | **3.41 (3.07,3.75)** | **$1.13\times10^{-85}$** | **0.42** | **0.19 (0.17,0.21)** | **$2.03\times10^{-90}$** | **0.99** |
| BMI | SBP | BMI, male giant weights | Men | 2.99 (2.59,3.38) | $4.14\times10^{-50}$ | 0.38 | 0.17 (0.15,0.19) | $1.26\times10^{-52}$ | 0.82 |
| BMI | SBP | BMI, unweighted | Men | 3.74 (3.36,4.11) | $5.42\times10^{-85}$ | 0.84 | 0.21 (0.19,0.23) | $2.05\times10^{-89}$ | 0.32 |
| **BMI** | **SBP** | **BMI, female sex-specific estimates** | **Women** | **3.61 (3.27,3.95)** | **$4.07\times10^{-97}$** | **0.42** | **0.19 (0.17,0.21)** | **$5.69\times10^{-101}$** | **0.99** |
| BMI | SBP | BMI, female GIANT weights | Women | 3.23 (2.85,3.62) | $2.87\times10^{-60}$ | 0.38 | 0.17 (0.15,0.19) | $3.58\times10^{-62}$ | 0.82 |
| BMI | SBP | BMI, unweighted | Women | 3.69 (3.32,4.05) | $8.62\times10^{-88}$ | 0.84 | 0.20 (0.18,0.21) | $2.73\times10^{-91}$ | 0.32 |
| **WHR** | **DBP** | **WHR, combined estimates** | **Combined** | **2.68 (2.49,2.86)** | **$1.40\times10^{-171}$** | **-** | **0.25 (0.23,0.26)** | **$6.13\times10^{-174}$** | **-** |
| WHR | DBP | WHR, combined GIANT weights | Combined | 2.54 (2.32,2.75) | $4.44\times10^{-119}$ | - | 0.23 (0.21,0.25) | $2.24\times10^{-120}$ | - |
| WHR | DBP | WHR, unweighted | Combined | 2.83 (2.62,3.03) | $3.17\times10^{-163}$ | - | 0.26 (0.24,0.28) | $8.30\times10^{-166}$ | - |
| **WHR** | **DBP** | **WHR, male sex-specific estimates** | **Men** | **2.80 (2.47,3.12)** | **$2.53\times10^{-63}$** | **0.01** | **0.26 (0.23,0.29)** | **$1.53\times10^{-64}$** | **0.01** |
| WHR | DBP | WHR, male GIANT weights | Men | 2.84 (2.44,3.24) | $2.03\times10^{-43}$ | 0.08 | 0.26 (0.23,0.30) | $2.98\times10^{-44}$ | 0.07 |

| Exposure | Outcome | GRS | Sex-strata | Estimate, Clin (95% CI)[a] | P (Clin)[b] | $P_{het}$ (Clin)[c] | Estimate, SD (95% CI)[a] | P (SD)[b] | $P_{het}$ (SD)[c] |
|---|---|---|---|---|---|---|---|---|---|
| WHR | DBP | WHR, unweighted | Men | 2.64 (2.27,3.01) | $5.69\times10^{-44}$ | 0.98 | 0.25 (0.21,0.28) | $7.48\times10^{-45}$ | 0.96 |
| **WHR** | **DBP** | **WHR, female sex-specific estimates** | **Women** | **2.30 (2.10,2.51)** | $\mathbf{3.95\times10^{-108}}$ | **0.01** | **0.21 (0.19,0.23)** | $\mathbf{8.97\times10^{-110}}$ | **0.01** |
| WHR | DBP | WHR, female GIANT weights | Women | 2.42 (2.17,2.67) | $8.79\times10^{-81}$ | 0.08 | 0.22 (0.20,0.25) | $1.19\times10^{-81}$ | 0.07 |
| WHR | DBP | WHR, unweighted | Women | 2.63 (2.40,2.87) | $4.21\times10^{-109}$ | 0.98 | 0.24 (0.22,0.27) | $4.86\times10^{-111}$ | 0.96 |
| **WHR** | **SBP** | **WHR, combined estimates** | **Combined** | **3.98 (3.66,4.30)** | $\mathbf{2.81\times10^{-129}}$ | **-** | **0.22 (0.20,0.23)** | $\mathbf{2.56\times10^{-134}}$ | **-** |
| WHR | SBP | WHR, combined GIANT weights | Combined | 3.89 (3.52,4.26) | $7.08\times10^{-95}$ | - | 0.21 (0.19,0.23) | $8.26\times10^{-99}$ | - |
| WHR | SBP | WHR, unweighted | Combined | 4.33 (3.98,4.68) | $5.60\times10^{-130}$ | - | 0.23 (0.22,0.25) | $1.32\times10^{-134}$ | - |
| **WHR** | **SBP** | **WHR, male sex-specific estimates** | **Men** | **3.61 (3.07,4.15)** | $\mathbf{4.45\times10^{-39}}$ | **0.94** | **0.20 (0.17,0.23)** | $\mathbf{1.63\times10^{-40}}$ | **0.55** |
| WHR | SBP | WHR, male GIANT weights | Men | 3.96 (3.29,4.63) | $4.71\times10^{-31}$ | 0.93 | 0.22 (0.18,0.26) | $3.80\times10^{-32}$ | 0.61 |
| WHR | SBP | WHR, unweighted | Men | 3.84 (3.22,4.46) | $7.63\times10^{-34}$ | 0.65 | 0.21 (0.18,0.25) | $6.75\times10^{-35}$ | 0.99 |
| **WHR** | **SBP** | **WHR, female sex-specific estimates** | **Women** | **3.59 (3.23,3.95)** | $\mathbf{2.43\times10^{-85}}$ | **0.94** | **0.19 (0.17,0.21)** | $\mathbf{5.02\times10^{-89}}$ | **0.55** |
| WHR | SBP | WHR, female GIANT weights | Women | 3.93 (3.49,4.36) | $5.35\times10^{-69}$ | 0.93 | 0.21 (0.19,0.23) | $2.16\times10^{-72}$ | 0.61 |
| WHR | SBP | WHR, unweighted | Women | 4.01 (3.61,4.42) | $7.00\times10^{-83}$ | 0.65 | 0.21 (0.19,0.23) | $2.72\times10^{-86}$ | 0.99 |
| **WHRadjBMI** | **DBP** | **WHRadjBMI, combined estimates** | **Combined** | **1.07 (0.91,1.23)** | $\mathbf{1.87\times10^{-38}}$ | **-** | **0.10 (0.08,0.11)** | $\mathbf{3.24\times10^{-39}}$ | **-** |
| WHRadjBMI | DBP | WHRadjBMI, combined GIANT weights | Combined | 0.92 (0.73,1.10) | $1.60\times10^{-22}$ | - | 0.09 (0.07,0.10) | $4.57\times10^{-23}$ | - |
| WHRadjBMI | DBP | WHRadjBMI, unweighted | Combined | 0.86 (0.68,1.04) | $1.29\times10^{-20}$ | - | 0.08 (0.06,0.10) | $2.66\times10^{-21}$ | - |
| **WHRadjBMI** | **DBP** | **WHRadjBMI, male sex-specific estimates** | **Men** | **0.96 (0.66,1.26)** | $\mathbf{3.50\times10^{-10}}$ | **0.75** | **0.09 (0.06,0.12)** | $\mathbf{2.51\times10^{-10}}$ | **0.74** |
| WHRadjBMI | DBP | WHRadjBMI, male GIANT weights | Men | 1.21 (0.81,1.61) | $2.06\times10^{-09}$ | 0.36 | 0.11 (0.07,0.15) | $1.93\times10^{-09}$ | 0.39 |
| WHRadjBMI | DBP | WHRadjBMI, unweighted | Men | 0.77 (0.41,1.13) | $2.26\times10^{-05}$ | 0.90 | 0.07 (0.04,0.10) | $1.67\times10^{-05}$ | 0.89 |
| **WHRadjBMI** | **DBP** | **WHRadjBMI, female sex-specific estimates** | **Women** | **1.02 (0.84,1.19)** | $\mathbf{2.36\times10^{-30}}$ | **0.75** | **0.09 (0.08,0.11)** | $\mathbf{6.41\times10^{-31}}$ | **0.74** |
| WHRadjBMI | DBP | WHRadjBMI, female GIANT weights | Women | 1.00 (0.79,1.21) | $5.14\times10^{-20}$ | 0.36 | 0.09 (0.07,0.11) | $1.63\times10^{-20}$ | 0.39 |
| WHRadjBMI | DBP | WHRadjBMI, unweighted | Women | 0.80 (0.59,1.00) | $8.91\times10^{-15}$ | 0.90 | 0.07 (0.06,0.09) | $2.65\times10^{-15}$ | 0.89 |

| Exposure | Outcome | GRS | Sex-strata | Estimate, Clin (95% CI)[a] | P (Clin)[b] | $P_{het}$ (Clin)[c] | Estimate, SD (95% CI)[a] | P (SD)[b] | $P_{het}$ (SD)[c] |
|---|---|---|---|---|---|---|---|---|---|
| **WHRadjBMI** | **SBP** | **WHRadjBMI, combined estimates** | **Combined** | **2.27 (1.99,2.55)** | **$7.50\times10^{-57}$** | **-** | **0.12 (0.11,0.14)** | **$2.97\times10^{-59}$** | **-** |
| WHRadjBMI | SBP | WHRadjBMI, combined GIANT weights | Combined | 2.13 (1.81,2.45) | $3.69\times10^{-39}$ | - | 0.12 (0.10,0.13) | $7.48\times10^{-41}$ | - |
| WHRadjBMI | SBP | WHRadjBMI, unweighted | Combined | 2.25 (1.94,2.56) | $3.43\times10^{-45}$ | - | 0.12 (0.11,0.14) | $3.21\times10^{-47}$ | - |
| **WHRadjBMI** | **SBP** | **WHRadjBMI, male sex-specific estimates** | **Men** | **1.58 (1.07,2.08)** | **$8.41\times10^{-10}$** | **0.04** | **0.09 (0.06,0.12)** | **$4.38\times10^{-10}$** | **0.08** |
| WHRadjBMI | SBP | WHRadjBMI, male GIANT weights | Men | 2.33 (1.67,3.00) | $6.20\times10^{-12}$ | 1.00 | 0.13 (0.09,0.17) | $3.59\times10^{-12}$ | 0.86 |
| WHRadjBMI | SBP | WHRadjBMI, unweighted | Men | 2.12 (1.52,2.72) | $3.87\times10^{-12}$ | 0.84 | 0.12 (0.09,0.15) | $1.62\times10^{-12}$ | 0.65 |
| **WHRadjBMI** | **SBP** | **WHRadjBMI, female sex-specific estimates** | **Women** | **2.18 (1.87,2.49)** | **$7.24\times10^{-44}$** | **0.04** | **0.12 (0.10,0.13)** | **$3.69\times10^{-46}$** | **0.08** |
| WHRadjBMI | SBP | WHRadjBMI, female GIANT weights | Women | 2.33 (1.96,2.71) | $1.33\times10^{-33}$ | 1.00 | 0.13 (0.11,0.14) | $1.30\times10^{-35}$ | 0.86 |
| WHRadjBMI | SBP | WHRadjBMI, unweighted | Women | 2.05 (1.70,2.41) | $1.14\times10^{-29}$ | 0.84 | 0.11 (0.09,0.13) | $3.81\times10^{-31}$ | 0.65 |

BMI, body mass index; CI, confidence interval; Clin, clinical; DBP, diastolic blood pressure; GRS, genetic risk score; P, P-value; SBP, systolic blood pressure; SD, standard deviation; WHR, waist-hip-ratio; WHRadjBMI, waist-hip-ratio adjusted for body mass index.

The GRSs considered the main approach, using the sex-specific estimates approach, in bold.

[a]Estimates per 1-SD higher risk factor, with outcome either in clinical units in mmHg or in SD units

[b]P-value threshold set at <0.003 (=0.05/15) for 15 obesity trait-risk factor combinations, including DBP, SBP, fasting glucose, fasting insulin and smoking status

[c]$P_{het}$-values from Cochran's Q test given for comparisons between male and female estimates for matching traits and units. $P_{het}$-threshold set at <0.003 (=0.05/15) for 15 obesity trait-risk factor combinations, including DBP, SBP, fasting glucose, fasting insulin and smoking status

**Table L. Mendelian randomization analyses with blood pressure, unadjusted for smoking in British ancestry only subset.**

| Exposure | Outcome | Sex-strata | Estimate, Clin (95% CI)[a] | P (Clin)[b] | $P_{het}$ (Clin)[c] | Estimate, SD (95% CI)[a] | P (SD)[b] | $P_{het}$ (SD)[c] |
|---|---|---|---|---|---|---|---|---|
| BMI | DBP | Combined | 2.91 (2.76,3.06) | $<1\times10^{-200}$ | - | 0.27 (0.26,0.28) | $<1\times10^{-200}$ | - |
| BMI | DBP | Men | 2.59 (2.38,2.81) | $2.44\times10^{-123}$ | $4.63\times10^{-05}$ | 0.24 (0.22,0.26) | $4.88\times10^{-127}$ | $1.24\times10^{-04}$ |
| BMI | DBP | Women | 3.21 (3.01,3.42) | $3.18\times10^{-200}$ | | 0.30 (0.28,0.31) | $<1\times10^{-200}$ | |
| BMI | SBP | Combined | 3.51 (3.25,3.77) | $4.26\times10^{-155}$ | - | 0.19 (0.18,0.20) | $8.29\times10^{-163}$ | - |
| BMI | SBP | Men | 3.40 (3.04,3.75) | $8.05\times10^{-77}$ | 0.45 | 0.19 (0.17,0.21) | $3.36\times10^{-81}$ | 0.97 |
| BMI | SBP | Women | 3.59 (3.23,3.95) | $1.69\times10^{-84}$ | | 0.19 (0.17,0.21) | $8.35\times10^{-88}$ | |
| WHR | DBP | Combined | 2.66 (2.46,2.86) | $2.45\times10^{-152}$ | - | 0.25 (0.23,0.26) | $2.79\times10^{-154}$ | - |
| WHR | DBP | Men | 2.82 (2.48,3.16) | $1.56\times10^{-58}$ | 0.009 | 0.26 (0.23,0.29) | $8.56\times10^{-60}$ | 0.007 |
| WHR | DBP | Women | 2.28 (2.06,2.49) | $5.07\times10^{-94}$ | | 0.21 (0.19,0.23) | $3.00\times10^{-95}$ | |
| WHR | SBP | Combined | 3.99 (3.65,4.33) | $5.35\times10^{-116}$ | - | 0.22 (0.20,0.23) | $1.76\times10^{-120}$ | - |
| WHR | SBP | Men | 3.64 (3.07,4.21) | $4.84\times10^{-36}$ | 0.87 | 0.20 (0.17,0.23) | $2.31\times10^{-37}$ | 0.52 |
| WHR | SBP | Women | 3.59 (3.21,3.97) | $1.43\times10^{-75}$ | | 0.19 (0.17,0.21) | $5.63\times10^{-79}$ | |
| WHRadjBMI | DBP | Combined | 1.06 (0.89,1.23) | $2.07\times10^{-34}$ | - | 0.10 (0.08,0.11) | $6.77\times10^{-35}$ | - |
| WHRadjBMI | DBP | Men | 0.98 (0.66,1.30) | $1.20\times10^{-09}$ | 0.94 | 0.09 (0.06,0.12) | $9.88\times10^{-10}$ | 0.94 |
| WHRadjBMI | DBP | Women | 0.99 (0.81,1.18) | $2.85\times10^{-26}$ | | 0.09 (0.07,0.11) | $1.19\times10^{-26}$ | |
| WHRadjBMI | SBP | Combined | 2.27 (1.97,2.57) | $2.76\times10^{-51}$ | - | 0.12 (0.11,0.14) | $2.63\times10^{-53}$ | - |
| WHRadjBMI | SBP | Men | 1.62 (1.09,2.15) | $2.28\times10^{-09}$ | 0.09 | 0.09 (0.06,0.12) | $1.30\times10^{-09}$ | 0.14 |
| WHRadjBMI | SBP | Women | 2.16 (1.84,2.49) | $6.30\times10^{-39}$ | | 0.12 (0.10,0.13) | $5.93\times10^{-41}$ | |

BMI, body mass index; CI, confidence interval; Clin, clinical; DBP, diastolic blood pressure; GRS, genetic risk score; P, P-value; SBP, systolic blood pressure; SD, standard deviation; WHR, waist-hip-ratio; WHRadjBMI, waist-hip-ratio adjusted for body mass index.
Participants were denoted as "British" if they were in the British ancestry subset as defined by the UK Biobank (14) (based on self-report of British ancestry and similar ancestry according to principal components analysis).
[a]Estimates per 1-SD higher risk factor, with outcome either in clinical units in mmHg or in SD units and using the sex-specific estimates approach to construct genetic risk scores
[b]P-value threshold set at <0.003 (= 0.05/15) for 15 obesity trait-outcome combinations, including SBP, DBP, fasting glucose, fasting insulin, and smoking
[c]$P_{het}$-value from Cochran's Q test given for comparisons between male and female estimates for matching traits and units. $P_{het}$-threshold set at <0.003 (= 0.05/15) for 15 obesity trait-outcome combinations, including SBP, DBP, fasting glucose, fasting insulin, and smoking

**Table M. Mendelian randomization analyses for blood pressure traits, unadjusted and adjusted for smoking status.**

| Exposure | Outcome | Sex-strata | Estimate, unadj (95% CI)[a] | P, unadj[b] | $P_{het}$, unadj[c] | Estimate, adj (95% CI)[a] | P, adj[b] | $P_{het}$, adj[c] | % Change[d] |
|---|---|---|---|---|---|---|---|---|---|
| BMI | DBP | Combined | 0.27 (0.26,0.28) | $<1\times10^{-200}$ | - | 0.27 (0.26,0.29) | $<1\times10^{-200}$ | - | 0 |
| BMI | DBP | Men | 0.24 (0.22,0.26) | $4.88\times10^{-127}$ | $1.24\times10^{-04}$ | 0.24 (0.22,0.26) | $1.81\times10^{-137}$ | $2.49\times10^{-05}$ | 0 |
| BMI | DBP | Women | 0.30 (0.28,0.31) | $<1\times10^{-200}$ | | 0.30 (0.28,0.32) | $<1\times10^{-200}$ | | 0 |
| BMI | SBP | Combined | 0.19 (0.18,0.20) | $8.29\times10^{-163}$ | - | 0.19 (0.18,0.21) | $8.51\times10^{-186}$ | - | 0 |
| BMI | SBP | Men | 0.19 (0.17,0.21) | $3.36\times10^{-81}$ | 0.97 | 0.19 (0.17,0.21) | $1.14\times10^{-86}$ | 0.68 | 0 |
| BMI | SBP | Women | 0.19 (0.17,0.21) | $8.35\times10^{-88}$ | | 0.19 (0.18,0.21) | $5.56\times10^{-104}$ | | 0 |
| WHR | DBP | Combined | 0.25 (0.23,0.26) | $2.79\times10^{-154}$ | - | 0.25 (0.23,0.27) | $7.12\times10^{-174}$ | - | 0 |
| WHR | DBP | Men | 0.26 (0.23,0.29) | $8.56\times10^{-60}$ | 0.007 | 0.26 (0.23,0.29) | $6.08\times10^{-63}$ | 0.01 | 0 |
| WHR | DBP | Women | 0.21 (0.19,0.23) | $3.00\times10^{-95}$ | | 0.21 (0.20,0.23) | $1.82\times10^{-110}$ | | 0 |
| WHR | SBP | Combined | 0.22 (0.20,0.23) | $1.76\times10^{-120}$ | - | 0.22 (0.20,0.24) | $7.81\times10^{-135}$ | - | 0 |
| WHR | SBP | Men | 0.20 (0.17,0.23) | $2.31\times10^{-37}$ | 0.52 | 0.20 (0.17,0.23) | $1.24\times10^{-38}$ | 0.68 | 0 |
| WHR | SBP | Women | 0.19 (0.17,0.21) | $5.63\times10^{-79}$ | | 0.19 (0.17,0.21) | $2.97\times10^{-90}$ | | 0 |
| WHRadjBMI | DBP | Combined | 0.10 (0.08,0.11) | $6.77\times10^{-35}$ | - | 0.10 (0.08,0.11) | $3.24\times10^{-39}$ | - | 0 |
| WHRadjBMI | DBP | Men | 0.09 (0.06,0.12) | $9.88\times10^{-10}$ | 0.94 | 0.09 (0.06,0.12) | $3.50\times10^{-10}$ | 0.72 | 0 |
| WHRadjBMI | DBP | Women | 0.09 (0.07,0.11) | $1.19\times10^{-26}$ | | 0.09 (0.08,0.11) | $6.09\times10^{-31}$ | | 0 |
| WHRadjBMI | SBP | Combined | 0.12 (0.11,0.14) | $2.63\times10^{-53}$ | - | 0.12 (0.11,0.14) | $2.60\times10^{-59}$ | - | 0 |
| WHRadjBMI | SBP | Men | 0.09 (0.06,0.12) | $1.30\times10^{-09}$ | 0.14 | 0.09 (0.06,0.11) | $7.12\times10^{-10}$ | 0.07 | 0 |
| WHRadjBMI | SBP | Women | 0.12 (0.10,0.13) | $5.93\times10^{-41}$ | | 0.12 (0.10,0.13) | $3.13\times10^{-46}$ | | 0 |

Adj, adjusted for smoking status; BMI, body mass index; CI, confidence interval; DBP, diastolic blood pressure; GRS, genetic risk score; P, P-value; SBP, systolic blood pressure; SD, standard deviation; unadj, unadjusted for smoking status.

[a]Estimates per 1-SD higher obesity trait, with outcome in SD units, for the sex-specific estimates approach to construct genetic risk scores

[b]P-value threshold set at <0.003 (=0.05/15) for 15 obesity trait-risk factor combinations, including fasting glucose, fasting insulin and smoking status as well as DBP and SBP

[c]$P_{het}$ from Cochran's Q test given for comparisons between male and female estimates for matching traits. $P_{het}$-threshold set at <0.003 (=0.05/15) for 15 obesity trait-risk factor combinations, including fasting glucose, fasting insulin and smoking status as well as DBP and SBP

[d]% Change is the relative change for the adjusted for smoking point estimate compared to the unadjusted for smoking status point estimate

**Table N. Two-sample Mendelian randomization analyses for fasting insulin and fasting glucose.**

| Exposure | Outcome | Sex-Strata | N SNPs available (N SNPs GRS)[a] | Outcome Unit | Method | Estimate (95% CI)[b] | P[c] | $P_{het}$[d] |
|---|---|---|---|---|---|---|---|---|
| BMI | FG | Combined | 482 (565) | mmol/L | IVW | 0.075 (0.054,0.095) | $6.72\times10^{-13}$ | - |
| BMI | FG | Combined | 482 (565) | mmol/L | MR-Egger | 0.102 (0.049,0.156) | $1.85\times10^{-04}$ | - |
| BMI | FG | Combined | 482 (565) | mmol/L | MR-Egger (intercept) | -0.000 (-0.001,0.000) | 0.27 | - |
| BMI | FG | Combined | 482 (565) | mmol/L | Weighted median | 0.078 (0.050,0.107) | $7.71\times10^{-08}$ | - |
| BMI | FG | Men | 482 (565) | mmol/L | IVW | 0.066 (0.039,0.092) | $1.51\times10^{-06}$ | 0.42 |
| BMI | FG | Men | 482 (565) | mmol/L | MR-Egger | 0.060 (-0.003,0.124) | 0.06 | 0.11 |
| BMI | FG | Men | 482 (565) | mmol/L | MR-Egger (intercept) | 0.000 (-0.001,0.001) | 0.86 | 0.16 |
| BMI | FG | Men | 482 (565) | mmol/L | Weighted median | 0.074 (0.025,0.123) | 0.003 | 0.26 |
| BMI | FG | Women | 482 (565) | mmol/L | IVW | 0.081 (0.056,0.106) | $3.51\times10^{-10}$ | 0.42 |
| BMI | FG | Women | 482 (565) | mmol/L | MR-Egger | 0.136 (0.069,0.202) | $5.95\times10^{-05}$ | 0.11 |
| BMI | FG | Women | 482 (565) | mmol/L | MR-Egger (intercept) | -0.001 (-0.002,0.000) | 0.08 | 0.16 |
| BMI | FG | Women | 482 (565) | mmol/L | Weighted median | 0.109 (0.073,0.145) | $3.20\times10^{-09}$ | 0.26 |
| BMI | FI | Combined | 482 (565) | ln(pmol/L) | IVW | 0.214 (0.191,0.236) | $2.25\times10^{-78}$ | - |
| BMI | FI | Combined | 482 (565) | ln(pmol/L) | MR-Egger | 0.214 (0.155,0.273) | $1.17\times10^{-12}$ | - |
| BMI | FI | Combined | 482 (565) | ln(pmol/L) | MR-Egger (intercept) | -0.000 (-0.001,0.001) | 0.99 | - |
| BMI | FI | Combined | 482 (565) | ln(pmol/L) | Weighted median | 0.222 (0.184,0.260) | $7.89\times10^{-31}$ | - |
| BMI | FI | Men | 482 (565) | ln(pmol/L) | IVW | 0.222 (0.191,0.253) | $2.94\times10^{-44}$ | 0.50 |
| BMI | FI | Men | 482 (565) | ln(pmol/L) | MR-Egger | 0.253 (0.179,0.326) | $1.67\times10^{-11}$ | 0.21 |
| BMI | FI | Men | 482 (565) | ln(pmol/L) | MR-Egger (intercept) | -0.001 (-0.002,0.001) | 0.36 | 0.29 |
| BMI | FI | Men | 482 (565) | ln(pmol/L) | Weighted median | 0.221 (0.166,0.276) | $2.52\times10^{-15}$ | 0.89 |
| BMI | FI | Women | 482 (565) | ln(pmol/L) | IVW | 0.207 (0.178,0.236) | $2.36\times10^{-44}$ | 0.50 |
| BMI | FI | Women | 482 (565) | ln(pmol/L) | MR-Egger | 0.185 (0.109,0.261) | $1.96\times10^{-06}$ | 0.21 |
| BMI | FI | Women | 482 (565) | ln(pmol/L) | MR-Egger (intercept) | 0.000 (-0.001,0.002) | 0.55 | 0.29 |
| BMI | FI | Women | 482 (565) | ln(pmol/L) | Weighted median | 0.216 (0.167,0.264) | $2.77\times10^{-18}$ | 0.89 |
| WHR | FG | Combined | 261 (324) | mmol/L | IVW | 0.078 (0.050,0.106) | $4.54\times10^{-08}$ | - |
| WHR | FG | Combined | 261 (324) | mmol/L | MR-Egger | 0.077 (-0.015,0.169) | 0.10 | - |

| Exposure | Outcome | Sex-Strata | N SNPs available (N SNPs GRS)[a] | Outcome Unit | Method | Estimate (95% CI)[b] | P[c] | $P_{het}$[d] |
|---|---|---|---|---|---|---|---|---|
| WHR | FG | Combined | 261 (324) | mmol/L | MR-Egger (intercept) | 0.000 (-0.001,0.001) | 0.98 | - |
| WHR | FG | Combined | 261 (324) | mmol/L | Weighted median | 0.075 (0.038,0.113) | $7.26\times10^{-05}$ | - |
| WHR | FG | Men | 261 (324) | mmol/L | IVW | 0.061 (0.019,0.102) | 0.004 | 0.41 |
| WHR | FG | Men | 261 (324) | mmol/L | MR-Egger | 0.083 (-0.006,0.172) | 0.07 | 0.84 |
| WHR | FG | Men | 261 (324) | mmol/L | MR-Egger (intercept) | -0.000 (-0.002,0.001) | 0.58 | 0.93 |
| WHR | FG | Men | 261 (324) | mmol/L | Weighted median | 0.080 (0.015,0.145) | 0.02 | 0.88 |
| WHR | FG | Women | 261 (324) | mmol/L | IVW | 0.082 (0.053,0.111) | $4.21\times10^{-08}$ | 0.41 |
| WHR | FG | Women | 261 (324) | mmol/L | MR-Egger | 0.095 (0.023,0.167) | 0.01 | 0.84 |
| WHR | FG | Women | 261 (324) | mmol/L | MR-Egger (intercept) | -0.000 (-0.002,0.001) | 0.70 | 0.93 |
| WHR | FG | Women | 261 (324) | mmol/L | Weighted median | 0.086 (0.044,0.128) | $5.77\times10^{-05}$ | 0.88 |
| WHR | FI | Combined | 261 (324) | ln(pmol/L) | IVW | 0.217 (0.179,0.254) | $3.46\times10^{-29}$ | - |
| WHR | FI | Combined | 261 (324) | ln(pmol/L) | MR-Egger | 0.212 (0.089,0.336) | $7.53\times10^{-04}$ | - |
| WHR | FI | Combined | 261 (324) | ln(pmol/L) | MR-Egger (intercept) | 0.000 (-0.002,0.002) | 0.94 | - |
| WHR | FI | Combined | 261 (324) | ln(pmol/L) | Weighted median | 0.223 (0.176,0.271) | $3.17\times10^{-20}$ | - |
| WHR | FI | Men | 261 (324) | ln(pmol/L) | IVW | 0.195 (0.137,0.253) | $3.65\times10^{-11}$ | 0.82 |
| WHR | FI | Men | 261 (324) | ln(pmol/L) | MR-Egger | 0.200 (0.075,0.324) | 0.002 | 0.90 |
| WHR | FI | Men | 261 (324) | ln(pmol/L) | MR-Egger (intercept) | -0.000 (-0.002,0.002) | 0.93 | 0.96 |
| WHR | FI | Men | 261 (324) | ln(pmol/L) | Weighted median | 0.207 (0.130,0.285) | $1.56\times10^{-07}$ | 0.95 |
| WHR | FI | Women | 261 (324) | ln(pmol/L) | IVW | 0.203 (0.165,0.241) | $4.32\times10^{-26}$ | 0.82 |
| WHR | FI | Women | 261 (324) | ln(pmol/L) | MR-Egger | 0.210 (0.117,0.302) | $8.76\times10^{-06}$ | 0.90 |
| WHR | FI | Women | 261 (324) | ln(pmol/L) | MR-Egger (intercept) | -0.000 (-0.002,0.002) | 0.88 | 0.96 |
| WHR | FI | Women | 261 (324) | ln(pmol/L) | Weighted median | 0.205 (0.151,0.258) | $6.51\times10^{-14}$ | 0.95 |
| WHRadjBMI | FG | Combined | 267 (337) | mmol/L | IVW | 0.046 (0.019,0.073) | $8.38\times10^{-04}$ | - |
| WHRadjBMI | FG | Combined | 267 (337) | mmol/L | MR-Egger | 0.053 (-0.022,0.128) | 0.17 | - |
| WHRadjBMI | FG | Combined | 267 (337) | mmol/L | MR-Egger (intercept) | -0.000 (-0.001,0.001) | 0.84 | - |
| WHRadjBMI | FG | Combined | 267 (337) | mmol/L | Weighted median | 0.040 (0.007,0.074) | 0.02 | - |
| WHRadjBMI | FG | Men | 267 (337) | mmol/L | IVW | 0.028 (-0.017,0.074) | 0.22 | 0.46 |

| Exposure | Outcome | Sex-Strata | N SNPs available (N SNPs GRS)[a] | Outcome Unit | Method | Estimate (95% CI)[b] | P[c] | $P_{het}$[d] |
|---|---|---|---|---|---|---|---|---|
| WHRadjBMI | FG | Men | 267 (337) | mmol/L | MR-Egger | -0.008 (-0.102,0.086) | 0.87 | 0.06 |
| WHRadjBMI | FG | Men | 267 (337) | mmol/L | MR-Egger (intercept) | 0.001 (-0.001,0.002) | 0.39 | 0.06 |
| WHRadjBMI | FG | Men | 267 (337) | mmol/L | Weighted median | 0.024 (-0.036,0.084) | 0.43 | 0.45 |
| WHRadjBMI | FG | Women | 267 (337) | mmol/L | IVW | 0.048 (0.023,0.072) | $1.50\times10^{-04}$ | 0.46 |
| WHRadjBMI | FG | Women | 267 (337) | mmol/L | MR-Egger | 0.096 (0.040,0.153) | $7.98\times10^{-04}$ | 0.06 |
| WHRadjBMI | FG | Women | 267 (337) | mmol/L | MR-Egger (intercept) | -0.001 (-0.002,0.000) | 0.06 | 0.06 |
| WHRadjBMI | FG | Women | 267 (337) | mmol/L | Weighted median | 0.052 (0.015,0.088) | 0.006 | 0.45 |
| WHRadjBMI | FI | Combined | 267 (337) | ln(pmol/L) | IVW | 0.137 (0.104,0.170) | $2.23\times10^{-16}$ | - |
| WHRadjBMI | FI | Combined | 267 (337) | ln(pmol/L) | MR-Egger | 0.187 (0.096,0.278) | $5.91\times10^{-05}$ | - |
| WHRadjBMI | FI | Combined | 267 (337) | ln(pmol/L) | MR-Egger (intercept) | -0.001 (-0.002,0.001) | 0.26 | - |
| WHRadjBMI | FI | Combined | 267 (337) | ln(pmol/L) | Weighted median | 0.141 (0.099,0.184) | $7.98\times10^{-11}$ | - |
| WHRadjBMI | FI | Men | 267 (337) | ln(pmol/L) | IVW | 0.093 (0.041,0.144) | $4.32\times10^{-04}$ | 0.13 |
| WHRadjBMI | FI | Men | 267 (337) | ln(pmol/L) | MR-Egger | 0.037 (-0.069,0.143) | 0.49 | 0.005 |
| WHRadjBMI | FI | Men | 267 (337) | ln(pmol/L) | MR-Egger (intercept) | 0.001 (-0.001,0.002) | 0.24 | 0.01 |
| WHRadjBMI | FI | Men | 267 (337) | ln(pmol/L) | Weighted median | 0.078 (0.006,0.150) | 0.03 | 0.08 |
| WHRadjBMI | FI | Women | 267 (337) | ln(pmol/L) | IVW | 0.140 (0.108,0.173) | $2.57\times10^{-17}$ | 0.13 |
| WHRadjBMI | FI | Women | 267 (337) | ln(pmol/L) | MR-Egger | 0.223 (0.149,0.296) | $3.22\times10^{-09}$ | 0.005 |
| WHRadjBMI | FI | Women | 267 (337) | ln(pmol/L) | MR-Egger (intercept) | -0.002 (-0.004,-0.000) | 0.02 | 0.01 |
| WHRadjBMI | FI | Women | 267 (337) | ln(pmol/L) | Weighted median | 0.155 (0.107,0.203) | $2.42\times10^{-10}$ | 0.08 |

BMI, body mass index; CI, confidence interval; FG, fasting glucose; FI, fasting insulin; GRS, genetic risk score; IVW, inverse-variance weighted method; MR, Mendelian randomization; P, P-value; SD, standard deviation; SNP, single nucleotide polymorphism; WHR, waist-hip-ratio; WHRadjBMI, waist-hip-ratio adjusted for body mass index. The method considered the main method, IVW, in bold.

[a]N SNPs used is the number of SNPs for which summary statistics for FI and FG could be obtained

[b]Results in plasma mmol/L levels, untransformed, for FG and serum pmol/L levels, ln-transformed, for FI per 1-SD higher obesity trait, using the sex-specific estimates approach to construct genetic risk scores

[c]P-value threshold set at <0.003 (=0.05/15) for 15 obesity trait-risk factor combinations (including FG, FI, systolic and diastolic blood pressure and smoking status)

[d]$P_{het}$ values from Cochran's Q test given for comparisons between male and female estimates for matching traits and methods. $P_{het}$-threshold set at <0.003 (=0.05/15) for 15 obesity trait-risk factor combinations (including FG, FI, systolic and diastolic blood pressure and smoking status)

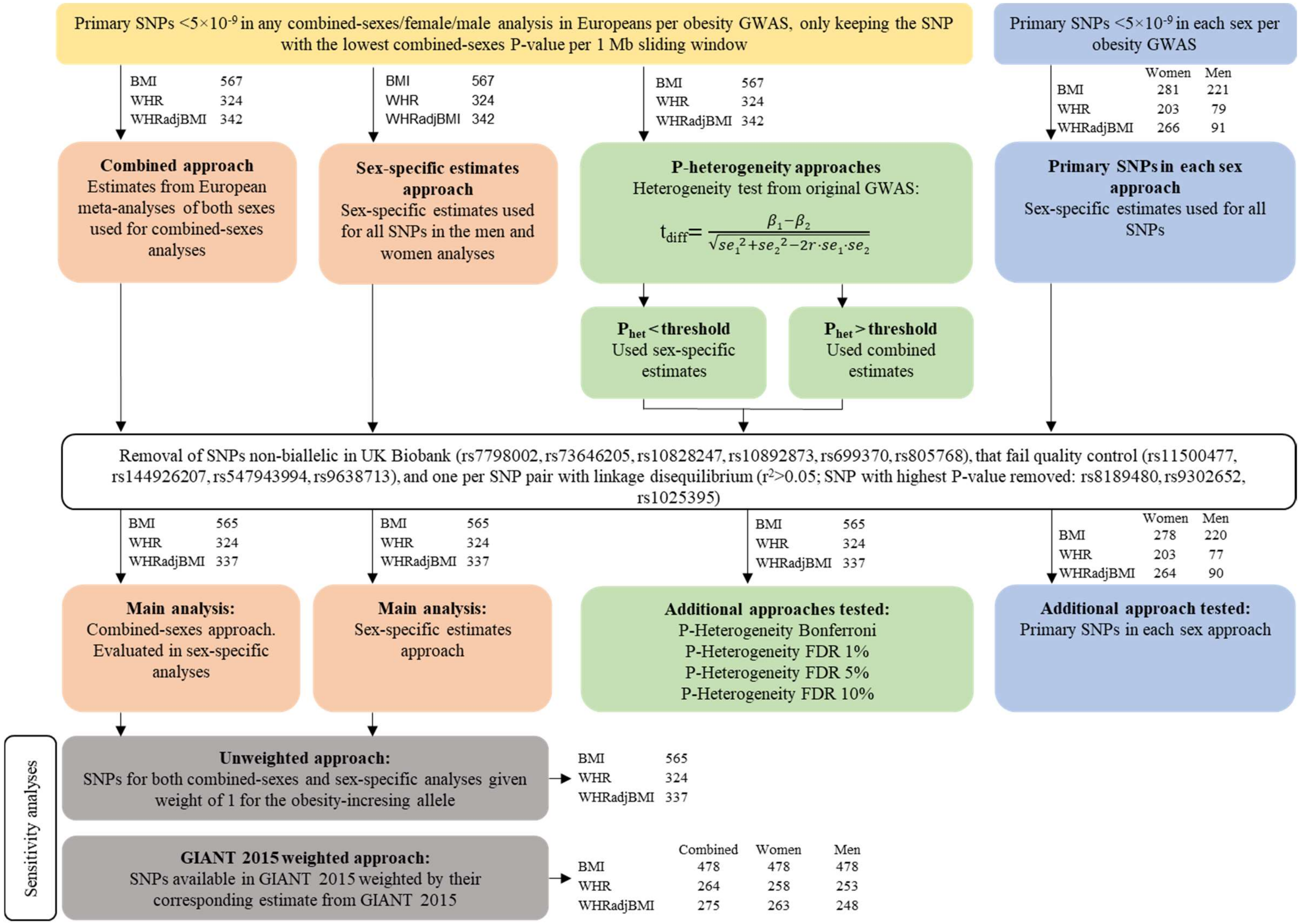


**Figure A. SNP- and weight selection flowchart for all GRS construction approaches.**
Number of SNPs for each trait given after the obesity trait. SNPs were selected by including the primary ("index") variants for each associated (with SNPs $P<5\times10^{-9}$) locus (assessed for a minimum of ±5 Mb of the top SNP and including all SNPs in linkage disequilibrium $R^2>0.05$ and $P<0.05$, and with primary variants as determined through

joint and conditional testing using GCTA in the original study (6)), in any of the men, women, and combined-sexes genome-wide association studies for each obesity trait (6). For the P-heterogeneity approaches, threshold was set to 0.05/N of SNPs for the P-heterogeneity Bonferroni approach, and according to the % FDR for the P-heterogeneity FDR approaches using the $P_{het}$-value ("psexdiff") from the original GWAS (6). BMI, body mass index; FDR, false discovery rate; N, number; SNP, single nucleotide polymorphism; WHR, waist-hip-ratio; WHRadjBMI, waist-hip-ratio adjusted for body mass index.

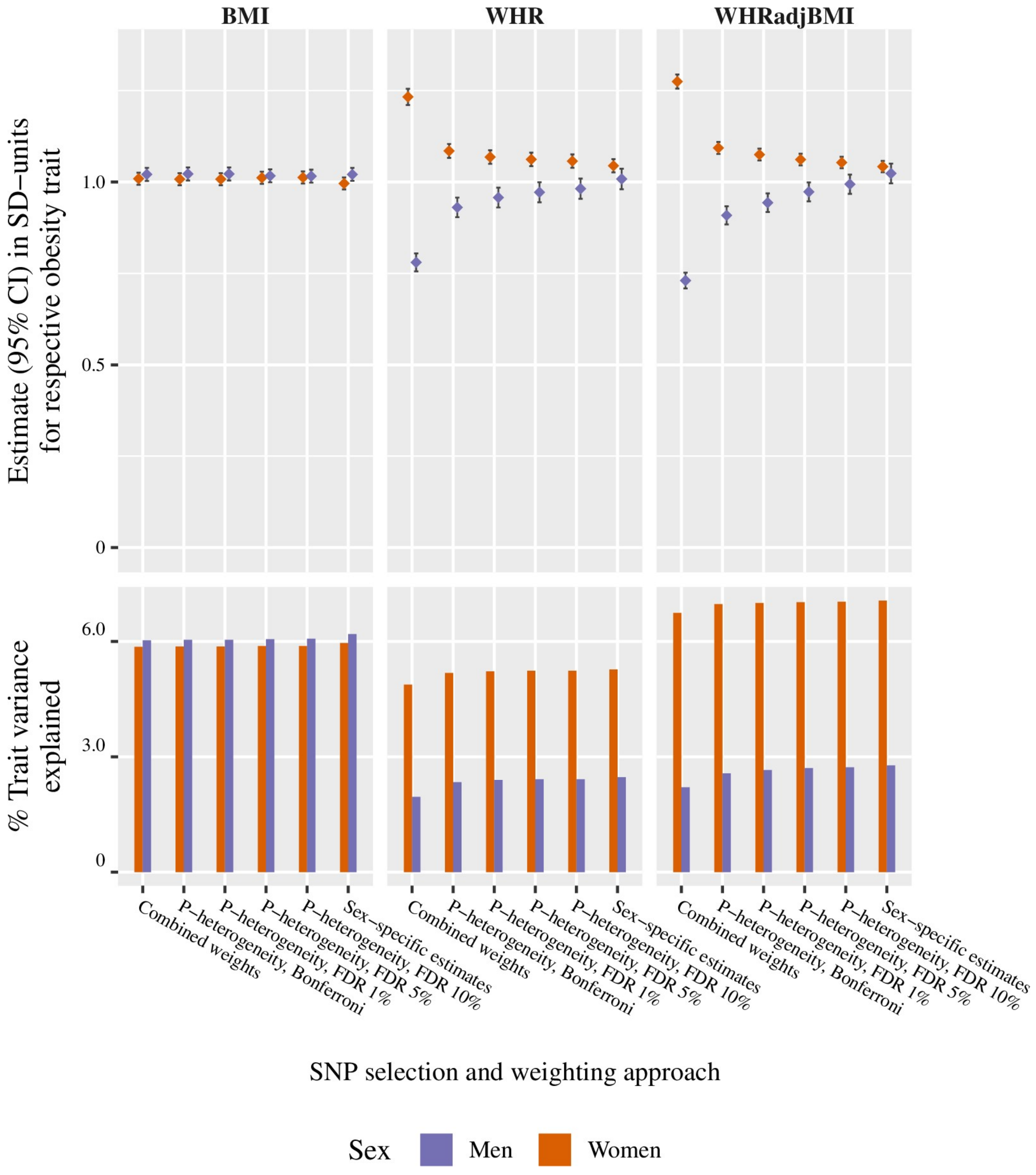


**Figure B. Estimates and trait variance explained for the different sex-specific SNP weighting approaches using the same set of SNPs, separated by trait and sex.**

For the combined weights, P-heterogeneity approaches, and sex-specific estimates the same set of independent SNPs with P<5x10-9 in each trait were included. For the combined weights approach, all SNPs were weighted by the combined-sexes estimate. For the sex-specific estimates approach, all SNPs were weighted by their sex-specific estimate. For the P-heterogeneity approaches, SNPs were weighted by either the sex-specific estimate or the combined-sexes estimates, depending on if the individual SNP had a $P_{het}$-value below the specified threshold. The $P_{het}$ threshold was set to 0.05/N of SNPs for the P-heterogeneity Bonferroni approach and according to the % FDR for the P-heterogeneity FDR approaches. There is a gradual increase in trait variance explained, $R^2$, the more sex-specific weights are used, and with male and female estimates corresponding to a more similar change in the obesity trait, especially for the waist-related traits. BMI, body mass index; CI, confidence interval; FDR, false discovery rate; N, number; Phet, heterogeneity P-value; SNP, single nucleotide polymorphism; WHR, waist-hip-ratio; WHRadjBMI, waist-hip-ratio adjusted for body mass index.

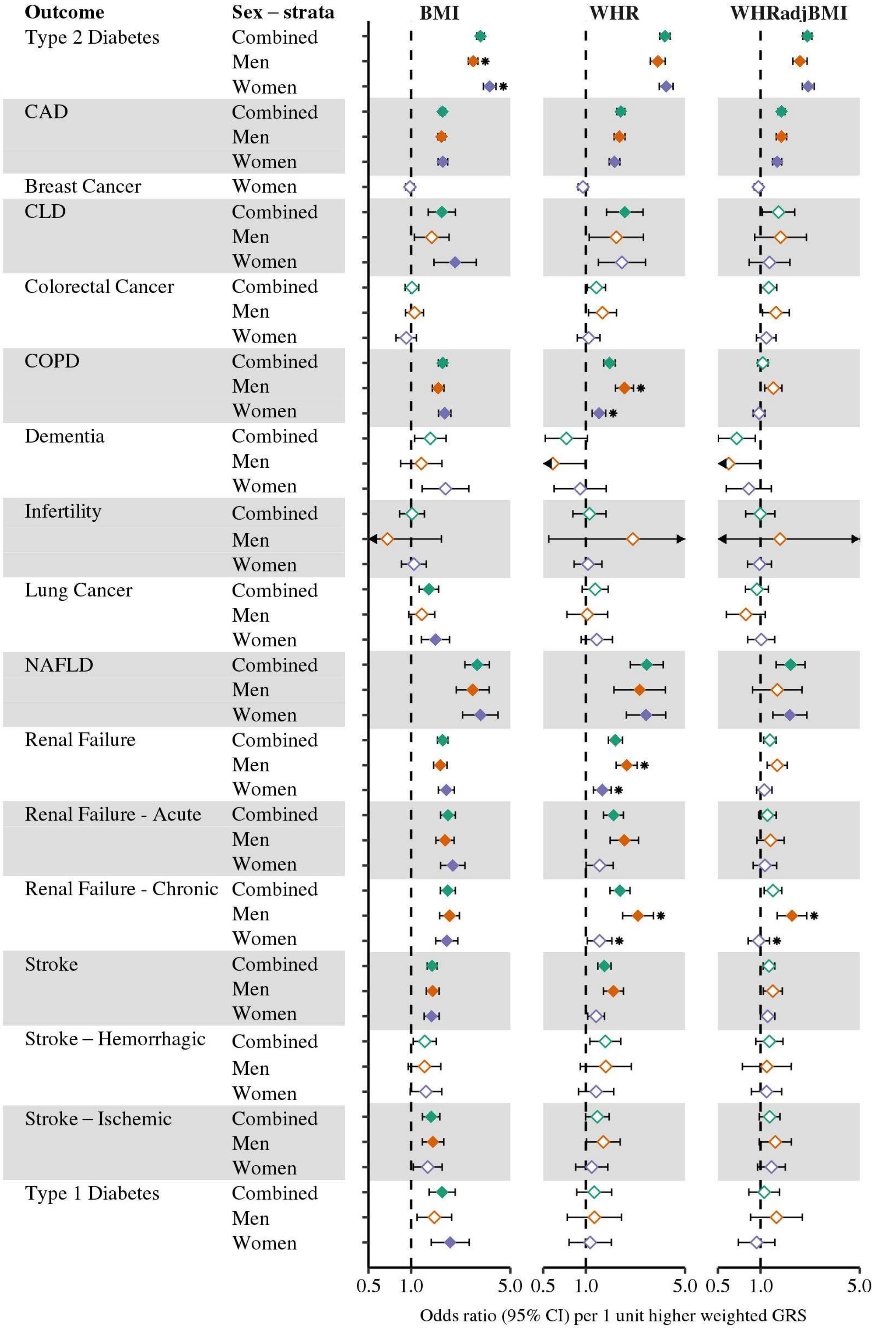


**Figure C. Genetic risk scores association with disease outcomes, stratified by sex.**
Estimates given in odds ratio (95% CI) per 1-unit higher weighted GRS, for exact figures see corresponding Table F. Filled diamonds indicate that the P-value for the obesity trait to disease endpoint surpasses our

threshold for multiple testing; empty diamonds indicate that the P-value does not surpass this threshold (Bonferroni-adjusted P-value-threshold set at <0.001 (=0.05/51) for 51 obesity trait-disease outcome combinations in the study). * denotes that the P-value for heterogeneity (from Cochran's Q test) surpasses our threshold for multiple testing; $P_{het}$-threshold set at <0.001 (=0.05/48) for 48 male-female comparisons in the study (fewer since breast cancer analyses were performed in women only). ♦, combined-sexes estimates; ♦, male estimates; ♦, female estimates; BMI, body mass index; CAD, coronary artery disease; CLD, chronic liver disease; COPD, chronic obstructive pulmonary disease; GRS, genetic risk score; NAFLD, non-alcoholic fatty liver disease; WHR, waist-hip-ratio; WHRadjBMI, waist-hip-ratio adjusted for body mass index.

| Outcome | Risk factor | Sex | OR (95% CI) | P | $P_{het}$ |
|---|---|---|---|---|---|
| Type 2 diabetes | BMI | Combined | 3.03 (2.82,3.27) | $7.2 \times 10^{-192}$ | |
| | | Men | 2.70 (2.46,2.97) | $6.5 \times 10^{-95}$ | $1.3 \times 10^{-4}$ * |
| | | Women | 3.66 (3.24,4.14) | $3.9 \times 10^{-94}$ | |
| | WHR | Combined | 3.70 (3.35,4.08) | $4.1 \times 10^{-147}$ | |
| | | Men | 3.62 (3.08,4.25) | $6.1 \times 10^{-55}$ | 0.51 |
| | | Women | 3.89 (3.38,4.47) | $7.6 \times 10^{-82}$ | |
| | WHRadjBMI | Combined | 2.14 (1.97,2.32) | $1.4 \times 10^{-71}$ | |
| | | Men | 1.97 (1.69,2.30) | $5.4 \times 10^{-18}$ | 0.52 |
| | | Women | 2.10 (1.87,2.36) | $2.2 \times 10^{-35}$ | |
| CAD | BMI | Combined | 1.58 (1.49,1.66) | $3.2 \times 10^{-65}$ | |
| | | Men | 1.55 (1.45,1.65) | $7.9 \times 10^{-39}$ | 0.68 |
| | | Women | 1.58 (1.45,1.73) | $3.3 \times 10^{-23}$ | |
| | WHR | Combined | 1.76 (1.64,1.89) | $1.7 \times 10^{-56}$ | |
| | | Men | 1.80 (1.61,2.01) | $1.1 \times 10^{-24}$ | 0.03 |
| | | Women | 1.52 (1.38,1.69) | $7.0 \times 10^{-16}$ | |
| | WHRadjBMI | Combined | 1.35 (1.27,1.43) | $2.4 \times 10^{-22}$ | |
| | | Men | 1.38 (1.23,1.54) | $2.2 \times 10^{-8}$ | 0.25 |
| | | Women | 1.27 (1.16,1.39) | $1.3 \times 10^{-7}$ | |
| CLD | BMI | Combined | 1.58 (1.24,2.02) | $2.4 \times 10^{-4}$ | |
| | | Men | 1.27 (0.92,1.75) | 0.14 | 0.07 |
| | | Women | 2.04 (1.37,3.04) | $4.4 \times 10^{-4}$ | |
| | WHR | Combined | 1.66 (1.20,2.31) | $2.5 \times 10^{-3}$ | |
| COPD | BMI | Combined | 1.55 (1.43,1.67) | $2.4 \times 10^{-28}$ | |
| | | Men | 1.47 (1.32,1.63) | $3.6 \times 10^{-12}$ | 0.26 |
| | | Women | 1.61 (1.43,1.80) | $1.3 \times 10^{-15}$ | |
| | WHR | Combined | 1.46 (1.31,1.62) | $1.5 \times 10^{-12}$ | |
| | | Men | 1.86 (1.55,2.24) | $3.8 \times 10^{-11}$ | $3.8 \times 10^{-4}$ * |
| | | Women | 1.24 (1.09,1.41) | $1.3 \times 10^{-3}$ | |
| Lung cancer | BMI | Combined | 1.22 (1.02,1.45) | 0.03 | |
| | | Men | 1.05 (0.83,1.33) | 0.69 | 0.12 |
| | | Women | 1.39 (1.07,1.81) | 0.01 | |
| NAFLD | BMI | Combined | 2.77 (2.22,3.45) | $2.4 \times 10^{-19}$ | |
| | | Men | 2.78 (2.05,3.77) | $5.5 \times 10^{-11}$ | 0.87 |
| | | Women | 2.89 (2.07,4.03) | $4.6 \times 10^{-10}$ | |
| | WHR | Combined | 2.35 (1.75,3.17) | $1.6 \times 10^{-8}$ | |
| | | Men | 2.25 (1.34,3.76) | $2.0 \times 10^{-3}$ | 0.85 |
| | | Women | 2.39 (1.64,3.48) | $5.1 \times 10^{-6}$ | |
| | WHRadjBMI | Combined | 1.75 (1.35,2.27) | $2.6 \times 10^{-5}$ | |
| | | Men | 1.24 (0.74,2.08) | 0.42 | 0.24 |
| | | Women | 1.78 (1.29,2.47) | $5.3 \times 10^{-4}$ | |
| Renal failure | BMI | Combined | 1.56 (1.42,1.71) | $4.5 \times 10^{-20}$ | |
| | | Men | 1.48 (1.31,1.68) | $7.5 \times 10^{-10}$ | 0.12 |
| | | Women | 1.72 (1.48,2.01) | $1.6 \times 10^{-12}$ | |
| | WHR | Combined | 1.61 (1.41,1.82) | $2.4 \times 10^{-13}$ | |
| | | Men | 1.70 (1.38,2.10) | $6.5 \times 10^{-7}$ | 0.18 |
| | | Women | 1.41 (1.19,1.68) | $5.7 \times 10^{-5}$ | |
| Renal failure – acute | BMI | Combined | 1.71 (1.49,1.95) | $6.2 \times 10^{-15}$ | |
| | | Men | 1.60 (1.35,1.90) | $4.5 \times 10^{-8}$ | 0.12 |
| | | Women | 2.01 (1.60,2.53) | $2.9 \times 10^{-9}$ | |
| | WHR | Combined | 1.61 (1.35,1.93) | $1.7 \times 10^{-7}$ | |
| | | Men | 1.81 (1.36,2.40) | $4.5 \times 10^{-5}$ | 0.15 |
| | | Women | 1.37 (1.05,1.77) | 0.02 | |
| Renal failure – chronic | BMI | Combined | 1.67 (1.46,1.92) | $1.1 \times 10^{-13}$ | |
| | | Men | 1.66 (1.39,2.00) | $5.0 \times 10^{-8}$ | 0.96 |
| | | Women | 1.68 (1.36,2.07) | $1.3 \times 10^{-6}$ | |
| | WHR | Combined | 1.74 (1.45,2.09) | $2.0 \times 10^{-9}$ | |
| | | Men | 2.08 (1.53,2.83) | $3.4 \times 10^{-6}$ | 0.01 |
| | | Women | 1.28 (1.01,1.62) | 0.04 | |
| | WHRadjBMI | Men | 1.52 (1.12,2.07) | $7.6 \times 10^{-3}$ | 0.03 |
| | | Women | 1.00 (0.81,1.22) | 0.99 | |
| Stroke | BMI | Combined | 1.31 (1.19,1.43) | $4.1 \times 10^{-9}$ | |
| | | Men | 1.37 (1.22,1.54) | $1.6 \times 10^{-7}$ | 0.32 |
| | | Women | 1.25 (1.09,1.44) | $1.9 \times 10^{-3}$ | |
| | WHR | Combined | 1.28 (1.14,1.44) | $4.5 \times 10^{-5}$ | |
| | | Men | 1.38 (1.13,1.68) | $1.8 \times 10^{-3}$ | 0.16 |
| | | Women | 1.15 (0.98,1.34) | 0.09 | |
| Stroke – ischemic | BMI | Combined | 1.31 (1.12,1.53) | $7.8 \times 10^{-4}$ | |
| | | Men | 1.34 (1.10,1.63) | $4.0 \times 10^{-3}$ | 0.57 |
| | | Women | 1.21 (0.93,1.59) | 0.16 | |
| Type 1 diabetes | BMI | Combined | 1.67 (1.31,2.13) | $3.5 \times 10^{-5}$ | |
| | | Men | 1.55 (1.11,2.17) | $9.6 \times 10^{-3}$ | 0.36 |
| | | Women | 1.96 (1.36,2.83) | $3.5 \times 10^{-4}$ | |

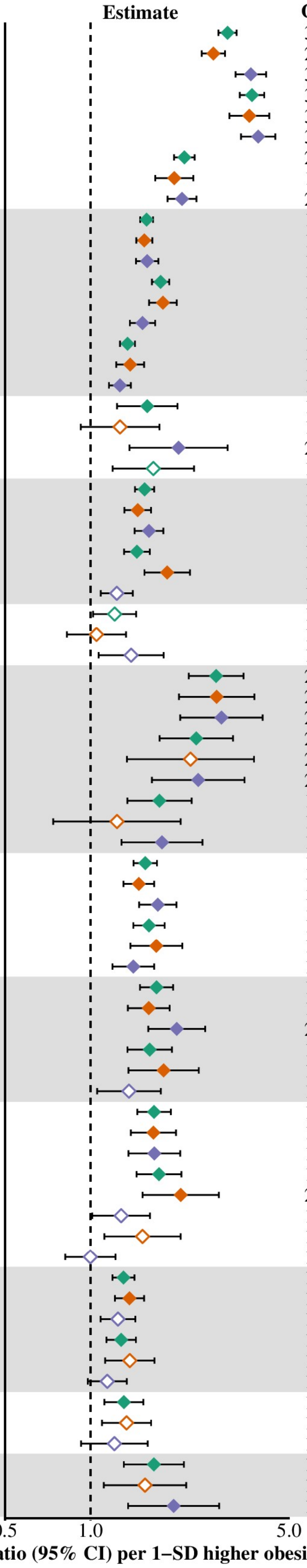


**Figure D. Effect of obesity risk factors on disease outcomes, stratified by sex, using GIANT 2015 sex-specific estimates as weights.**

The obesity trait-disease combinations brought forward for Mendelian randomization, with estimates given in odds ratio (95% CI) per 1-SD higher obesity trait. Filled diamonds indicate that the P-value for the obesity trait to disease endpoint surpasses our threshold for multiple testing; empty diamonds indicate that the P-value does not surpass this threshold (Bonferroni-adjusted P-value-threshold set at <0.001 (=0.05/51) for 51 obesity trait-disease outcome combinations in the study). * denotes that the P-value for heterogeneity (from Cochran's Q test) surpasses our threshold for multiple testing; $P_{het}$-threshold set at <0.001 (=0.05/48) for 48 male-female comparisons in the study (fewer since breast cancer analyses were performed in women only). ♦, combined-sexes estimates; ♦, male estimates; ♦, female estimates; BMI, body mass index; CAD, coronary artery disease; COPD, chronic obstructive pulmonary disease; NAFLD, non-alcoholic fatty liver disease; SD, standard deviation; WHR, waist-hip-ratio; WHRadjBMI, waist-hip-ratio adjusted for body mass index.

| Outcome | Risk factor | Sex | OR (95% CI) | P | $P_{het}$ |
|---|---|---|---|---|---|
| Type 2 diabetes | BMI | Combined | 3.13 (2.92,3.36) | $<1.0 \times 10^{-200}$ | |
| | | Men | 2.93 (2.68,3.21) | $9.6 \times 10^{-122}$ | 0.01 |
| | | Women | 3.52 (3.14,3.96) | $2.3 \times 10^{-101}$ | |
| | WHR | Combined | 3.93 (3.58,4.32) | $4.2 \times 10^{-178}$ | |
| | | Men | 3.64 (3.13,4.23) | $1.5 \times 10^{-63}$ | 0.15 |
| | | Women | 4.22 (3.71,4.80) | $9.9 \times 10^{-105}$ | |
| | WHRadjBMI | Combined | 2.25 (2.07,2.44) | $9.2 \times 10^{-84}$ | |
| | | Men | 2.30 (2.00,2.65) | $6.1 \times 10^{-31}$ | 0.36 |
| | | Women | 2.11 (1.89,2.36) | $2.0 \times 10^{-40}$ | |
| CAD | BMI | Combined | 1.72 (1.63,1.81) | $6.7 \times 10^{-99}$ | |
| | | Men | 1.69 (1.59,1.80) | $2.6 \times 10^{-61}$ | 0.43 |
| | | Women | 1.77 (1.62,1.93) | $4.3 \times 10^{-39}$ | |
| | WHR | Combined | 1.86 (1.74,1.99) | $3.2 \times 10^{-74}$ | |
| | | Men | 1.87 (1.68,2.08) | $9.7 \times 10^{-32}$ | 0.05 |
| | | Women | 1.63 (1.48,1.79) | $1.6 \times 10^{-23}$ | |
| | WHRadjBMI | Combined | 1.37 (1.29,1.46) | $9.3 \times 10^{-26}$ | |
| | | Men | 1.48 (1.33,1.64) | $5.5 \times 10^{-14}$ | 0.02 |
| | | Women | 1.27 (1.17,1.38) | $2.4 \times 10^{-8}$ | |
| CLD | BMI | Combined | 1.76 (1.39,2.23) | $2.3 \times 10^{-6}$ | |
| | | Men | 1.54 (1.13,2.08) | $5.4 \times 10^{-3}$ | 0.15 |
| | | Women | 2.18 (1.50,3.16) | $4.2 \times 10^{-5}$ | |
| | WHR | Combined | 1.82 (1.33,2.49) | $1.6 \times 10^{-4}$ | |
| COPD | BMI | Combined | 1.71 (1.59,1.84) | $7.3 \times 10^{-45}$ | |
| | | Men | 1.68 (1.52,1.87) | $2.0 \times 10^{-23}$ | 0.69 |
| | | Women | 1.74 (1.56,1.93) | $3.3 \times 10^{-23}$ | |
| | WHR | Combined | 1.53 (1.39,1.69) | $2.2 \times 10^{-17}$ | |
| | | Men | 1.79 (1.51,2.12) | $2.1 \times 10^{-11}$ | 0.01 |
| | | Women | 1.38 (1.22,1.56) | $2.0 \times 10^{-7}$ | |
| Lung cancer | BMI | Combined | 1.40 (1.18,1.65) | $8.7 \times 10^{-5}$ | |
| | | Men | 1.28 (1.02,1.61) | 0.03 | 0.27 |
| | | Women | 1.55 (1.21,1.98) | $5.2 \times 10^{-4}$ | |
| NAFLD | BMI | Combined | 3.09 (2.50,3.83) | $2.5 \times 10^{-25}$ | |
| | | Men | 2.66 (1.99,3.56) | $3.2 \times 10^{-11}$ | 0.15 |
| | | Women | 3.65 (2.67,5.00) | $4.8 \times 10^{-16}$ | |
| | WHR | Combined | 2.68 (2.03,3.55) | $6.0 \times 10^{-12}$ | |
| | | Men | 2.42 (1.50,3.89) | $2.9 \times 10^{-4}$ | 0.74 |
| | | Women | 2.67 (1.88,3.79) | $3.5 \times 10^{-8}$ | |
| | WHRadjBMI | Combined | 1.64 (1.27,2.12) | $1.4 \times 10^{-4}$ | |
| | | Men | 1.66 (1.04,2.64) | 0.03 | 0.97 |
| | | Women | 1.67 (1.23,2.28) | $1.1 \times 10^{-3}$ | |
| Renal failure | BMI | Combined | 1.70 (1.55,1.86) | $2.4 \times 10^{-30}$ | |
| | | Men | 1.63 (1.45,1.84) | $4.1 \times 10^{-16}$ | 0.29 |
| | | Women | 1.80 (1.57,2.08) | $3.6 \times 10^{-16}$ | |
| | WHR | Combined | 1.70 (1.51,1.92) | $3.0 \times 10^{-18}$ | |
| | | Men | 2.29 (1.88,2.79) | $1.8 \times 10^{-16}$ | $2.4 \times 10^{-4}$ * |
| | | Women | 1.43 (1.22,1.67) | $9.9 \times 10^{-6}$ | |
| Renal failure – acute | BMI | Combined | 1.82 (1.60,2.07) | $7.0 \times 10^{-20}$ | |
| | | Men | 1.73 (1.48,2.03) | $1.7 \times 10^{-11}$ | 0.31 |
| | | Women | 1.99 (1.60,2.47) | $4.2 \times 10^{-10}$ | |
| | WHR | Combined | 1.67 (1.41,1.98) | $3.7 \times 10^{-9}$ | |
| | | Men | 2.10 (1.61,2.74) | $4.9 \times 10^{-8}$ | 0.04 |
| | | Women | 1.43 (1.13,1.82) | $3.3 \times 10^{-3}$ | |
| Renal failure – chronic | BMI | Combined | 1.89 (1.66,2.15) | $9.1 \times 10^{-22}$ | |
| | | Men | 1.88 (1.58,2.23) | $1.3 \times 10^{-12}$ | 0.87 |
| | | Women | 1.92 (1.57,2.33) | $8.6 \times 10^{-11}$ | |
| | WHR | Combined | 1.85 (1.56,2.20) | $1.9 \times 10^{-12}$ | |
| | | Men | 2.79 (2.09,3.72) | $3.5 \times 10^{-12}$ | $2.4 \times 10^{-4}$ * |
| | | Women | 1.41 (1.14,1.76) | $1.8 \times 10^{-3}$ | |
| | WHRadjBMI | Men | 1.73 (1.31,2.29) | $1.1 \times 10^{-4}$ | $9.2 \times 10^{-4}$ * |
| | | Women | 0.98 (0.81,1.18) | 0.81 | |
| Stroke | BMI | Combined | 1.46 (1.34,1.59) | $5.0 \times 10^{-18}$ | |
| | | Men | 1.45 (1.29,1.62) | $9.5 \times 10^{-11}$ | 0.85 |
| | | Women | 1.47 (1.29,1.68) | $8.8 \times 10^{-9}$ | |
| | WHR | Combined | 1.34 (1.20,1.50) | $2.7 \times 10^{-7}$ | |
| | | Men | 1.50 (1.24,1.81) | $2.0 \times 10^{-5}$ | 0.14 |
| | | Women | 1.25 (1.08,1.45) | $2.7 \times 10^{-3}$ | |
| Stroke – ischemic | BMI | Combined | 1.42 (1.22,1.65) | $5.3 \times 10^{-6}$ | |
| | | Men | 1.44 (1.19,1.74) | $1.4 \times 10^{-4}$ | 0.79 |
| | | Women | 1.38 (1.07,1.78) | 0.01 | |
| Type 1 diabetes | BMI | Combined | 1.55 (1.23,1.96) | $2.2 \times 10^{-4}$ | |
| | | Men | 1.44 (1.05,1.98) | 0.02 | 0.48 |
| | | Women | 1.71 (1.21,2.41) | $2.4 \times 10^{-3}$ | |

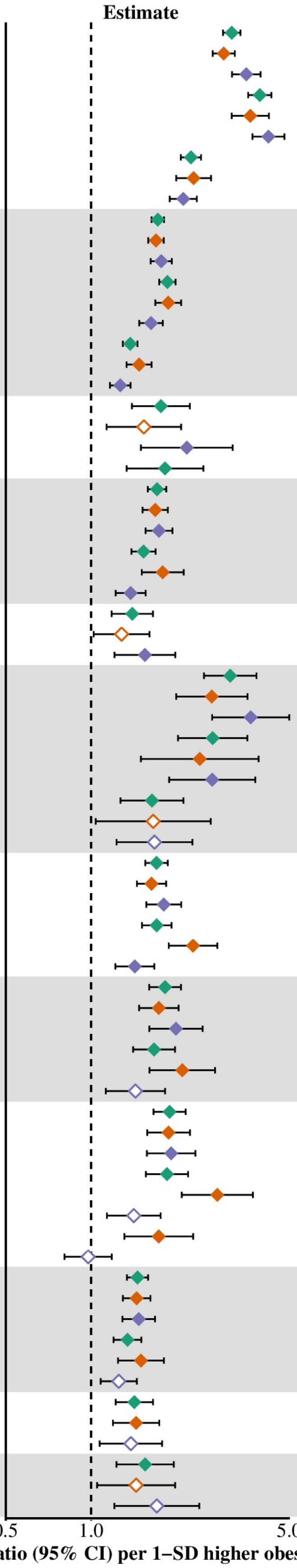


**Figure E. Effect of obesity risk factors on disease outcomes, stratified by sex, using unweighted genetic risk scores.**

The obesity trait-disease combinations brought forward for Mendelian randomization, with estimates given in odds ratio (95% CI) per 1-SD higher obesity trait. Filled diamonds indicate that the P-value for the obesity trait to disease endpoint surpasses our threshold for multiple testing; empty diamonds indicate that the P-value does not surpass this threshold (Bonferroni-adjusted P-value-threshold set at <0.001 (=0.05/51) for 51 obesity trait-disease outcome combinations in the study). * denotes that the P-value for heterogeneity (from Cochran's Q test) surpasses our threshold for multiple testing; $P_{het}$-threshold set at <0.001 (=0.05/48) for 48 male-female comparisons in the study (fewer since breast cancer analyses were performed in women only). ♦, combined-sexes estimates; ♦, male estimates; ♦, female estimates; BMI, body mass index; CAD, coronary artery disease; COPD, chronic obstructive pulmonary disease; NAFLD, non-alcoholic fatty liver disease; SD, standard deviation; WHR, waist-hip-ratio; WHRadjBMI, waist-hip-ratio adjusted for body mass index.

| Outcome | Risk factor | Sex | OR (95% CI) | P | $P_{het}$ |
|---|---|---|---|---|---|
| Type 2 diabetes | BMI | Combined | 3.22 (3.00,3.45) | $<1.0 \times 10^{-200}$ | |
| | | Men | 2.80 (2.57,3.05) | $1.3 \times 10^{-122}$ | $7.1 \times 10^{-6}$ * |
| | | Women | 3.88 (3.46,4.36) | $1.7 \times 10^{-118}$ | |
| | WHR | Combined | 3.66 (3.35,4.01) | $1.9 \times 10^{-171}$ | |
| | | Men | 3.43 (2.99,3.93) | $2.9 \times 10^{-69}$ | 0.52 |
| | | Women | 3.64 (3.23,4.10) | $3.0 \times 10^{-100}$ | |
| | WHRadjBMI | Combined | 2.11 (1.95,2.28) | $8.2 \times 10^{-82}$ | |
| | | Men | 1.92 (1.70,2.17) | $4.3 \times 10^{-25}$ | 0.24 |
| | | Women | 2.11 (1.91,2.33) | $2.1 \times 10^{-49}$ | |
| CAD | BMI | Combined | 1.64 (1.56,1.72) | $4.1 \times 10^{-86}$ | |
| | | Men | 1.61 (1.51,1.71) | $2.7 \times 10^{-55}$ | 0.48 |
| | | Women | 1.67 (1.53,1.81) | $1.8 \times 10^{-32}$ | |
| | WHR | Combined | 1.70 (1.59,1.81) | $1.5 \times 10^{-58}$ | |
| | | Men | 1.71 (1.55,1.88) | $1.2 \times 10^{-28}$ | 0.11 |
| | | Women | 1.54 (1.41,1.68) | $3.1 \times 10^{-21}$ | |
| | WHRadjBMI | Combined | 1.38 (1.31,1.46) | $2.9 \times 10^{-30}$ | |
| | | Men | 1.42 (1.30,1.55) | $1.6 \times 10^{-14}$ | 0.15 |
| | | Women | 1.30 (1.21,1.40) | $8.5 \times 10^{-12}$ | |
| CLD | BMI | Combined | 1.65 (1.30,2.08) | $2.8 \times 10^{-5}$ | |
| | | Men | 1.47 (1.10,1.97) | 0.01 | 0.30 |
| | | Women | 1.88 (1.30,2.73) | $8.5 \times 10^{-4}$ | |
| | WHR | Combined | 1.97 (1.44,2.67) | $1.7 \times 10^{-5}$ | |
| COPD | BMI | Combined | 1.65 (1.53,1.77) | $3.5 \times 10^{-40}$ | |
| | | Men | 1.51 (1.37,1.66) | $2.7 \times 10^{-16}$ | 0.05 |
| | | Women | 1.74 (1.56,1.94) | $1.2 \times 10^{-23}$ | |
| | WHR | Combined | 1.45 (1.32,1.60) | $4.4 \times 10^{-14}$ | |
| | | Men | 1.89 (1.61,2.21) | $2.2 \times 10^{-15}$ | $4.9 \times 10^{-6}$ * |
| | | Women | 1.20 (1.07,1.35) | $1.6 \times 10^{-3}$ | |
| Lung cancer | BMI | Combined | 1.39 (1.17,1.64) | $1.1 \times 10^{-4}$ | |
| | | Men | 1.21 (0.97,1.51) | 0.08 | 0.10 |
| | | Women | 1.60 (1.25,2.05) | $1.8 \times 10^{-4}$ | |
| NAFLD | BMI | Combined | 2.76 (2.24,3.41) | $4.3 \times 10^{-21}$ | |
| | | Men | 2.56 (1.93,3.39) | $5.6 \times 10^{-11}$ | 0.49 |
| | | Women | 2.96 (2.18,4.04) | $5.6 \times 10^{-12}$ | |
| | WHR | Combined | 2.85 (2.16,3.76) | $1.1 \times 10^{-13}$ | |
| | | Men | 2.63 (1.68,4.09) | $2.1 \times 10^{-5}$ | 0.89 |
| | | Women | 2.73 (1.97,3.78) | $1.4 \times 10^{-9}$ | |
| | WHRadjBMI | Combined | 1.71 (1.34,2.17) | $1.4 \times 10^{-5}$ | |
| | | Men | 1.35 (0.89,2.05) | 0.16 | 0.42 |
| | | Women | 1.66 (1.25,2.19) | $3.8 \times 10^{-4}$ | |
| Renal failure | BMI | Combined | 1.69 (1.54,1.85) | $1.4 \times 10^{-30}$ | |
| | | Men | 1.62 (1.45,1.82) | $3.8 \times 10^{-17}$ | 0.27 |
| | | Women | 1.80 (1.56,2.07) | $2.1 \times 10^{-16}$ | |
| | WHR | Combined | 1.61 (1.43,1.81) | $1.3 \times 10^{-15}$ | |
| | | Men | 1.98 (1.65,2.37) | $8.2 \times 10^{-14}$ | $1.9 \times 10^{-4}$ * |
| | | Women | 1.27 (1.10,1.47) | $1.3 \times 10^{-3}$ | |
| Renal failure – acute | BMI | Combined | 1.85 (1.63,2.10) | $2.5 \times 10^{-21}$ | |
| | | Men | 1.74 (1.49,2.03) | $1.6 \times 10^{-12}$ | 0.22 |
| | | Women | 2.05 (1.66,2.54) | $4.6 \times 10^{-11}$ | |
| | WHR | Combined | 1.57 (1.33,1.85) | $1.2 \times 10^{-7}$ | |
| | | Men | 1.86 (1.46,2.37) | $5.9 \times 10^{-7}$ | 0.04 |
| | | Women | 1.30 (1.04,1.63) | 0.02 | |
| Renal failure – chronic | BMI | Combined | 1.83 (1.61,2.08) | $1.8 \times 10^{-20}$ | |
| | | Men | 1.89 (1.60,2.23) | $5.4 \times 10^{-14}$ | 0.67 |
| | | Women | 1.79 (1.47,2.17) | $3.2 \times 10^{-9}$ | |
| | WHR | Combined | 1.73 (1.46,2.04) | $1.1 \times 10^{-10}$ | |
| | | Men | 2.38 (1.83,3.10) | $9.1 \times 10^{-11}$ | $8.3 \times 10^{-5}$ * |
| | | Women | 1.23 (1.00,1.50) | 0.05 | |
| | WHRadjBMI | Men | 1.60 (1.25,2.05) | $1.8 \times 10^{-4}$ | $7.5 \times 10^{-4}$ * |
| | | Women | 0.95 (0.80,1.13) | 0.59 | |
| Stroke | BMI | Combined | 1.41 (1.29,1.53) | $1.8 \times 10^{-15}$ | |
| | | Men | 1.39 (1.25,1.55) | $1.2 \times 10^{-9}$ | 0.75 |
| | | Women | 1.43 (1.26,1.63) | $6.3 \times 10^{-8}$ | |
| | WHR | Combined | 1.32 (1.18,1.47) | $7.4 \times 10^{-7}$ | |
| | | Men | 1.51 (1.27,1.79) | $1.9 \times 10^{-6}$ | 0.03 |
| | | Women | 1.18 (1.03,1.36) | 0.02 | |
| Stroke – ischemic | BMI | Combined | 1.33 (1.14,1.54) | $1.8 \times 10^{-4}$ | |
| | | Men | 1.35 (1.12,1.61) | $1.2 \times 10^{-3}$ | 0.80 |
| | | Women | 1.29 (1.01,1.66) | 0.04 | |
| Type 1 diabetes | BMI | Combined | 1.71 (1.36,2.15) | $5.2 \times 10^{-6}$ | |
| | | Men | 1.51 (1.12,2.05) | $7.0 \times 10^{-3}$ | 0.29 |
| | | Women | 1.94 (1.38,2.73) | $1.5 \times 10^{-4}$ | |

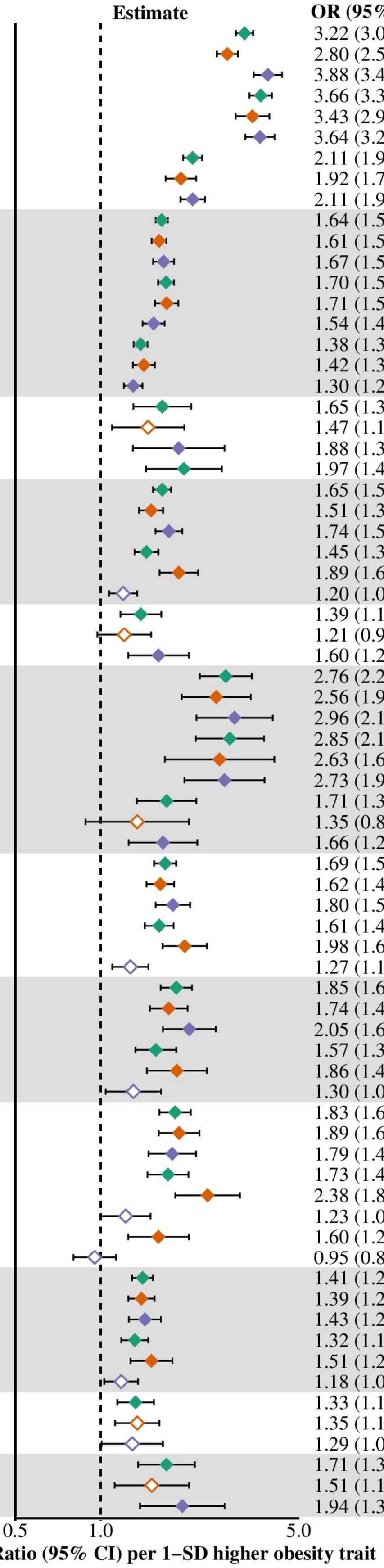


**Figure F. Effect of obesity risk factors on disease outcomes, stratified by sex, performed in the British ancestry only subset.**

Participants were denoted as "British" if they were in the British ancestry subset as defined by the UK Biobank (14) (based on self-report of British ancestry and similar ancestry according to principal components analysis).

The figure shows the obesity trait-disease combinations brought forward for Mendelian randomization, with estimates given in odds ratio (95% CI) per 1-SD higher obesity trait. Filled diamonds indicate that the P-value for the obesity trait to disease endpoint surpasses our threshold for multiple testing; empty diamonds indicate that the P-value does not surpass this threshold (Bonferroni-adjusted P-value-threshold set at <0.001 (=0.05/51) for 51 obesity trait-disease outcome combinations in the study). * denotes that the P-value for heterogeneity (from Cochran's Q test) surpasses our threshold for multiple testing; $P_{het}$-threshold set at <0.001 (=0.05/48) for 48 male-female comparisons in the study (fewer since breast cancer analyses were performed in women only). ♦, combined-sexes estimates; ♦, male estimates; ♦, female estimates; BMI, body mass index; CAD, coronary artery disease; COPD, chronic obstructive pulmonary disease; NAFLD, non-alcoholic fatty liver disease; SD, standard deviation; WHR, waist-hip-ratio; WHRadjBMI, waist-hip-ratio adjusted for body mass index.

27. Burgess S, Scott RA, Timpson NJ, Davey Smith G, Thompson SG, EPIC- InterAct

Consortium. Using published data in Mendelian randomization: a blueprint for efficient identification of causal risk factors. Eur J Epidemiol [Internet]. 2015 Jul 15 [cited 2018 Jul 9];30(7):543–52. Available from: http://link.springer.com/10.1007/s10654-015-0011-z
